# CD276 Immature Glycosylation Drives Colorectal Cancer Aggressiveness and T-cell Mediated Immune Escape

**DOI:** 10.1101/2025.08.01.668091

**Authors:** Janine Soares, Dylan Ferreira, Andreia Miranda, Martina Gonçalves, Marta Relvas-Santos, Andreia Brandão, Paula Paulo, Sofia Cotton, Rui Freitas, Mariana Magalhães, Eduardo Ferreira, Beatriz Marinho-Santos, Luís Pedro Afonso, André M N Silva, Carlos Palmeira, Francisco Amado, Andreia Peixoto, Lúcio Lara Santos, José Alexandre Ferreira

## Abstract

Colorectal cancer (CRC) progression is fueled by immune evasion, yet the underlying molecular mechanisms remain to be fully characterized. CD276 (B7-H3), an immune checkpoint glycoprotein frequently overexpressed in aggressive tumours, is extensively modified by glycosylation, a process known to regulate protein stability, localization, and immune interactions. However, its glycosylation-dependent functions in CRC remain unclear. This study shows that poor-prognosis CRC tumours exhibit dysregulated glycogene expression, leading to an immature *O*-glycosylation phenotype that enhances malignancy. Using *C1GALT1* knockout CRC cells, which recapitulate the cancer glycocalyx, it was demonstrated that this glycosylation shift promotes proliferation and invasion. We further identify CD276 as a metastasis-associated glycoprotein whose function is shaped by *O*-glycan modifications. Mechanistically, immature *O*-glycosylation of CD276 enhances invasion, suppresses T-cell activation, and induces an immunosuppressive cytokine milieu, reinforcing its role in tumour immune escape. These findings establish CD276 as a glycosylation-dependent immune checkpoint and a promising therapeutic target to overcome immune evasion in CRC.

**Graphical Abstract:** 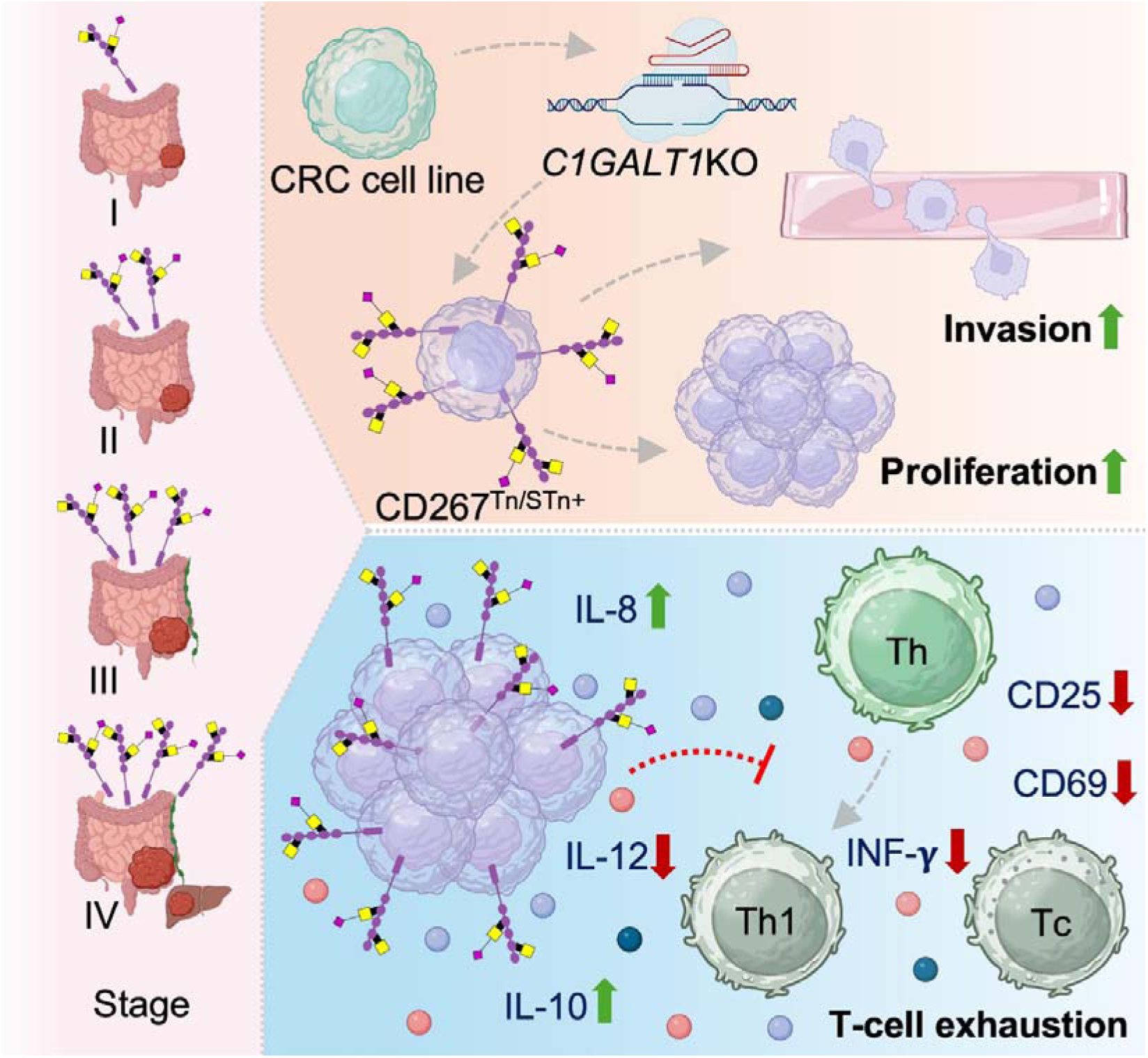

**Table of Contents:** Colorectal cancer (CRC) progression is closely linked to immune evasion, yet the molecular mechanisms underlying this process remain poorly understood. This study identifies CD276 (B7-H3) as a glycosylation-driven regulator of CRC aggressiveness and demonstrates how *O*-glycosylation remodeling promotes tumour immune escape. These findings establish CD276 as a potential therapeutic target and highlight the role of glycoproteoform-specific immune modulation in cancer progression.

## Introduction

Colorectal cancer (CRC) remains one of the most lethal malignancies worldwide, with poor survival rates for patients with metastatic disease [1]. While immune checkpoint inhibitors (ICIs) have transformed cancer therapy, their efficacy is largely restricted to tumours with microsatellite instability (MSI) and high tumour mutational burden (TMB) [2]. Most CRC cases, particularly those with microsatellite stability (MSS), overall representing about 95% of the metastatic CRCs, remain resistant [3], highlighting a critical gap in understanding alternative immune evasion mechanisms that drive tumour progression.

Among the emerging immune modulators in cancer, CD276 (B7-H3) has gained attention as a non-canonical immune checkpoint molecule with a pivotal role in tumour progression. CD276 is a type I transmembrane glycoprotein belonging to the B7 family of co-stimulatory and co-inhibitory immune regulators, yet its physiological immune function remains controversial [4]. Unlike classical immune checkpoints such as PD-L1 and CTLA-4, CD276 lacks a well-defined receptor and can exert both immunosuppressive and pro-tumorigenic functions depending on the tumour context [4b, 5]. Nonetheless, CD276 overexpression has been reported across multiple malignancies, including lung [6], breast [7], prostate [8], and colorectal cancers [9], where it correlates with poor prognosis, increased metastatic potential, and therapy resistance. Beyond immune regulation, CD276 has been implicated in epithelial-mesenchymal transition (EMT) [10], resistance to apoptosis [11], and angiogenesis [12]. Still the molecular mechanisms governing its function remain incompletely understood, particularly at the post-translational level.

Glycosylation plays a crucial role in regulating immune checkpoint function and response to immunotherapy, including in CD276 [13]. Recent evidence indicates that core fucose *N*-glycosylation, mediated by FUT8, stabilizes CD276, modulates its surface expression, and suppresses T-cell activation and proliferation in triple-negative breast cancer [13a]. Inhibiting core fucosylation has emerged as a promising strategy to enhance anti-tumour immune responses, underscoring the significance of glycosylation as a therapeutic target. However, the role of *O*-glycosylation, an essential post-translational modification in immune regulation and tumour progression [14], remains poorly defined in CD276 function. In CRC, extensive remodeling of the cancer glycocalyx leads to an immature *O*-glycosylation phenotype, primarily due to Cosmc (encoded by *C1GALT1C1*) dysregulation [15]. Cosmc is essential for the function of Core 1 β1,3-galactosyltransferase 1 (encoded by *C1GALT1*), the key enzyme responsible for *O*-glycan elongation via Core 1 synthesis. Loss of Cosmc activity results in the accumulation of the Tn antigen, a hallmark of aberrant *O*-glycosylation [16]. These alterations are predominantly driven by *C1GALT1C1* promoter hypermethylation and, in some cases, loss-of-function mutations [17]. This dysregulation has been strongly linked to cancer invasion mainly by disrupting normal *O*-glycosylation of cell surface proteins, leading to impaired cell-cell adhesion, enhanced migration, and metastasis [18], while also promoting immune evasion via macrophage-binding lectins on innate immune cells [19]. While the role of C1GALT1-specific chaperone 1 in CRC is well established, the direct contribution of Core 1 β1,3-galactosyltransferase 1 itself in tumour progression remains poorly understood [15a]. More broadly, despite growing evidence of *O*-glycosylation’s impact on cancer biology and immune modulation, how it shapes CD276 function, particularly in the context of immune interactions and tumour progression, remains largely unknown.

In this study, leveraging a systematic analysis of the cancer glycome, we identify CD276 as a metastasis-associated glycoprotein extensively modified by *O*-glycosylation in CRC. We demonstrate that tumours with poor prognostic features exhibit aberrant glycogene expression, leading to an immature *O*-glycosylation phenotype that enhances malignancy. Using *C1GALT1* knockout CRC cells, which recapitulate the cancer-associated glycocalyx, we reveal that *O*-glycan remodeling amplifies CD276-driven tumour invasion and impairs T-cell activation. These findings establish CD276 as a glycosylation-dependent immune checkpoint and highlight its potential as a therapeutic target to overcome immune evasion in CRC.

## Results

In this study, we started by investigating the role of immature protein *O*-glycosylation in aggressive CRC. We first analyzed glycogene expression patterns in patient samples and then profiled glycan composition in CRC tissues across differentiation states. Finally, we focused on CD276, a metastasis-associated immune checkpoint glycoprotein, to examine how *O*-glycan remodeling influences tumour cell behavior and immune suppression.

### Transcriptomic shifts in *O*-glycosylation pathways define CRC aggressiveness

We began by addressing the expression of key glycogenes encoding glycosyltransferases involved in the initial steps of protein *O*-glycosylation. Analysis of TCGA dataset, comprising over 600 CRC cases across all disease stages and normal mucosa samples, revealed significant alterations in glycosylation pathways associated with tumour progression and aggressiveness. Most notably, *B3GNT6*, which initiates core 3 *O*-glycan biosynthesis, was markedly downregulated in CRC (**Figure 1A**), suggesting a reduction in core 3-type *O*-glycans in malignant tissues. In parallel, *ST6GALNAC1-4*, which mediate *O*-6 GalNAc sialylation, were consistently decreased in cancer tissues compared to healthy mucosa. In contrast, there was significant upregulation of *C1GALT1* and its chaperone *C1GALT1C1*, which extend glycans beyond the Tn antigen into core 1 structures. *ST3GAL1,* a sialyltransferase promoting sialyl-T antigen formation, was also upregulated. These findings indicate a biosynthetic shift favoring core 1 over core 3 *O*-glycans in CRC (as highlighted by the schematic *O*-glycosylation pathway representation in **Figure S1**, Supporting Information). Importantly, low *B3GNT6* expression was strongly associated with poor clinical outcomes, with patients with *B3GNT6*^Low^ tumours showing significantly worse overall survival (*p* = 0.0002; HR = 2.32, *p* < 0.001; **Figure 1B**).

**Figure 1.**
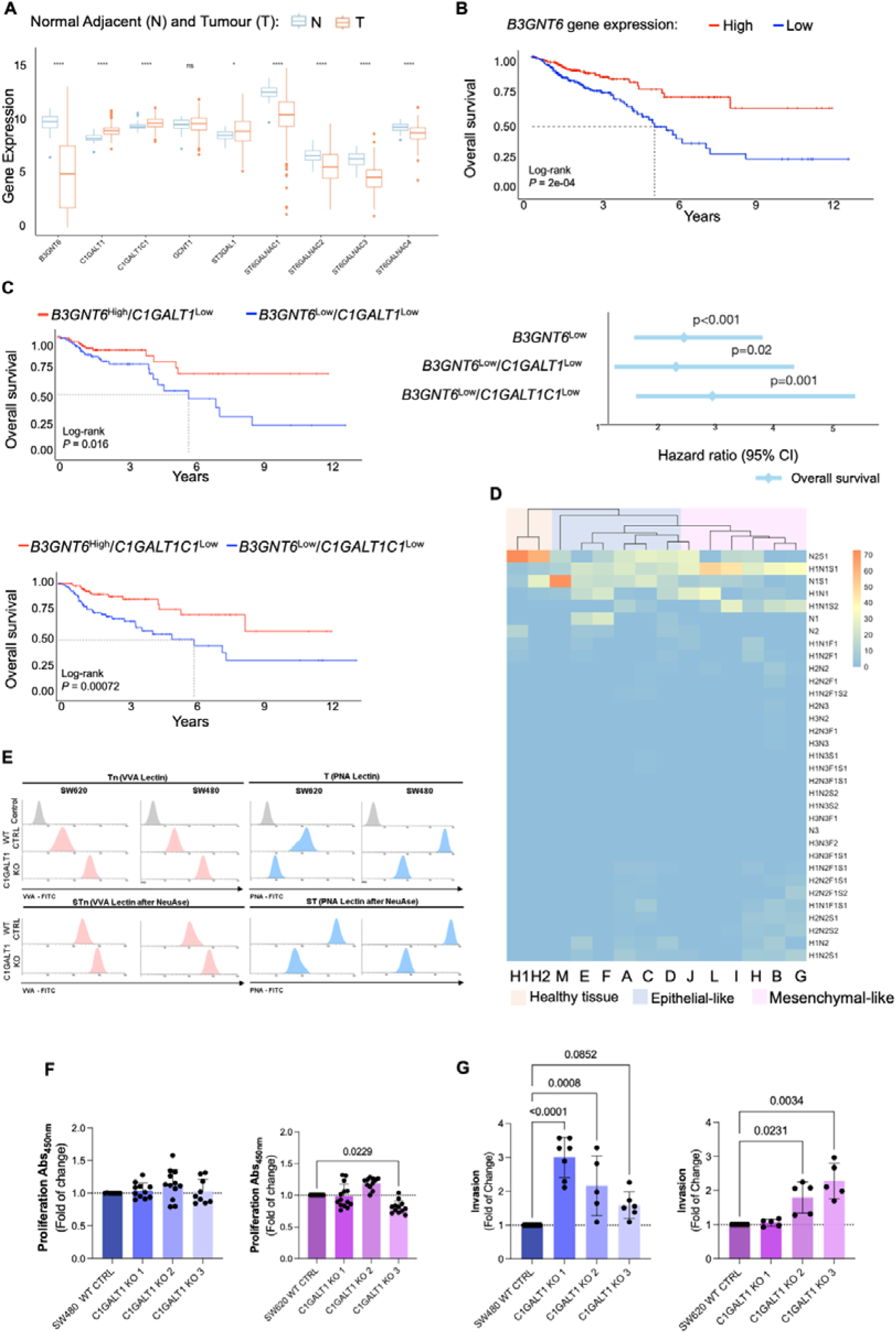
**Downregulation of *B3GNT6* and *C1GALT1* promotes immature *O*-glycosylation, defines poor-prognosis CRC subtypes, and enhances tumour cell aggressiveness. A. Key *O*-glycosylation enzymes are downregulated in colorectal tumours**. Transcriptomic analysis of the TCGA-COADREAD dataset comprising 427 samples (376 tumour and 51 normal adjacent tissues) revealed significant downregulation of *B3GNT6* in tumour samples compared to normal tissue (Wilcoxon test, *p* < 0.05), indicating impairment in core 3 *O*-glycan synthesis. **B. *B3GNT6* expression is associated with patient prognosis.** Kaplan–Meier survival analysis showed that lower *B3GNT6* expression is significantly associated with decreased overall survival in CRC patients (log-rank *p* < 0.05). **C. Combined downregulation of *B3GNT6* and *C1GALT1* or *C1GALT1C1* identifies high-risk subgroups.** Patients with low *B3GNT6* and either low *C1GALT1* or *C1GALT1C1* expression (associated with an immature glycophenotype) had significantly worse survival (log-rank *p* < 0.05), reinforcing the prognostic relevance of immature *O*-glycosylation profiles. **D. Glycomic profiling distinguishes tumour differentiation states, metastatic potential, and deviation from normal glycosylation.** Mass spectrometry-based glycomic analysis revealed that healthy colon mucosa (H1, H2) predominantly expresses sialylated core 3 *O*-glycans. In contrast, epithelial-like tumours are enriched in Tn/sTn antigens, while mesenchymal-like tumours present mono- or di-sialylated core 1 structures. Metastatic tumours exhibit elongated core 2 *O*-glycans irrespective of differentiation state, illustrating the progressive shift from mature to truncated *O*-glycan profiles across CRC progression. **E. *C1GALT1* knockout induces immature *O*-glycans in CRC cells, mimicking tumour-associated glycophenotypes.** Flow cytometry analysis of SW480 and SW620 *C1GALT1* KO cells (n=3 independent replicates per condition) revealed reduced PNA binding, both with and without neuraminidase treatment, indicating loss of core 1 and sialylated core 1 glycans, consistent with impaired *O*-glycan elongation. Concurrently, increased VVA binding was observed, revealing accumulation of Tn antigens. After neuraminidase treatment, a further increase in VVA signal confirmed the presence of sTn, demonstrating that many Tn structures are capped by sialic acids. Together, these results validate the induction of a truncated and sialylated *O*-glycophenotype in *C1GALT1*-deficient cells, which closely mimics glycosylation features of colorectal tumours. **F. Immature glycosylation does not affect cell proliferation.** *C1GALT1* KO does not significantly change proliferation compared to wild-type controls (n = 15 wells per condition, from 3 independent experiments; one-way ANOVA with Tukey post-hoc; *p* < 0.05). **G. Immature glycosylation increases tumour cell invasiveness.** Matrigel invasion assays increase in invasiveness in both SW480 as well as two clones of SW620 *C1GALT1* KO cells compared to wild type (n = 8 wells per condition, from 3 independent experiments. Differences were statistically significant (*p* < 0.05; one-way ANOVA with Tukey post-hoc test).

To further investigate glycophenotypic patterns, we stratified tumours based on *B3GNT6* expression in combination with low levels of *C1GALT1* or *C1GALT1C1 (***Figure 1C**). Although *C1GALT1* and *C1GALT1C1* are upregulated in CRC (**Figure 1A**), their reduced expression, when coupled with low *B3GNT6*, may limit core 1 and core 3 synthesis, leading to immature, cancer-associated glycosylation. Accordingly, tumours with a *B3GNT6*^Low^/*C1GALT1*^Low^ or *B3GNT6*^Low^/*C1GALT1C1*^Low^ profile, likely reflecting an immature glycosylation state, were associated with the poorest prognosis (*B3GNT6*^Low^/*C1GALT1*^Low^: HR = 2.18, *p* < 0.02; *B3GNT6*^Low^/*C1GALT1C1*^Low^: HR = 2.79, *p* < 0.001; **Figure 1C**). Together, these findings suggest that CRC progression is linked to simple glycophenotypes, marked by the accumulation of Tn and potentially sTn antigens, which correlate with more aggressive disease and poorer clinical outcomes.

### Differentiation-linked glycome profiles reveal simple glycophenotypes in advanced CRC

To explore whether these transcriptional alterations translate into distinct glycan phenotypes, we next profiled the tissue glycome in healthy mucosa and advanced CRC (stage ≥3), focusing on how tumour differentiation shapes *O*-glycosylation patterns. The tumours were further characterized based on their differentiation status, a feature often linked to consensus molecular subtypes (CMS) [20]. Accordingly, we categorized tumours into two groups: i) Differentiated epithelial tumours, characterized by high CDX2 and low FRMD6 and HTR2B expression levels (**Figure S2a**), typically corresponding to CMS2 and CMS3 subtypes; and ii) Undifferentiated mesenchymal tumours, defined by low CDX2 and high FRMD6 and HTR2B levels, generally associated with CMS1 or CMS4 [21].

No differences in survival were observed between these groups (**Figure S2a**), likely reflecting the uniformly advanced stage of disease in the cohort. In healthy mucosa, the glycome was primarily composed of core 3 structures (N2, N2S1; **Figure 1D**, **Figure S3, Table S1 and Table S2**), predominantly decorated with *O-6-GalNAc-*linked sialic acids (**Figure S3**). The sTn antigen (N1S1) was also present at lower levels, consistent with previous reports [22]. These features align with the higher expression of *B3GNT6* and *ST6GALNAC* sialyltransferase genes in healthy colon tissue (**Figure 1A**). By contrast, tumour samples showed significantly lower levels of core 3 glycans and were predominantly enriched in sialyl-T antigens (H1N1S1) and di-sialyl-T antigens (H1N1S2; **Figure 1D**). Most tumours also expressed Tn (N1) and/or sTn (N1S1), while a smaller subset, particularly among mesenchymal-like tumours, also exhibited more extended core 2 *O*-glycans, including H2N2, H2N2F1, H1N2F1S2, H1N2S1, H2N2S2, H2N2S1, H1N2F1, and H2N2F1S2. Nevertheless, a striking difference emerged between epithelial-like and mesenchymal-like tumours. Epithelial-like tumours exhibited higher levels of core 3 and sTn and *de novo* expression of Tn antigens, consistent with a shift toward simpler glycophenotypes (**Figure 1D, Table S2**). In contrast, mesenchymal-like tumours were enriched in mono- and di-sialylated T antigens, with low relative expression of other glycan structures. Remarkably, all tumours with distant metastases, regardless of differentiation status, exhibited multiple sialylated core 2-related structures (H1N2S1, H2N2S2, H2N2S1, H2N2F1S2; **Figure 1D**). No additional associations were observed between tumour stage, grade, patient survival, and glycome composition (data not shown). Complementary immunohistochemistry confirmed the presence of Tn and/or sTn antigens in 90% of primary tumours (**Table S3**), in agreement with MS analysis. Interestingly, while the relative abundance of these glycans varied between epithelial- and mesenchymal-like tumours (**Figure 1D)**, immunohistochemistry analysis showed that overall expression levels of Tn and sTn remained consistent across differentiation states (data not shown). Notably, the expression of sTn showed an association with the T stage (**Table S3**).

In summary, these findings highlight distinct glycome profiles between tumour differentiation states in aggressive colorectal cancer. Immature glycosylation patterns were prevalent across advanced tumours, including lymph node and distant metastases. Epithelial-like lesions displayed an intermediate glycophenotype, situated between healthy colonic mucosa and mesenchymal-like tumours. This profile was marked by a partial loss of core 3 structures, reduced glycan complexity, and increased expression of Tn and sTn antigens. In contrast, mesenchymal tumours were more enriched in sialylated core 1 and extended core 2 glycans. Despite these subtype-specific differences, simple glycophenotypes, particularly Tn and sTn, were widely present across tumour types, with their relative abundance shaped by differentiation status. These findings align with transcriptomics data, confirming that downregulation of core 3–associated enzyme in CRC (**Figure 1A**). They further support the presence of simple glycophenotypes driven also by reduced expression of *C1GALT1* and *C1GALT1C1*, which vary with tumour differentiation and are associated with disease progression.

### Simple glycophenotypes enhance CRC cell invasiveness

To explore the functional role of simple glycophenotypes, we first profiled the *O*-glycome of a panel of CRC cell lines representative of epithelial-like (HCA7, SW480) and mesenchymal-like (SW620, HCT116, RKO, LS174T) phenotypes (**Figure S4 and Table S4**). These profiles closely mirrored human tumours, with epithelial-like cells predominantly expressing core 3 glycans and sialylated T antigens, while mesenchymal-like lines were mainly enriched in core 1 structures. Notably, sTn antigens was absent in epithelial-like cells, suggesting their expression in tumours may be influenced by the tumour microenvironment.

To experimentally induce simple glycophenotypes, we knocked out *C1GALT1* using CRISPR-Cas9 in both SW480 (epithelial-like, derived from a primary tumour) and SW620 (mesenchymal-like, from a lymph node metastasis of the same patient) cell lines (**Figure S5**). Despite their differing baseline glycomes, *C1GALT1* knockout completely suppressed core 1 structures in both cell lines and led to robust overexpression of Tn and sTn antigens (**Figure 1E, Figure S6**). Additionally, both cell lines presented a marked decrease in core 3 glycans. These changes were accompanied by reduced expression of *B3GNT6* (**Figure S5C**), suggesting a degree of co-regulation between core 1 and core 3 glycosyltransferases, in mimicry of the *C1GALT1*^Low^/*B3GNT6*^Low^ phenotype associated with poor prognosis in patient samples (**Figure 1B-C**). Both SW480 and SW620 C1GALT1 knockout cells showed no change in proliferation compared to controls (**Figure 1F**) but exhibited increased matrigel invasion *in vitro* (**Figure 1G**). Collectively, these findings demonstrate that acquisition of simple glycophenotypes, marked by the loss of core 1 and core 3 glycans and accumulation of Tn/sTn antigens, enhances the invasive potential of CRC cells independently of proliferation. This glycome remodeling mirrors the poor-prognosis transcriptional signatures identified in patient tumours and highlights a functional role for immature *O*-glycans in promoting CRC aggressiveness.

### CD276 carries immature *O*-glycans in CRC and associates with poor prognosis

We next investigated whether CD276 could act as a carrier of immature *O*-glycans in CRC. Using immunohistochemistry, we confirmed CD276 expression in approximately 80% of primary advanced CRC tumours, as well as in corresponding lymph node and distant metastases. Notably, CD276 co-localized with areas of high Tn and/or sTn expression in all CD276-positive tumours (**Figure 2A**). In healthy colon tissue, although some apparent co-localization was observed, CD276 was primarily expressed in enterocytes, whereas Tn and sTn antigens were restricted to goblet cells and were much less abundant than in tumours (**Figure 2A**). To further assess whether CD276 directly carried these immature glycans, proximity ligation assay (PLA) and immunofluorescence analyses were performed. Both approaches provided positive signals in CRC samples, supporting the notion that CD276 is a major carrier of Tn and sTn antigens in cancer (**Figure 2B-C**). In contrast, no evidence of CD276 carrying abnormal glycosylation was found in healthy colon tissue or in other normal tissues, including stomach, liver, pancreas, appendix, gallbladder, lung, testis and thyroid (**Figure 2D**), suggesting a cancer-specific nature.

**Figure 2.**
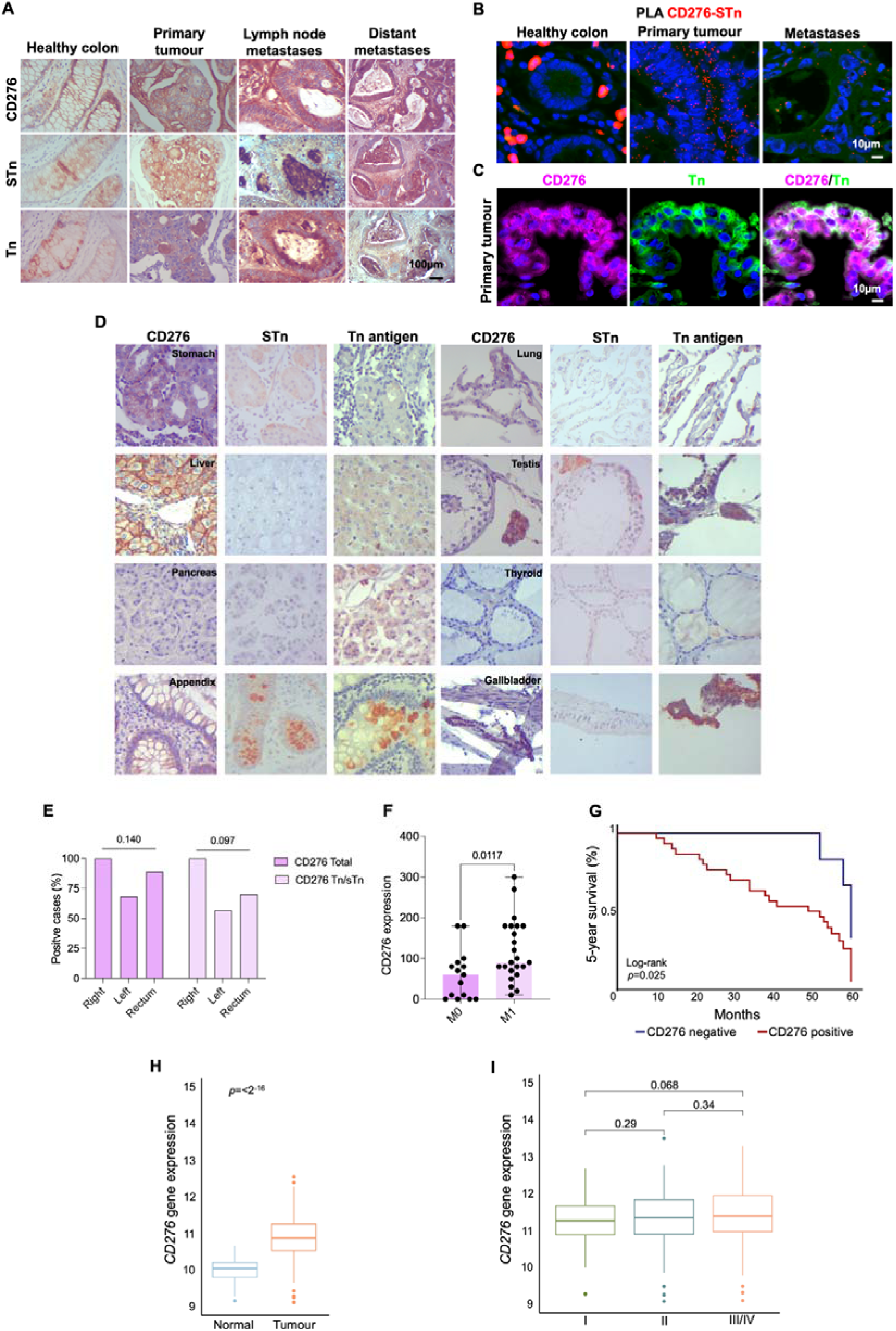
**CD276 is a marker of poor prognosis in colorectal cancer and displays cancer-specific aberrant *O*-glycosylation. A. CD276 colocalizes with Tn and sTn antigens in CRC and metastases.** Immunohistochemistry reveals diffuse CD276 expression throughout tumour regions, both in the cytoplasm and at the cell membrane, overlapping with Tn and sTn antigen distribution. Unlike sTn, CD276 is also detected in the extracellular matrix surrounding tumour areas. Tn expression appears in scattered niches without a consistent pattern. In contrast, the healthy colon shows only weak cytoplasmic staining for CD276, Tn, and sTn, with no evidence of colocalization. **B. *In situ* proximity ligation assay (PLA) confirms CD276-sTn glycoproteoforms in CRC and metastases.** PLA reveals strong spatial proximity (< 30□nm; red puncta) between CD276 and sTn in tumour cells from both primary tumours and metastases, but not in healthy colon, highlighting the cancer-specific nature of this aberrant glycoform. **C. Double immunofluorescence reveals CD276-Tn glycoproteoforms in CRC and metastases.** In CD276-enriched tumours, CD276 (magenta) and Tn (green) co-localize at both the membrane and cytoplasm of the same cells, confirming CD276 as a direct carrier of immature O-glycans. **D. Aberrant CD276 glycoforms are cancer-specific.** Immunohistochemistry analysis of healthy tissues reveals limited CD276 expression, primarily cytoplasmic in select epithelial cells (e.g. enterocytes, hepatocytes, Leydig cells), and no co-localization with Tn or sTn antigens except occasional overlap in Leydig cells. In contrast to tumours, sTn and Tn were either absent or restricted to secretory or immune compartments, supporting the tumour-specific nature of CD276-Tn/sTn glycoforms. **E. CD276 and its aberrantly glycosylated forms are enriched in right-sided colorectal tumours** CD276 and CD276-Tn/sTn glycoform prevalence across tumour locations revealed anatomical distinctions in both expression and glycosylation. In the right colon, 100% of tumours were CD276-positive and all exhibited aberrant glycosylation with Tn and/or sTn antigens. These expressions were trendily lower for the left side colon and rectum. **F. CD276 expression is elevated in metastatic CRC.** IHC quantification shows significantly higher CD276 scores in M1 (metastatic) versus M0 (non-metastatic) tumours (*p*□=□0.0117, unpaired t-test). **F. CD276 expression is associated with poor prognosis.** Kaplan–Meier survival analysis reveals significantly worse 5-year survival in patients with CD276-positive tumours compared to CD276-negative cases (*p*□*=*□0.025, log-rank test). H. *CD276* is significantly upregulated in CRC compared to matched normal tissue. TCGA analysis of tumour samples and adjacent normal tissues shows robust overexpression of CD276 in tumours (*p*□<□0.0001, Wilcoxon test), confirming its cancer-associated expression profile. I. *CD276* expression increases with tumour stage. TCGA data supports a progressive increase of CD276 expression from stage I to stage III/IV (*p*□=□0.068, one-way ANOVA).

Clinically, CD276 carrying immature glycosylation was more frequently observed in right-sided colon tumors compared to left-sided and rectal lesions (**Figure 2E**), a subgroup associated with more aggressive clinical behavior, as well as in metastatic tumors (*p* = 0.0117; **Figure 2F**). Furthermore, CD276-positive tumours were associated with significantly worse prognosis compared to CD276-negative lesions (*p* = 0.0025; **Figure 2G**). Interestingly, no differences were observed in the frequency of CD276 expression or in the prevalence of its abnormally glycosylated forms between epithelial-like and mesenchymal-like tumours (data not shown).

Having established CD276 as a potential cancer-specific carrier of immature *O*-glycans, we next explored its gene expression profile. TCGA analysis revealed that CD276 was significantly upregulated in CRC compared to adjacent normal mucosa (**Figure 2H**). Moreover, CD276 expression showed a trend toward higher levels in more advanced tumour stages (*p* = 0.07 for stage III/IV; **Figure 2I**), reinforcing the link between CD276 and poor prognosis.

### CD276 exhibits distinct glycocodes across colorectal cancer differentiation states

To further dissect the glycocode of CD276, we interrogated a well-characterized proteomics dataset from the PRIDE repository, comprising 95 CRC patients, stratified by tumour stage, location, and differentiation status according to CMS. Protein identifications were derived from individual search result files for each sample **(Supporting Data File 1**). Although the dataset was not specifically optimized for glycoprotein detection, it provided valuable insights into CD276 glycosylation patterns across CRC subtypes. Over 80% of these tumours were positive for CD276, with widespread expression across all stages of the disease (**Figure 3A**). Most tumours in stages III and IV exhibited CD276 positivity (**Figure 3A**). Notably, the proportion of abnormally glycosylated CD276 (Tn/sTn glycoforms) was significantly higher in stage II–IV tumours compared to stage I (40–60% vs. 5%; **Figure 3A**). When stratified by tumour differentiation states, mesenchymal-like tumours (CMS1/4) exhibited significantly higher CD276 protein expression compared to epithelial-like tumours (CMS2/3) (*p* < 0.0001; **Figure 3B**). However, no significant differences were observed in the expression of abnormally glycosylated CD276 forms between epithelial- and mesenchymal-like subtypes (**Figure 3C, Figure S7 and Supporting Data File 2**). Furthermore, tandem mass spectrometry revealed distinct CD276 glycosylation patterns associated with tumour differentiation, independent of anatomical location or stage (**Figure 3D**). While site-specific assignments remain tentative due to the labile nature of O-glycans under CID fragmentation, multiple HexNAc-retaining peptide fragments were detected. These support the presence of immature O-glycosylation in CD276, in agreement with immunoassay data, and align with subtype-specific profiles. Notably, in epithelial-like tumours, glycosylation was more abundant and clustered along the immunoglobulin variable (IgV) and constant (IgC) domains of CD276 (**Figure 3E**). In contrast, mesenchymal-like tumours exhibited sparser and more membrane-proximal glycosylation. Interestingly, O-glycosylation site prediction using NetOGlyc identified only three putative sites, including T244, which was also supported by our analysis. However, MS-based profiling revealed a substantially broader and phenotype-dependent glycosylation landscape, underscoring the limitations of prediction tools and the added value of experimental mapping (**Figure 3E**). Altogether, the results indicate that CD276 glycosylation is primarily linked to tumour differentiation, rather than anatomical location or metastatic status. This provides a foundation for understanding its functional role in cancer and for guiding therapeutic targeting strategies.

**Figure 3.**
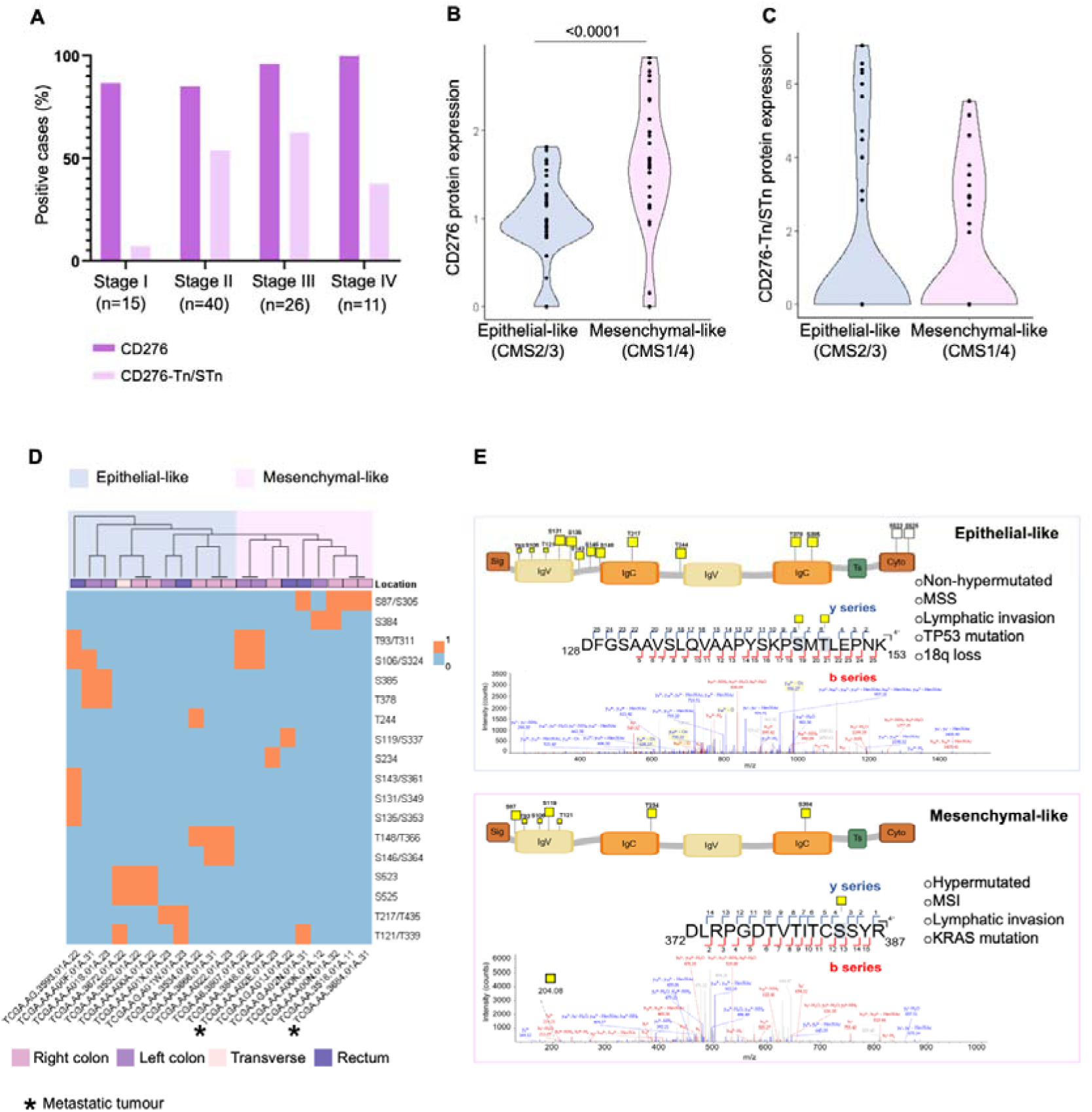
**CD276 is widely expressed across CRC stages and displays distinct glycosylation patterns according to tumour differentiation. A. CD276 is broadly expressed across colorectal cancer stages, while immature CD276 glycoforms (Tn/sTn-modified) are enriched in advanced tumors.** The plot concerning TCGA-based proteomics data shows the percentage of positive cases for total CD276 and CD276 glycosylated with Tn/sTn antigens across CRC stages I–IV. Aberrant CD276 glycoforms were significantly more frequent in stages II–IV compared to stage I (Fisher’s exact test, *p*□=□0.0026). **B. CD276 expression is associated with tumour differentiation state.** Mesenchymal-like tumours (CMS1/4) exhibited significantly higher CD276 protein expression than epithelial-like tumours (CMS2/3) (*p*□<□0.0001, Wilcoxon test). **C. Glycosylated CD276 is not significantly associated with tumour subtype.** No difference in CD276-Tn/sTn glycoform expression was observed between epithelial-like and mesenchymal-like tumours. **D. CD276 glycosylation patterns differ between epithelial-like and mesenchymal-like colorectal tumours.** Heatmap showing the distribution of inferred O-glycosylation sites across tumour samples, stratified by transcriptomic subtype (CMS2/3: epithelial-like; CMS1/4: mesenchymal-like). Each row represents an amino acid position, and each column a tumour; coloured squares indicate detected glycosylation. Epithelial-like tumours display broader glycosylation coverage, with sites clustered within the IgV and IgC domains, whereas mesenchymal-like tumours exhibit sparser and more membrane-proximal modifications. **E. Representative glycopeptides identified by CID MS/MS highlight subtype-specific O-glycosylation within the extracellular domain of CD276.** In epithelial-like tumours (top), glycosylation was denser and predominantly located between the IgV and IgC domains. In contrast, mesenchymal-like tumours (bottom) showed sparser and more membrane-proximal glycosylation. Annotated MS/MS spectra display HexNAc-retaining b- and y-ions supporting glycopeptide identification despite CID-related limitations in site resolution. Schematic domain maps indicate glycosite positions, and subtype-associated molecular features are also summarized.

### CD276 overexpression is driven by immature glycosylation

We next investigated CD276 glycoproteoforms in both wild-type and *C1GALT1* knockout CRC cell models. Immunoblotting using an antibody directed against the membrane portion of CD276 revealed an increase in the 100 kDa band following *C1GALT1* knockout in both SW480 and SW620 cells (**Figure 4A**). A similar pattern was observed using an antibody targeting the cytoplasmic domain (**Figure 4B**). The 100 kDa band is consistent with the full-length CD276 isoform containing two IgV–IgC domain pairs, with the increased apparent molecular weight reflecting extensive glycosylation. These findings indicate that *C1GALT1* knockouts consistently lead to the accumulation of a densely glycosylated CD276 form, independent of the antibody epitope targeted. Additional lower molecular weight bands were detected with the cytoplasmic domain targeting antibody (**Figure 4B**), potentially representing alternative isoforms; however, these bands did not show consistent glycosylation patterns. To further validate the presence of immature *O*-glycans, CD276 was immunoprecipitated from wild-type and *C1GALT1* knockout cells. Isotype control immunoprecipitations showed no detectable bands, confirming the specificity of CD276 pulldown. In a validation experiment using SW620 cells, the predominant 100 kDa CD276 band was enriched and exhibited strong reactivity with the VVA lectin, confirming the accumulation of Tn antigen on CD276 following disruption of glycosylation extension (**Figure 4C and Figure S8**). To evaluate whether *C1GALT1* knockout affects CD276 stability, cells were treated with cycloheximide to block protein synthesis, and CD276 levels were tracked over time. In wild-type cells, CD276 levels declined as expected (**Figure 4D**). In contrast, knockout cells showed delayed degradation, indicating increased protein stability. This effect was statistically significant in SW620 cells (*p* < 0.05) and showed a similar, though non-significant, trend in SW480 cells (**Figure 4D**). These results suggest that loss of core 1 O-glycosylation impairs CD276 turnover, enhancing its post-translational stability.

**Figure 4.**
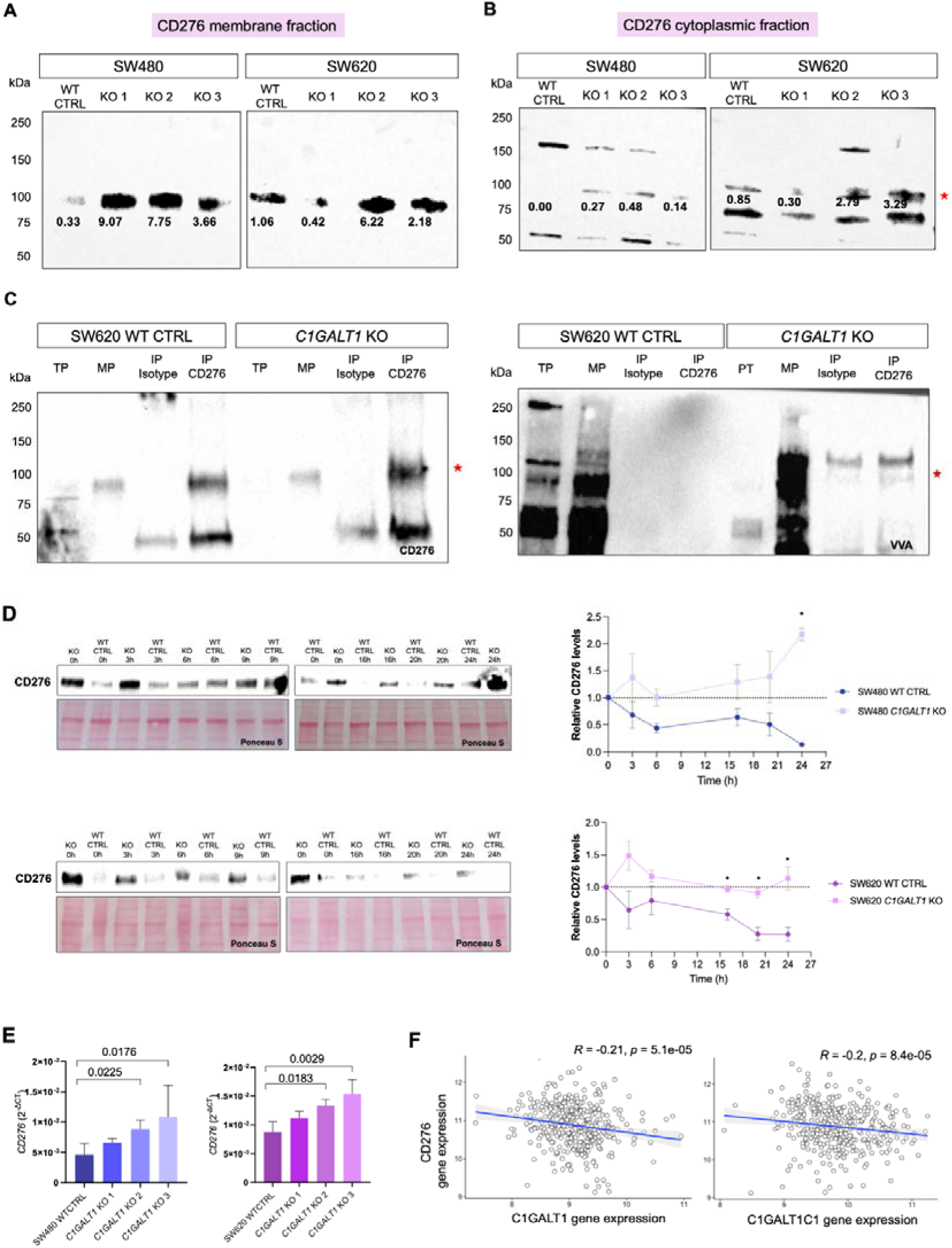
**Loss of *C1GALT1* expression increases CD276 expression and results in accumulation of immature Tn-modified glycoforms. A–B) Loss of *C1GALT1* leads to an increase in CD276 protein levels in CRC cells.** Western blot analysis was performed on total protein lysates (T) and plasma membrane fractions (PM) from SW480 and SW620 wild-type (WT) and *C1GALT1* knockout (KO) cells using antibodies against either the extracellular (**A**) or cytoplasmic (**B**) domains of CD276. Bands were normalized to Ponceau S and are highlighted in bold in the blots. KO cells showed increased CD276 protein expression compared to WT. A prominent 100□kDa band (red asterisk), compatible a CD276 glycoproteoform, was observed in *C1GALT1* KO cells. **C. Immunoprecipitation of CD276 confirms increased protein levels and enrichment of a 100**□kDa glycoform in *C1GALT1* KO cells. CD276 was immunoprecipitated from plasma membrane fractions of SW620 WT and KO cells. Western blotting detected a clear ∼100□kDa band in *C1GALT1* KO samples, which was absent in isotype controls and enhanced relative to WT. This result confirms successful IP and increased expression of CD276 glycoproteoforms in KO cells. VVA lectin blotting was also performed to assess Tn antigen on CD276, confirming the presence of immature glycoforms in CD236 IPs. Ponceau S staining was used to verify equal protein loading. D. Cycloheximide treatment reveals enhanced CD276 protein stability in *C1GALT1* knockout cells. SW480 and SW620 wild-type and C1GALT1 KOs were treated with cycloheximide to block protein synthesis, and CD276 levels were analyzed by immunoblotting overtime. Quantification shows significantly increased CD276 stability in SW620 KO cells compared to WT (*p* < 0.05), and a similar non-significant trend in SW480 cells, indicating that loss of O-glycan elongation stabilizes CD276 post-translationally. E. Loss of *C1GALT1* leads to transcriptional upregulation of *CD276*. In both cell lines, CD276 gene expression was significantly elevated in *C1GALT1* KO clones compared to WT controls. Each bar represents a biologically independent C1GALT1 KO clone, and results are shown as the mean of technical triplicates. Statistical significance was determined using one-way ANOVA followed by Tukey’s multiple comparison test. F. Reduced expression of core 1 *O*-glycosylation enzymes is associated with increased CD276 expression in CRC. TCGA transcriptomic analysis revealed that *CD276* gene expression inversely correlates with that of *C1GALT1* (R□=□–0.21, *p*□=□5.1□×□10^⁻^□) and *C1GALT1C1* (R□=□-0.20, *p*□=□8.4□×□10^⁻5^). These results support a negative association between core 1 *O*-glycan biosynthesis and CD276 expression in CRC, consistent with increased CD276 levels and immature glycosylation under reduced *C1GALT1* expression.

Finally, we addressed CD276 transcript levels, which were significantly increased in *C1GALT1* knockout cells compared to wild-type controls in both cell lines (**Figure 4E**). Analysis of patient TCGA datasets further revealed a significant inverse correlation between *CD276* expression and both *C1GALT1* and *C1GALT1C1* expression (**Figure 4F**), reinforcing the possible transcriptional co-regulation mechanisms linking immature *O*-glycosylation to CD276 upregulation suggested by glycoengineered cell lines. Similar observations were made for *GCNT1*, involved in core 2 formation, reinforcing a link with immature glycosylation. In addition, CD276 associated with increased expression of *ST3GAL1*, *ST6GALNAC2*, and *ST6GALNAC3*, suggesting a shift toward enhanced sialylation of immature *O*-glycans, particularly Tn and core 1 structures (**Figure S8**). These observations point to broader, previously unrecognized regulatory networks linking *O*-glycosylation remodeling and CD276 expression in CRC, warranting further investigation.

Collectively, these findings demonstrate that immature *O*-glycosylation driven by *C1GALT1* loss promotes CD276 overexpression through combined effects on protein stability and gene transcription, highlighting a mechanistic link between glycome remodeling and CD276 regulation in CRC.

### CD276 immature glycosylation drives cancer cell aggressiveness and oncogenic signaling

We used SW620 cells, a metastatic model with mesenchymal traits, to explore how aberrant CD276 glycosylation drives tumour aggressiveness in the context of advanced disease. *CD276* was transiently knocked down using two siRNAs (si1 and si2) in SW480 and SW620 wild-type cells, as well as in their corresponding *C1GALT1* knockout models (**Figure 5A**). Effects on proliferation and invasion were then assessed (**Figure 5B-C**). Efficient knockdown was confirmed by a strong reduction in *CD276* mRNA levels and a near-complete loss of CD276 protein within 24 hours, which was sustained for at least 5 days (**Figure 5A and B**). CD276 knockdown significantly reduced proliferation in *C1GALT1-*deficient cells but had no effect in wild-type cells, indicating that immature O-glycosylation enhances its pro-proliferative function. Silencing CD276 also decreased invasion across both backgrounds, confirming a baseline pro-invasive role. Notably, in *C1GALT1*-deficient cells, CD276 knockdown fully reversed the glycosylation-induced increase in invasion, demonstrating that aberrant glycosylation specifically drives CD276-dependent invasiveness.

**Figure 5.**
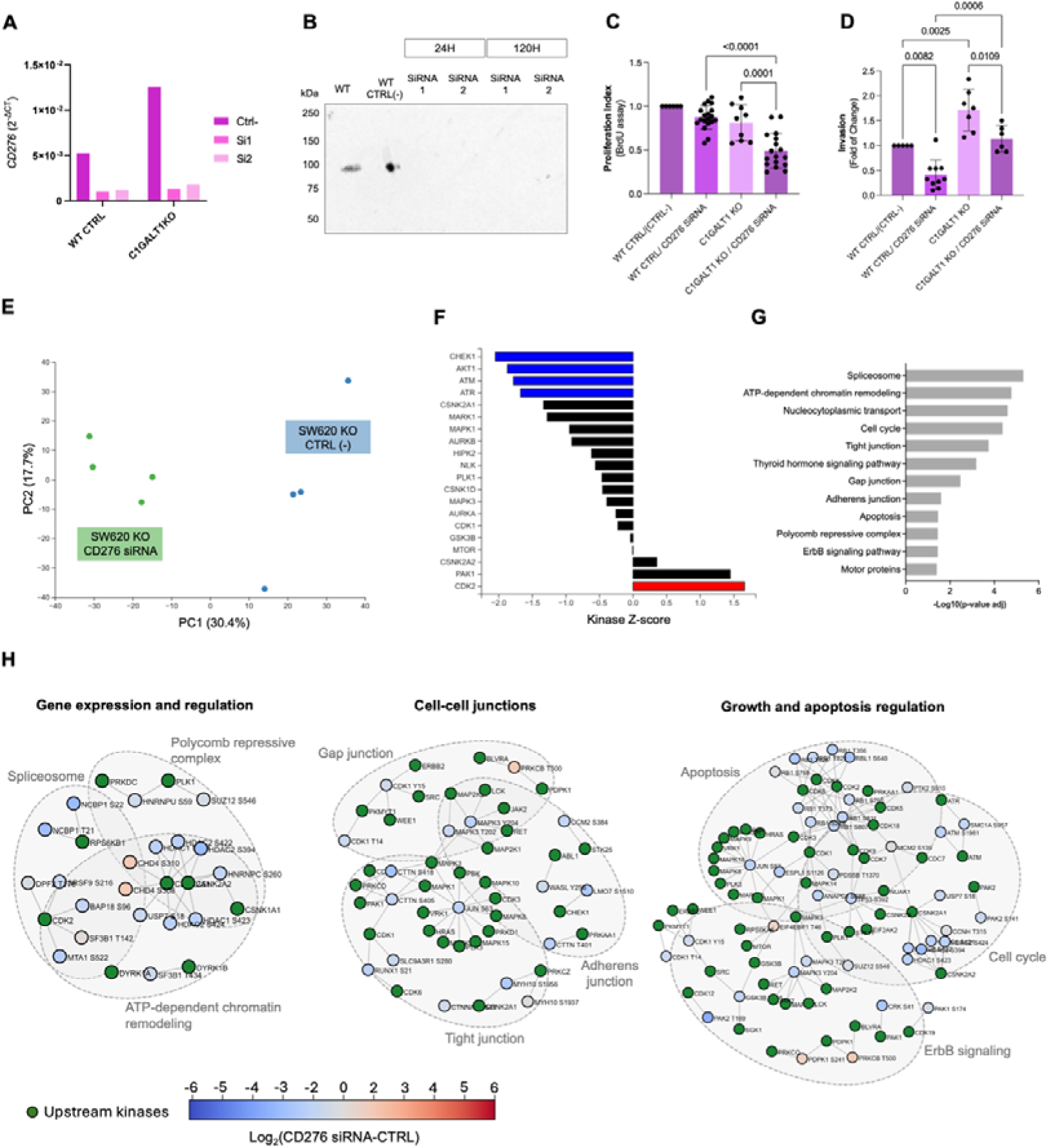
**A-B. *CD276* silencing using siRNA effectively reduces gene and protein expression in both wild-type and *C1GALT1* KO CRC cells.** qPCR shows ∼85% knockdown of *CD276* transcripts with two independent siRNAs (Si1 and Si2) in both cell models compared to non-targeting control (Ctrl-). Western blot analysis confirms complete abrogation of CD276 protein expression up to 5 days post-transfection. **C. Aberrantly glycosylated CD276 supports CRC cell proliferation in SW620 cells.** In SW620 cells, *CD276* silencing significantly reduced proliferation in *C1GALT1* KO cells in relation to controls (*p*□=□0.0001; one-way ANOVA, Tukey’s post hoc test). Data represents the mean fold change relative to control from three independent experiments. **D. Aberrantly glycosylated CD276 promotes CRC cell invasion in SW620 cells.** CD276 silencing decreased invasion in both wild-type and *C1GALT1*-deficient cells. Notably, in *C1GALT1* KO cells CD276 knockdown reversed the glycosylation-induced increase in invasion, confirming a glycan-dependent pro-invasive role. Data represents fold change relative to control and is derived from three independent experiments (one-way ANOVA, Tukey’s post hoc test). **E-H) Aberrantly glycosylated CD276 sustains phospho-signaling programs that promote proliferation, cytoskeletal remodeling in SW620 cells.** Phosphoproteomic profiling was performed on glycoengineered SW620 CRC cells expressing Tn-modified CD276, with or without *CD276* knockdown. CD276 glycosylation was previously associated with increased proliferation and invasion; here, mass spectrometry and Phosphomatics-based analysis reveal that these phenotypes are underpinned by distinct phospho-regulatory networks. **E. Immature glycosylated CD276 supports a distinct phospho-signaling landscape.** Principal component analysis (PCA) of phosphopeptide intensities revealed a clear separation between *CD276*-silenced and control cells, indicating widespread remodeling. **F. Abnormal CD276 glycosylation promotes kinase signaling programs linked to proliferation and cytoskeletal organization.** Kinase-substrate enrichment analysis (KSEA) revealed reduced activity of kinases such as CHEK1, AKT1, ATM and ATR upon *CD276* knockdown, indicating that these proliferative and motility-associated kinases are supported by immature CD276 glycosylation. In contrast, kinases involved in checkpoint control and stress responses, including CDK2, were activated following *CD276* silencing, consistent with induction of growth restraint and DNA damage surveillance mechanisms. **G. Abnormal CD276 glycosylation maintains signaling across pathways involved in chromatin remodeling, cytoskeletal integrity, and cell survival.** Pathway enrichment analysis of differentially phosphorylated proteins revealed that *CD276* knockdown suppresses phospho-regulated processes such as spliceosome and chromatin remodeling complexes (e.g., NuRD), tight junction organization, nucleocytoplasmic transport, and apoptotic resistance. These programs collectively support enhanced plasticity, proliferation, and invasive behavior observed in SW620 *C1GALT1* KO cells. **H. Abnormal CD276 glycosylation maintains phosphorylation of regulatory hubs that drive proliferation, invasion, chromatin remodeling.** Phosphomatics network analysis identified distinct phospho-signaling modules dependent on glycosylated CD276, which were disrupted upon its silencing. These include: i) Chromatin regulators (HDAC1/2, CHD4, CSNK2A2), promoting transcriptional reprogramming and epigenetic adaptability; ii) Cytoskeletal and junction-associated proteins (CTTN, MAPKs), supporting cell motility and invasiveness; iii) Growth and survival kinases (RB1, MAPK3), reflecting pathways that enable tumor persistence and proliferation. Together, these data demonstrate that aberrant CD276 *O*-glycosylation drives SW620 pro-oncogenic phosphorylation programs that coordinate enhanced proliferation and invasion.

To dissect the mechanistic events supporting these findings we also performed phosphoproteomic profiling of SW620 *C1GALT1* knockout cells with or without CD276 knockdown. Principal component analysis revealed distinct phospho-signaling profiles between CD276-silenced and control glycoengineered cells (**Figure 5E**). Kinase–substrate enrichment analysis showed significantly reduced activity (negative z-scores) for ATR, ATM, AKT1, and CHEK1, which are key regulators of DNA damage response, checkpoint control, and pro-survival signaling (**Figure 5F**) [23]. Inhibition of ATR, ATM, and CHEK1 may compromise DNA repair and genotoxic stress responses, while reduced AKT1 activity is likely to impair cytoskeletal dynamics and cell adhesion [24]. These results suggest that CD276 sustains kinase-driven programs supporting migration, survival, and immune evasion. Interestingly, CDK2 activity showed a modest increase after CD276 knockdown, potentially reflecting compensatory activation of cell cycle progression. Pathway enrichment analysis confirmed CD276’s role in regulating phosphorylation-dependent networks involved in spliceosome assembly, chromatin remodeling, cell cycle progression, and cytoskeletal organization (**Figure 5G**). Collectively, CD276 knockdown led to a coordinated downregulation of kinase activity and downstream signaling modules governing DNA repair, adhesion, and transcriptional regulation, which may reduce tumor cell adaptability and resilience. These molecular alterations were evident in substrate-centered network maps (**Figure 5H**) and align with the phenotypic consequences observed in CD276 loss-of-function models (**Figure 5C and D**), including reduced proliferation and impaired migration. It further suggests weakened immune evasion, which warrants deeper investigation. Network-based mapping of phosphosite changes further reinforced these findings, revealing prominent dephosphorylation of nuclear and chromatin-associated regulators following CD276 knockdown. This included core components of the Nucleosome Remodeling and Deacetylase (NuRD) complex such as CHD4 (S308, S310), HDAC2 (S394), and MTA1 (S522) (**Figure 5H)**. Downregulation of CHD4 phosphorylation is consistent with a shift towards a quiescent chromatin state and reduced proliferative capacity [25]. Also, HDAC2-S394 hypo-phosphorylation is associated with impaired deacetylase activity and induction of growth-inhibitory genes such as *CDKN1A/p21* [26], while MTA1-S522 loss has been linked to reduced cellular adaptability to stress [27]. CD276 knockdown also led to widespread dephosphorylation of cytoskeletal effectors controlling intercellular junctions and migratory behavior. This included CTTN, MYH10, and LMO7, key regulators of lamellipodia dynamics and microtubule remodeling [28]. Their decreased phosphorylation supports a loss of invasive plasticity upon CD276 depletion (**Figure 5H**). We further observed phosphosite losses for RB1 and JUN (S63), both known to be critical for cell cycle progression and apoptosis resistance [29]. Conversely, MAPK3 (ERK1) phosphorylation was sustained or elevated, likely reflecting compensatory ErbB signaling in response to CD276 silencing. However, the net signaling effect favored cell cycle arrest and reduced invasion. Collectively, these findings support a role for immature CD276 glycosylation in sustaining kinase-driven signaling programs that promote proliferation, cytoskeletal plasticity, and resistance to apoptosis. Accordingly, its silencing triggered a coordinated shutdown of these pathways, leading to reduced cell cycle activity, reinforcement of junctional architecture, and chromatin stabilization. By contrast, control cells expressing immaturely glycosylated CD276 retain phosphoproteomic signatures of checkpoint adaptation, alternative splicing, and EMT-like transitions, underscoring CD276’s role as a regulator of invasive reprogramming in colorectal cancer.

Collectively, we have highlighted that CD276 immature glycosylation enhances its oncogenic signaling capacity, promoting CRC cell proliferation, invasion, and resistance to stress through kinase-driven phosphoproteomic reprogramming.

### Aberrantly Glycosylated CD276 Promotes Immune Evasion in CRC

To investigate whether the glycosylation status of CD276 modulates its immune regulatory functions, we first conducted *in vitro* experiments by co-culturing pre-activated, carboxyfluorescein succinimidyl ester (CFSE)-labelled human T cells with SW620 wild-type and *C1GALT1* knockout cells, with or without CD276 knockdown. Flow cytometry analyses revealed that T cells co-cultured with *C1GALT1* KO cells exhibited significantly reduced proliferation, as indicated by higher CFSE mean fluorescence intensity in both CD4⁺ and CD8⁺ subsets (**Figure 6B**). This was accompanied by a marked reduction in early (CD69) and late (CD25) activation markers across both CD4^+^ and CD8^+^ T cell subsets (**Figure 6A, Figure S9**). Notably, these immunosuppressive effects were reversed upon CD276 knockdown in the *C1GALT1* KO background, strongly implicating aberrantly glycosylated CD276 as a key mediator of T cell inhibition.

**Figure 6.**
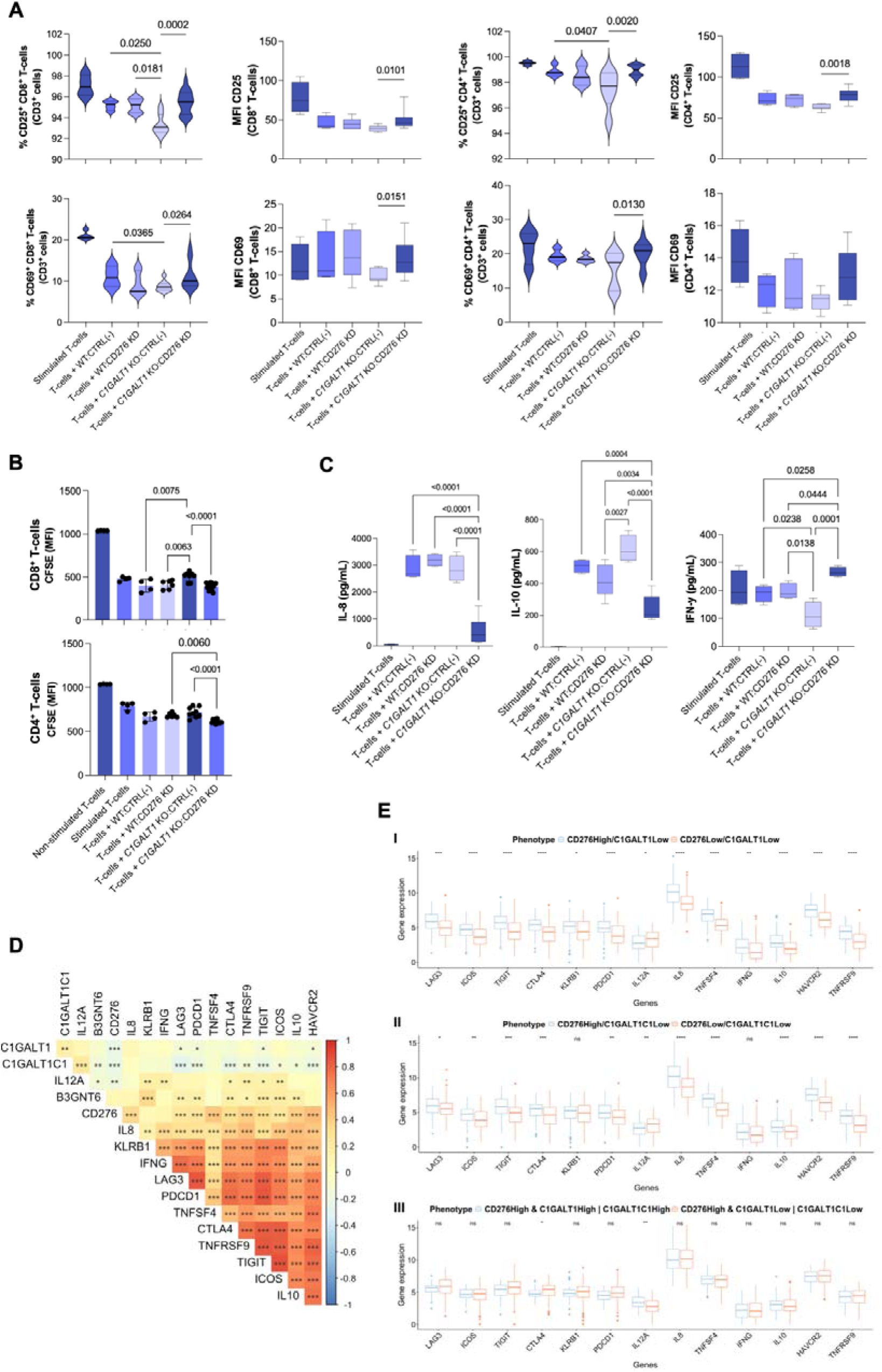
**Aberrant CD276 glycosylation suppresses T cell function and shapes an immunosuppressive tumor microenvironment. A–C) Aberrant CD276 glycosylation dampens T cell activation, proliferation, and functional cytokine output.** *In vitro* co-culture assays and cytokine profiling were conducted using SW620 colorectal cancer cells glycoengineered with *C1GALT1* KO (resulting in immature CD276-Tn glycoforms) and wild-type controls, with or without *CD276* knockdown. These cell models were co-cultured with pre-activated human T cells to evaluate functional immune responses. Across three independent replicates, T cell activation, proliferation, and cytokine secretion were significantly influenced by the glycosylation status of CD276. Statistical comparisons were performed using one-way ANOVA followed by Tukey’s multiple comparisons test. **A. Aberrant CD276 glycosylation dampens T cell activation.** Flow cytometry revealed a significant reduction in CD25^⁺^ cells within both CD4^⁺^ and CD8^⁺^T cell subsets after 5 days of co-culture with *C1GALT1* KO cells. Mean fluorescence intensity (MFI) of CD25 was also lower in these T cells, indicating a subdued activation profile. CD276 knockdown in the KO background restored CD25 expression (both percentage and MFI), implicating glycosylated CD276 in the suppressive phenotype. Similarly, both percentage of CD69^⁺^ cells and MFI were reduced in T cells after exposure to CD276-expressing KO cells, further supporting impaired activation. **B. Abnormal CD276 glycosylation inhibits T cell proliferation.** CFSE dilution assays showed higher fluorescence retention in both CD4^⁺^ and CD8^⁺^T cells co-cultured with *C1GALT1* KO cells, indicating reduced proliferation. Silencing CD276 rescued CFSE dilution, confirming that the glycosylated form of CD276 actively suppresses T cell proliferative responses. **C. Abnormal CD276 glycosylation reprograms cytokine secretion toward an immunosuppressive profile.** Supernatants from co-cultures with *C1GALT1* KO cells displayed elevated IL-8 and IL-10 and reduced IFN-γ levels, compared to *C1GALT1* KO *CD276* KD cells, supporting a shift toward immune suppression. D. Expression correlations suggest an association between CD276, core 1 *O*-glycosylation genes, and immune regulatory markers in patient samples. A correlogram of TCGA CRC samples revealed strong positive correlations between CD276 and immune checkpoint molecules associated with T cell exhaustion, including *PDCD1* (PD-1) and *HAVCR2* (TIM-3). Additionally, *CD276* expression was inversely correlated with *C1GALT1* and *C1GALT1C1*, further reinforcing that impaired core 1 *O*-glycosylation may be linked to enhanced CD276 expression and transcriptional profiles consistent with immune evasion. E. Tumours co-expressing high *CD276* and low levels of core 1 *O*-glycosylation enzymes exhibit transcriptional profiles indicative of T cell dysfunction and immune suppression. Analysis of TCGA CRC samples revealed that tumours with high *CD276* and low *C1GALT1* expression (**Panel I**) displayed a marked increase in genes associated with T cell exhaustion and elevated immunosuppressive cytokines *IL10* and *IL8*, closely mirroring the phenotypes observed in *in vitro* co-culture assays. **Panel II** reinforces this observation, showing that tumours with *CD276*^High^/*C1GALT1C1*^Low^ expression recapitulate the immunosuppressive phenotype. **Panel III** extends this observation by stratifying tumours co-expressing *CD276*^High^ with low expression of both *C1GALT1* and its chaperone *C1GALT1C1* (with high probability of expressing high levels of immature glycosylation), confirming a broader immunosuppressive transcriptional program that includes increased expression of exhaustion marker such as *CTLA4* and reduced pro-inflammatory cytokine IL12A.

Furthermore, T cells co-cultured with *C1GALT1* KO cells exhibited a cytokine profile skewed toward an immunosuppressive and regulatory phenotype, with significantly elevated levels of IL-10 and IL-8, and a corresponding decrease in IFN-γ (**Figure 6C and Figure S10**). Given that IL-8 and IL-10 are not canonical T cell cytokines, their elevation may also originate from cancer cells and reflect a contact-dependent response to T cell engagement modulated by the glycosylation status. Broad-spectrum cytokine profiling further revealed a consistent increase in the secretion of pro-inflammatory mediators, including IL-6, IL-23, IL-17A, and TNF-α, which can either stimulate immune surveillance or drive tumor-promoting inflammation. Notably, elevated IL-23 and IL-17A levels in CRC have been frequently linked to tumour growth and angiogenesis [30]. This is accompanied by increased levels of type 2 and tolerogenic cytokines, such as IL-13 and IL-5 (**Figure S10**), indicating a shift away from Th1/effector polarization toward a more immune-modulatory state. Importantly, these cytokine alterations were largely reversed upon CD276 knockdown in the *C1GALT1* KO background, restoring IFN-γ production and attenuating IL-10 and IL-8 secretion (**Figure 6C**). Together, these findings support a model in which immature CD276 glycosylation promotes immune escape by shaping a non-permissive cytokine environment. Although T cells were pre-activated, direct contact with *C1GALT1*-KO cells in co-culture suppressed their activation and prevented them from acquiring effector functions. This aligns with previous report showing that cancer cells can respond to T cell-derived inflammatory cues, such as TNF-α via NF-κB, by increasing IL-8 production, a chemokine known to recruit MDSCs and TAMs, promote angiogenesis, and reinforce immune exclusion [31]. In parallel, IL-10 suppresses T cell function and promotes peripheral tolerance [32], while diminished IFN-γ reflects impaired effector polarization of CD4⁺ T cells and reduced cytotoxic capacity of CD8+ T cells [33]. In summary, our data define a glycosylation-dependent mechanism of immune modulation, whereby the acquisition of immature glycosylation by CD276 in cancer cells contributes to the establishment of a suppressive cytokine milieu and an immune-excluded, tumor-permissive niche.

We then interrogated TCGA CRC datasets to find support for these findings in clinical samples. Correlation and clustering analyses revealed that low expression of the glycosyltransferases *C1GALT1* and *C1GALT1C1* correlated with increased expression of key immune checkpoints and T cell exhaustion markers, including PD-1 (*PDCD1*) and TIM-3 (*HAVCR2*) (**Figure 6D**) [34]. In contrast, *B3GNT6* expression showed weaker positive correlations with immune-related genes, suggesting a more limited role in modulating immune escape. CD276 expression itself was positively correlated with multiple immune checkpoints such as *PDCD1* and *HAVCR2*, and inversely correlated with *C1GALT1* and *C1GALT1C1* (**Figure 6D**), as previously described. To explore this further, we stratified tumors based on *CD276* and *C1GALT1* expression levels to mirror the *in vitro* phenotype for CD276 with immature glycosylation. Tumors with high *CD276* and low *C1GALT1* expressions (*CD276*^High^/*C1GALT1*^Low^) exhibited significantly elevated levels of exhaustion markers including *TIGIT*, *LAG3*, *HAVCR2*, and *PDCD1*, when compared to *CD276*^Low^/*C1GALT1*^Low^ tumors (**Figure 6E, Panel I**). A similar pattern emerged when stratifying by *C1GALT1C1* expression (*CD276*^High^/*C1GALT1C1*^Low^ vs. *CD276*^Low^/*C1GALT1C1*^Low^; **Figure 6E, Panel II**), reinforcing that deficiencies in core 1 *O*-glycosylation whether *via* C1GALT1 or its chaperone potentiate CD276-associated immune evasion. Cytokine gene expression analysis further supported the presence of a reprogrammed immune environment. *IL10* and *IL8* levels were significantly elevated, while *IL12A* was reduced in both *CD276*^High^/*C1GALT1*^Low^ and *CD276*^High^/*C1GALT1C1*^Low^ tumors, mirroring cytokine shifts observed *in vitro*. Interestingly, *IFNG* expression was only elevated in *CD276*^High^/*C1GALT1*^Low^ tumors, suggesting that T cells may become initially activated but subsequently driven toward exhaustion due to sustained immune checkpoint signaling, which warrants future demonstration. By contrast, when *CD276* expression remained high but tumors were stratified by high or low *C1GALT1*/*C1GALT1C1* expression, most immune-related genes were no longer differentially expressed, except for CTLA4 and IL12A (**Figure 6E, Panel III**). This suggests a strong influence of CD276 in driving immunosuppression, with impaired O-glycosylation amplifying this effect. Together with *in vitro* data, these transcriptomic associations suggest that aberrant *O*-glycosylation enhances the immunomodulatory capacity of CD276 and contributes to the establishment of an immunosuppressive program in CRC.

## Discussion

The glycome is pivotal in cancer progression and dissemination, offering significant potential for the precise detection, targeting, and elimination of cancer cells. Despite recent advances in understanding CRC glycome alterations by high-throughput mass spectrometry, a considerable gap in grasping the underlying functional and clinical implications of these alterations remains [22, 35]. This study addresses these gaps through an in-depth exploration of *O*-glycosylation and its impact in colorectal cancer.

We confirm previous findings linking premature termination in *O*-glycan elongation to cancer aggressiveness [36]. We began by identifying glycogene deregulation in colorectal tumours compared to normal tissue, with *B3GNT6* downregulation emerging as a poor prognosis marker in CRC. This, when was frequently accompanied by concomitant lower expression of *C1GALT1* and/or *C1GALT1C1* downregulation, also had a significant impact on the decreased overall survival of CRC patients. Advancing beyond current knowledge, we demonstrate that these events coincide with the loss of normal colonic glycosylation, predominantly characterized by *O*-6 sialylated core 3 structures, likely supported by *B3GNT6* and *ST6GALNACs* expression in healthy tissues. Moreover, we found that advanced epithelial-like and mesenchymal-like CRC showed unique glycocode profiles. Epithelial-like tumours displayed an intermediate glycome between normal colon tissue and mesenchymal-like cancers. In these cases, the loss of core 3-related structures was not compensated by the capacity to shift the glycophenotype toward core 1 and core 2-related structures, resulting in an enrichment for short Tn and sTn antigens. Mesenchymal-like tumours, while also exhibiting high levels of Tn and sTn antigens, were predominantly enriched for sialylated T antigens. Interestingly, tumours of both differentiation states that showed signs of distant metastases also displayed extended core 2 structures, including glycans compatible with the presence of sLe antigens. These antigens are known to facilitate cancer cell intravasation into the bloodstream and colonization of distant locations by mediating adhesion to endothelial cells via E-selectin [37]. These findings reinforce previous reports and underscore the heterogeneous and dynamic nature of the CRC glycome, where distinct subpopulations may coexist to drive aggressive traits [35b, 38]. We further broadened our understanding of the events potentially dictating immature glycosylation at the surface of CRC cells. Here, we show that in more aggressive tumours associated with an unfavorable prognosis, the concurrent downregulation of *B3GNT6* and *C1GALT1* play a decisive role in driving immature glycosylation, which had previously been largely attributed to *C1GALT1C1* loss-of-function mutations [39]. Utilizing *C1GALT1* knockout cell lines with distinct molecular backgrounds, we confirmed that immature glycosylation drives enhanced invasion and, in a context-dependent manner, increases proliferation. This reinforces previous findings regarding *C1GALT1C1* and decisively links the acquisition of immature glycophenotypes through different mechanisms to cancer aggressiveness [40].

We finally identified CD276 as a carrier of immature *O*-glycans in CRC. Also known as B7-H3, CD276 belongs to the B7 family of immune checkpoint molecules and can either enhance or suppress immune responses depending on context [41]. Namely, it has been shown to interact with cognate receptors on immune cells, particularly T cells, regulating their activation and function [41]. CD276 is overexpressed in several malignancies, including CRC [9], where its expression correlates with poor prognosis. Our findings go beyond the state-of-the-art by demonstrating that immature *O*-glycosylation also regulates CD276 stability, enhances its expression, and shapes its immunomodulatory role in CRC. This underscores that CD276 regulation in cancer is not solely governed by *N*-glycans but is also critically shaped by the *O*-glycome, revealing new layers of regulatory control and therapeutic potential. Specifically, we show that aberrantly *O*-glycosylated CD276 is abundant in primary CRC tumours and metastases but absent from healthy colon tissue, suggesting a cancer-specific modification. Moreover, expression of CD276 carrying immature *O*-glycans was enriched in right-sided tumours, which are commonly associated with more aggressive clinical behavior, and was linked to significantly worse patient survival. Proteomic and glycoproteomic analyses further suggest that, while CD276 is broadly expressed across tumour stages and differentiation states, the density and distribution of its glycosylation sites vary with tumour differentiation, highlighting potential implications for tumour behavior and therapeutic design. These findings underscore the need for dedicated site-specific glycoproteomic methodologies, including glycopeptide enrichment strategies and electron-transfer-based fragmentation, to resolve precise glycosylation sites and understand their structural and functional consequences. Such insights could refine CD276-targeted strategies by accounting for glycoform diversity that may influence antibody recognition, ligand binding, or immune evasion mechanisms. We also show that loss or downregulation of *C1GALT1* promotes immature *O*-glycosylation on CD276, leading to increased protein stability and elevated expression. Although our data support a posttranslational stabilization mechanism, the basis for the observed transcriptional upregulation of CD276 remains unclear. One plausible explanation is that defective *O*-glycosylation perturbs cellular homeostasis, activating compensatory stress response pathways. Previous studies have shown that glycosylation defects can induce endoplasmic reticulum stress or Golgi dysfunction, triggering transcriptional programs that upregulate membrane protein expression [42]. CD276 may therefore be indirectly upregulated as part of a broader adaptive response, which warrants further exploration.

Additionally, emerging evidence suggests that glycosylation status can modulate epigenetic and transcriptional regulators. Altered glycosylation has been shown to affect the stability and function of key transcription factors such as SP1 and STAT3, both implicated in immune checkpoint regulation [43]. It is conceivable that similar mechanisms contribute to increased CD276 transcription in CRC. Although the precise pathways remain to be defined, our findings lay the groundwork for future investigations into how glycome remodeling interfaces with CD276 transcriptional control in cancer.

Functionally, we demonstrated that immature glycosylation supports CD276-mediated invasion through phospho-signaling rewiring, enhanced cell proliferation, and suppressed T cell activation. *In vitro*, *CD276* knockdown in glycoengineered cells reduced both invasive capacity and proliferation, particularly in metastatic models. Quantitative phosphoproteomic analysis revealed that aberrantly glycosylated CD276 sustains pro-invasive and pro-proliferative signaling through kinases such as CSNK2A2, AKT1, and CDK2, and modulates chromatin and cytoskeletal regulators, including HDAC2, CHD4, and cortactin [23, 44]. These findings define a cell-intrinsic function of CD276, wherein its abnormal glycosylation promotes tumour aggressiveness through activation of oncogenic signaling cascades.

Aberrant CD276 glycosylation also reprogrammed cytokine profiles, promoting IL-10 and IL-8 production while decreasing pro-inflammatory IFN-γ secretion. These changes suggest that aberrantly glycosylated CD276 not only impairs T cell activation but also actively reconditions the tumour microenvironment to favor immune evasion [31–32, 45].

Together, these results position glycosylated CD276 as a multifunctional effector, orchestrating both tumour-intrinsic signaling and immune suppression. Importantly, TCGA data corroborated these findings at the clinical level, showing that tumours with high *CD276* and low *C1GALT1* expression exhibited reduced expression of T cell activation and cytotoxicity markers, and higher levels of immunosuppressive cytokines such as IL10, reinforcing the translational relevance of our findings. This intersection of intracellular and immunomodulatory effects likely provides a strong selective advantage to cancer cells, enabling both aggressive growth and immune escape.

In conclusion, our findings establish a mechanistic and functional framework linking glycosylation-driven CD276 remodeling to tumour aggressiveness and immune escape in colorectal cancer. Collectively, we position CD276 *O*-glycosylation remodeling as a novel driver of immune checkpoint activity and tumour aggressiveness in CRC. Future studies should focus on providing definitive demonstrations of the role of abnormally glycosylated CD276 as an immune checkpoint in CRC by exploiting *in vivo* models. Investigation into the impact of other CRC glycans, including extended core 2 structures associated with distant metastasis [22], on CD276 functions is also warranted. Moreover, gaining a deeper understanding of CD276 impact on other immune cells and its role in disease pathology will be key for designing appropriate countermeasures. Ultimately, this work provides foundational insights for the rational design of therapeutic strategies that selectively target aberrantly glycosylated CD276 in cancer. Together, our findings unveil a previously unrecognized link between cancer-associated glycome remodeling and immune evasion in colorectal cancer. By suggesting CD276 as a glycosylation-dependent immune checkpoint, this study opens new avenues for innovative therapeutic strategies aimed at selectively targeting tumour-specific glycoforms. Future efforts to decode the broader “glycocode” of the tumour microenvironment, including the identification of receptors and lectin networks that interpret this altered glycosylation landscape. This will be critical to unlocking next-generation immunotherapies and advancing precision oncology.

## Materials and Methods

### Clinical Tissue Samples and Ethical Compliance

A retrospective cohort of 40 formalin-fixed paraffin-embedded (FFPE) CRC tissue samples was selected from the institutional pathology biobank of the Portuguese Institute of Oncology of Porto (IPO Porto). Samples were collected from patients who underwent surgical resection for colorectal (adeno)carcinomas between 2005 and 2012. The cohort included 18 female and 22 male patients, aged 28 to 76 years (mean ± SD: 61□±□11 years). Whenever present, adjacent histologically normal mucosa was also included for comparative analysis. In addition to CRC samples, a reference panel of healthy tissues, including liver, stomach, pancreas, appendix, lung, testis, thyroid, and gallbladder, was analyzed. Tumour and healthy tissue sections were further screened for CD276 and aberrant glycosylation markers (Tn and sTn antigens) using multiple immunoassays: immunohistochemistry (IHC), PLA, and dual immunofluorescence staining with lectins and specific antibodies. All procedures were conducted in compliance with institutional ethical guidelines and approved by the IPO-Porto Ethics Committee (project reference: CES 86/017). Written informed consent was obtained from all patients. Clinical and pathological data were retrieved from patient medical records and are summarized in **Table 1**. Antibodies and experimental conditions are detailed in **Table S5.**

**Table 1.** Clinicopathological data associated with tumour tissues assessed in this study (n = 40).

|  | *n (%)* |
| --- | --- |
| **Stage** |  |
| I | 1 (2.5) |
| II | 4 (10.0) |
| III | 10 (25.0) |
| IV | 25 (62.5) |
| **Tumour (T)** |  |
| T1 | 1 (2.5) |
| T2 | 1 (2.5) |
| T3 | 31 (77.5) |
| T4 | 6 (15.0) |
| Missing Information | 1 (2.5) |
| **Lymph node metastases (N)** |  |
| N0 | 11 (27.5) |
| N1 | 12 (30.0) |
| N2 | 14 (35.0) |
| N3 | 3 (7.5) |
| **Tumour Location** |  |
| Right colon | 7 (17.5) |
| Left colon | 23 (57.5) |
| Rectum | 10 (25.0) |

### Transcriptomic Analysis of the TCGA-COADREAD Cohort

RNA sequencing (RNA-seq) data for CRC were obtained from The Cancer Genome Atlas (TCGA) COADREAD cohort, comprising 623 tumor tissues and 51 matched normal adjacent tissues. Processed gene expression data (log2(FPKM + 1)) and associated clinical annotations - including age, gender, tumor stage, histological subtype, and overall survival - were downloaded via the UCSC Xena platform (https://xena.ucsc.edu/), which hosts curated TCGA datasets. After excluding samples with incomplete clinical data, a final cohort of 598 tumor samples and 51 normal adjacent tissue samples was included for analysis. Clinical characteristics of the COADREAD patients are summarized in **Table 2**.

**Table 2.**
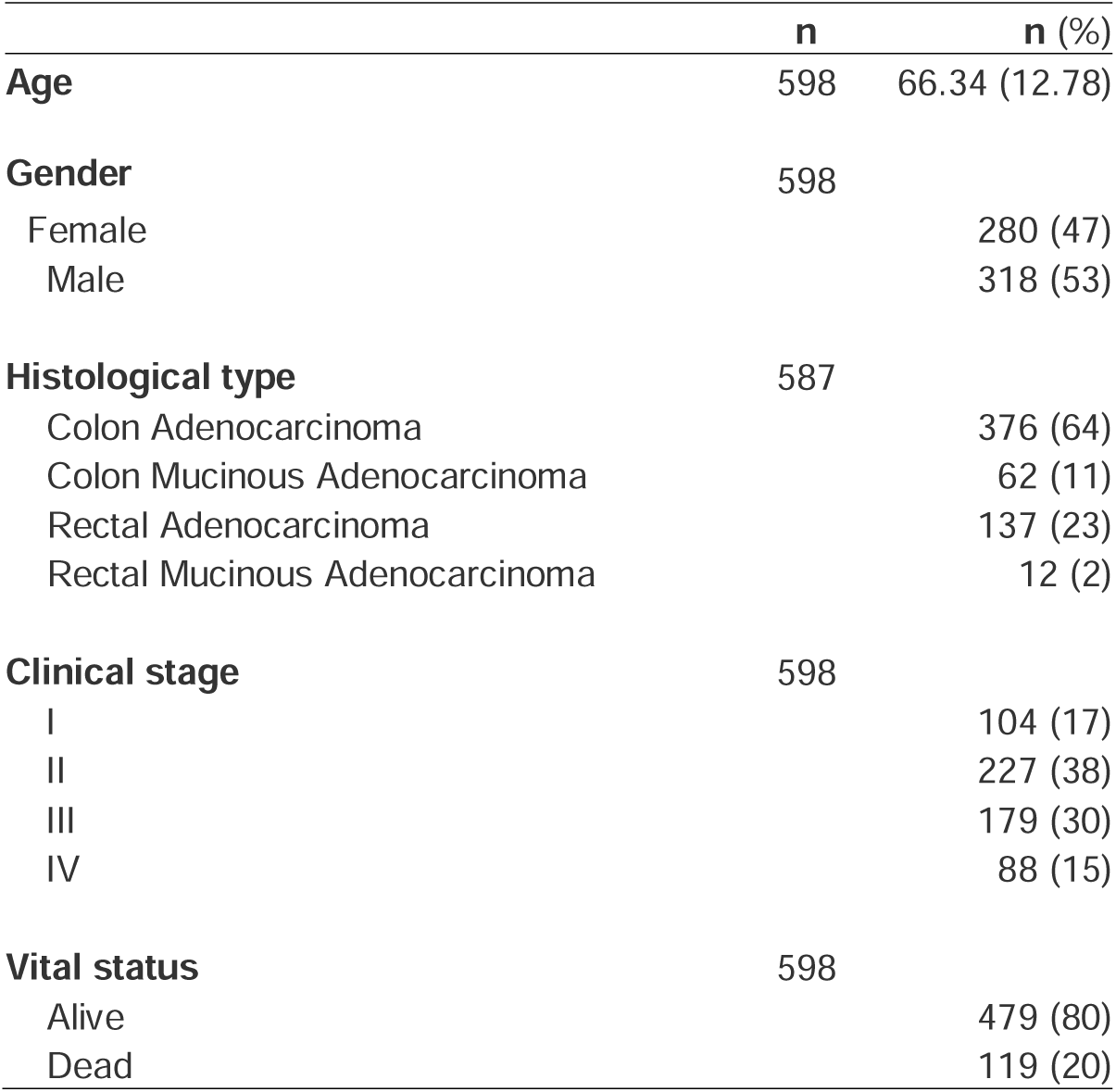
Clinicopathological characteristics of CRC cases included in the TCGA COADREAD dataset.

### Proteomic Revisitation of TCGA-COADREAD for CD276 Glycosylation

A subset of 95 colorectal cancer (COADREAD) cases from TCGA, analyzed by mass spectrometry and annotated with clinicopathological data in Zhang *et al.* [46], was retrieved from the PRIDE Archive (dataset identifier PXD002080). The proteomics data were reanalyzed to identify and quantify glycoproteins modified with Tn and sialyl-Tn (sTn) antigens. Samples were stratified into epithelial-like and mesenchymal-like subtypes according to the TCGA transcriptomic classification used by Zhang *et al.* [46]. Further classification was based on the presence of unglycosylated CD276 or CD276 glycosylated with Tn/sTn antigens (CD276-Tn/sTn). All data processing and visualization were performed in R version 4.2.3. Data wrangling and statistical plotting were conducted using the tidyverse [47] and ggpubr packages. CD276-Tn/sTn–positive samples were manually curated to identify candidate glycosylation sites specific to epithelial- and mesenchymal-like subtypes. These glycosites were visualized as a heatmap using the pheatmap package, enabling subgroup clustering and the identification of potential molecular patterns.

### Cell Lines and Culture Conditions

Human CRC cell lines SW480, SW620, HCT118, RKO, LS174T and HCA7 were obtained from the American Type Culture Collection (ATCC). Cells were cultured in RPMI 1640 GlutaMAX™ medium (Gibco) supplemented with 10% fetal bovine serum (FBS) and 1% penicillin-streptomycin at 37□°C in a humidified incubator with 5% CO₂. Cells were screened weekly for Mycoplasma contamination.

### CRISPR-Cas9 glycoengineered cell models

Two guide RNAs targeting the human *C1GALT1* gene (sequences: *GTAAAGCAGGGCTACATGAG* and *ACAACACTTTGTTACAACGC*) were cloned into a dual-guide RNA CRISPR-Cas9 expression vector (pRP[2CRISPR]-Puro-hCas; VectorBuilder). Wild-type SW480 and SW620 CRC cell lines were transfected with CRISPR-Cas9 plasmid (1□µg) using jetPRIME® transfection reagent (Polyplus) according to the manufacturer’s protocol. Single-cell clones were isolated by limiting dilution in 96-well plates, and successful *C1GALT1* KO clones were identified by Indel Detection by Amplicon Analysis (IDAA) using the ABI PRISM™ 3010 Genetic Analyzer (Thermo Fisher Scientific) and confirmed by Sanger sequencing. Three independent KO clones with distinct out-of-frame indels were selected for further studies. In addition, clones harboring silent mutations were used as phenotypic control cell lines. IDAA results were analyzed using Peak Scanner Software v1.0 (Thermo Fisher Scientific).

### siRNA-Mediated CD276 Silencing

*CD276* gene silencing was performed in SW620 glycoengineered colorectal cancer cell models using reverse transfection with siRNA. Two Silencer® Select siRNAs targeting distinct exons of human CD276 (s37289, exon 10; and s37288, exon 7; Invitrogen) were used to ensure target specificity. A Silencer® Select negative control siRNA (catalog no. 4390843, Invitrogen) was included in all experiments as a non-targeting control, as previously described [48]. Cells were detached and seeded into 24-well plates at a density of 1□×□10□□cells/well prior to transfection. siRNAs and Lipofectamine® RNAiMAX (Invitrogen) were separately diluted in Opti-MEM® Reduced Serum Medium (Gibco), incubated for 5□minutes at room temperature, and combined to form siRNA–lipid complexes. Cells were then incubated with the complexes for 72□hours at 37□°C. Each condition was plated in duplicate wells per experiment, and all experiments were performed in at least three independent biological replicates. To evaluate gene silencing efficiency, *CD276* mRNA expression was quantified by RT-qPCR using the TaqMan® Gene Expression Assay Hs00987207_m1 (Thermo Fisher Scientific). *GAPDH* and *ACTB* were used as endogenous controls. To confirm the specificity and rule out off-target effects, both non-targeting negative control siRNA and mock-transfected cells (no siRNA, Lipofectamine only) were included as negative controls. Effective knockdown was confirmed only when both *CD276*-targeting siRNAs produced consistent reductions in mRNA levels compared to controls. Cell viability and morphology were also monitored microscopically to exclude transfection-related cytotoxicity.

### Proliferation assay

Cell proliferation was assessed using the Cell Proliferation ELISA, BrdU (colorimetric) kit (Roche), according to the manufacturer’s instructions. Absorbance was measured at 450□nm using a microplate reader. Each assay was performed in experimental triplicates with three technical replicates per condition.

### Invasion Assay

Invasion potential was evaluated using Falcon® Permeable Supports for 24-well plates with 8.0□µm transparent PET membranes (Corning), coated with Matrigel® Basement Membrane Matrix (50□µg/mL, Corning). Inserts were pre-coated and incubated at 37□°C for 1□h prior to cell seeding. Subconfluent cells were detached using trypsin/EDTA (Thermo Fisher Scientific) and seeded in the upper chamber at a density of 5□×□10□□cells/mL. After 24□h of incubation at 37□°C, non-invading cells were removed from the upper surface, and the inserts were washed with PBS and fixed in 4% paraformaldehyde (Sigma-Aldrich). Filters were mounted in Vectashield with DAPI (Vector Laboratories) and imaged using a Zeiss Axiovert 200M fluorescence microscope (Carl Zeiss). Each assay was performed in experimental triplicates with five technical replicates per condition. Results were normalized to cell proliferation and expressed as fold change relative to control.

### RT-qPCR for CD276 and Glycogene Expression

Total RNA was extracted using TriPure Isolation Reagent (Roche Diagnostics). cDNA synthesis and mRNA quantification were performed as previously described [49]. Quantitative RT-PCR was carried out using TaqMan Gene Expression Assays for CD276 (Hs00987207_m1) and B3GNT6 (Hs00371066_s1) on a 7500 Fast Real-Time PCR System (Applied Biosystems). GAPDH (Hs03929097_g1) and ACTB (Hs99999903_m1) were used as endogenous controls. All samples were analyzed in duplicate and relative gene expression was calculated using the 2^–ΔCt method.

### Western Blot for CD276 and Glycan Detection

Total and plasma membrane protein extracts, as well as CD276-immunoprecipitated fractions, were analyzed by Western blotting. Total protein lysates were prepared using a buffer containing Tris-HCl (25□mM, pH 7.4), NaCl (150□mM), MgCl₂ (5mM), 1% NP-40, and 5% glycerol, supplemented with Halt™ Protease and Phosphatase Inhibitor Cocktail (Thermo Fisher Scientific). Plasma membrane proteins were isolated by ultracentrifugation, as described by Fernandes et al. [50]. Proteins were resolved by SDS-PAGE (4–20% precast gels, Bio-Rad) and transferred to nitrocellulose membranes (Cytiva). Membranes were probed with anti-CD276 antibodies and biotinylated Vicia villosa agglutinin (VVA) lectin (Vector Laboratories) to detect GalNAc residues associated with the Tn antigen. Lectin binding was visualized using streptavidin-HRP and chemiluminescent detection. The results were normalized to total protein content given by Ponceau S staining. Antibodies, lectins, and experimental conditions are detailed in **Table S5**.

### CD276 Immunoprecipitation

CD276 was immunoprecipitated from plasma membrane protein extracts using Pierce™ Protein G Agarose beads (Thermo Fisher Scientific). Beads were pre-blocked with 1% BSA (Sigma-Aldrich) for 1□h at 4□°C. Protein extracts were precleared with BSA-blocked beads for 2□h at 4□°C to reduce nonspecific binding. Supernatants were incubated with CD276 polyclonal antibody (10□µg) at 4□°C for 2□h, followed by overnight incubation with fresh blocked beads. After washing, immune complexes were eluted in SDS loading buffer at 95□°C. Eluted proteins were resolved on 4–20% gradient SDS-PAGE gels (Bio-Rad) and transferred to nitrocellulose membranes (Cytiva). Immunodetection was performed with anti-CD276 antibody and VVA lectin, as detailed in **Table S5.**

### Flow Cytometry for CD276 and Glycans Detection

Flow cytometry was used to assess the expression of Tn, sTn, T, and sT antigens, as well as CD276, as previously described by Peixoto et al. [51]. Detailed antibody information and staining conditions are listed in **Table S5**. To validate sialylated glycans specificity, isotype controls and cells treated with 70□mU of α-neuraminidase (*Clostridium perfringens*, Sigma-Aldrich) were used as negative controls. Data were acquired on a Beckman Coulter FC500 flow cytometer and analyzed using CXP software (Beckman Coulter).

### Protein Stability Assay Using Cycloheximide Chase

To assess CD276 protein stability, 1 × 10□ SW480 and SW620 colorectal carcinoma cells were seeded per well in 6-well plates and cultured overnight in complete RPMI 1640 GlutaMAX™ medium (Gibco). Cells were treated with cycloheximide (20□μM, CHX) (Sigma-Aldrich), dissolved in DMSO, to inhibit de novo protein synthesis. Control wells were treated with an equivalent volume of DMSO. Whole-cell protein extracts were collected overtime for 24 hours post-treatment using RIPA buffer (50□mM Tris-HCl, pH□8.0; 150□mM NaCl; 1% NP-40; 0.5% sodium deoxycholate; 0.1% SDS) freshly supplemented with Halt™ Protease and Phosphatase Inhibitor Cocktail (Thermo Fisher Scientific). For each time point, 10□µg of protein were resolved by SDS-PAGE on 4–20% precast gradient gels (Bio-Rad) and transferred onto nitrocellulose membranes (Cytiva). Protein transfer and equal loading were verified by staining with Ponceau S solution (Sigma-Aldrich). Membranes were blocked and immunoblotted using primary antibodies against CD276 and appropriate HRP-conjugated secondary antibodies, as detailed in **Table S5**.

### *O*-Glycomics on Cell Models and Patient Samples

*O*-glycome characterization of CRC and matched healthy tissues was performed on 10□µm FFPE sections. *O*-glycans were released after *N*-glycan extraction according to the described by Relvas-Santos et al [52]. Namely, on-tissue reductive β-elimination (1□M NaBH₄ in 50□mM KOH, overnight at 50□°C) was employed for *O*-glycan release, followed by desalting using cation-exchange resin (AG 50W-X8; Bio-Rad). For CRC cell lines, *O*-glycans were profiled using the Cellular *O*-Glycome Reporter/Amplification (CORA) approach [53], following the protocol of Fernandes *et al.* [50]. Released glycans were dried and permethylated as previously described by us [52]. nanoLC-MS/MS analysis was performed using an Ultimate 3000 nano-LC system coupled to a Q Exactive mass spectrometer with an EASY-Spray nano-electrospray source (Thermo Fisher Scientific). Eluent A was 0.2% formic acid in water; eluent B was 0.2% formic acid in acetonitrile. Samples were loaded onto a PepMap C18 trap column (5□µm, Thermo Fisher Scientific) and washed isocratically (90% A, 10% B, 30□µL/min). After 3□min, flow was redirected to the analytical column (PepMap C18, 100□Å, 150□mm × 75□µm, 3□µm particle size) operated at 0.3□µL/min and 35□°C. Glycan separation was achieved with a linear gradient: 10% B at 10□min, 38% B at 20□min, 50% B at 55□min, and 90% B at 65□min. The column was held at 90% B for 10□min before re-equilibration. Mass spectrometry (MS) was performed in positive ion mode with a scan range of *m/z* 400–2000 for analysis of cell lines and *m/z* 280–2000 for analysis of *O*-glycans extracted from tissues, 120,000 resolution (Full MS), spray voltage of 1.9□kV, and capillary temperature of 275□°C. Data-dependent MS/MS (Top 15) was acquired at 30,000 resolution using higher-energy collisional dissociation (HCD, 30% normalized collision energy, 4.0□*m/z* isolation window), with a 45□s dynamic exclusion. Data analysis was performed using Xcalibur v3.0 (Thermo Fisher Scientific), applying Gaussian smoothing, 10□ppm mass tolerance, four decimal mass precision, and default baseline subtraction. Glycan structures were assigned considering retention time, monoisotopic m/z, and MS/MS spectra. The relative abundance resulted from the sum of the extracted ion chromatogram areas for each glycan structure in relation to the sum of chromatographic areas of all identified glycans. Isomeric and isobaric structures were not differentiated. Glycan representations were created using GlycoWorkBench v2.1.

### Phosphoproteomics of Cell Models

Glycoengineered SW620 cells, with or without CD276 knockdown by siRNA, were harvested for whole protein extraction by cell scrapping using lysis buffer (1% sodium deoxycholate, 10 mM tris(2-carboxyethyl)phosphine hydrochloride), 40 mM chloroacetamide, and 100 mM TRIS) supplemented with Halt™ Protease and Phosphatase Inhibitor Cocktail (1x) (Thermo Fisher Scientific). Lysates were heated at 95 °C and sonicated in an ultrasound bath. After protein quantification, 200□μg of total protein was digested overnight at 37□°C using trypsin (Promega; 1□μg per 50□μg of protein) in 50□mM ammonium bicarbonate. Sodium deoxycholate was precipitated with 2% formic acid by centrifugation. Dried samples were desalted using Pierce™ Peptide Desalting Spin Columns (Thermo Fisher Scientific) and enriched for phosphopeptides using TiO₂-based spin columns (High-Select™ TiO2 Phosphopeptide Enrichment Kit, Thermo Fisher Scientific) according to the manufacturers’ protocol. Samples were analyzed using nanoLC-nESI-MS/MS on a Q Exactive™ mass spectrometer (Thermo Fisher Scientific) coupled to a Vanquish neoUHPLC nano-LC system. Eluent A was aqueous formic acid (0.1%) and eluent B was formic acid (0.1%) in 80% acetonitrile. Samples were injected directly into a trapping column (PEPMAP NEO C18, 5 μm particle size 300 μm × 5 mm) and separated in the analytical column (EASY-Spray C18 PepMap, 100 Å, 150 mm × 75 μm ID and 3 μm particle size) at a flow rate of 0.20 μL/min. Column temperature was set at 35 °C. Separation occurred using a multistep linear gradient to obtain 9% eluent B at 12 min, 36% eluent B at 102 min, and 99% eluent B at 107 min. The column was maintained at 99% eluent B for 6 min before re-equilibration at 2.5% eluent B. The mass spectrometer was operated in the positive ion mode, with an *m/z* range from 375 to 1600 with 70k resolution (Full MS), a spray voltage of 1.9 kV, and a transfer capillary temperature of 275 °C. Tandem MS (MS/MS) data were acquired using a data-dependent method with dynamic exclusion of 16.0 s at a 17,500 resolution. The top 10 most intense ions were selected for higher energy collisional dissociation (HCD), using 28% normalized collision energy (nce), and an isolation window of 1.4 m/z. Raw data were processed using MaxQuant v2.4.7.0 with the reviewed human proteome database from UniProt (20408 entries; accessed on 2023/07/07). In addition to the default settings for Orbitrap instrument and label-free quantification, the following settings were applied: trypsin/P as enzyme, three missed cleavages, fixed carbamidomethyl cysteine modification, and the variable modifications methionine oxidation, protein *N*-terminal acetylation, and serine/threonine/tyrosine phosphorylation, and match between runs enabled. Using the Perseus framework v2.0.11.0, only reverse phosphoproteins identified by site were included, while contaminants were excluded. Only phosphoproteins with a minimum of three valid values in at least one group were considered. Missing values were imputed based on a normal distribution. Phosphomatics was subsequently used for phosphosite mapping, functional annotation, and visualization of kinase-substrate networks. Differentially phosphorylated proteins were functionally annotated using KEGG and Reactome pathway enrichment, and kinase-substrate relationships were inferred via Kinase-Substrate Enrichment Analysis (KSEA).

### T-cell Activation and Proliferation Assays

Human peripheral T cells were isolated from buffy coats of healthy donors obtained from the Immunochemotherapy Department of Centro Hospitalar São João (CHSJ), Porto, Portugal. All procedures were conducted with informed consent and approved by the institutional ethics committee (Protocol reference 260/11). T cells were enriched using the RosetteSep™ Human T Cell Enrichment Cocktail (StemCell Technologies), following the manufacturer’s protocol. Briefly, buffy coats were centrifuged at 1200 × g for 30 minutes at room temperature (acceleration 5, no brake) to isolate peripheral blood mononuclear cells (PBMCs). PBMCs were incubated with RosetteSep reagent, diluted 1:1 in PBS containing 2% fetal bovine serum (FBS), and layered over Histopaque-1077 (Sigma-Aldrich) for density gradient centrifugation (30 minutes, 1200 × g, RT, no acceleration or brake). The T cell–enriched layer was collected and washed three times with PBS. Isolated T cells were cultured in RPMI 1640 GlutaMAX™ medium supplemented with 10% FBS, 1% penicillin/streptomycin, and 100 U/mL interleukin-2 (PeproTech, 200-02) and activated for 3 days using plate-bound anti-CD3 and anti-CD28 antibodies (**Table S5**). For proliferation tracking, T cells were labeled with CellTrace™ CFSE dye (Thermo Fisher Scientific). Cells were resuspended in PBS and incubated with CFSE (1□µL dye in 18□µL anhydrous DMSO) for 20 minutes at 37□°C. Labeling was quenched by adding complete medium, and cells were washed and resuspended for co-culture. Labeled T cells were co-cultured with glycoengineered SW620 cells (with or without *CD276* knockdown) at a tumor- to-T cell ratio of 1:5, in 24-well plates for 5 days at 37□°C and 5% CO₂. At the endpoint, conditioned media were collected for cytokine analysis, and cells were harvested by incubation with cold PBS on ice for 20 minutes. After washing, cells were stained with Fixable Viability Stain 780 for 15 minutes at 4□°C in the dark, followed by surface marker staining for CD3, CD4, CD8, CD25, and CD69 (**Table S5**) for 40 minutes at 4□°C. Flow cytometry analysis was performed on a NAVIOS flow cytometer (Beckman Coulter). Data were analyzed using Kaluza C software (v1.2) to assess viability, activation marker expression, mean fluorescence intensity (MFI), and the percentage of positive cells across relevant T cell subsets.

### Cytokine Evaluation

Conditioned media from CRC cell–T cell co-cultures were collected after 120 hours of direct contact. IL-1β, IL-2, IL-4, IL-5, IL-6, IL-8, IL-10, IL-12(p70), IL-13, IL-17A, IL-23, IFN-γ, TNF-α, and GM-CSF levels were quantified using a Luminex-based human multiplex cytokine assay, outsourced to EVE Technologies, according to their standard protocols.

### Tissue-Level Analysis of CD276 and Glycans by IHC

FFPE sections of colorectal tumors and healthy tissues were analyzed by IHC for CD276, Tn, and sTn antigens, as previously described by Peixoto *et al.* [51]. Antibodies, lectins, and detection reagents used are listed in **Table S5**. CD276 and anti-sTn antibodies were detected using the Novolink™ Polymer Detection System (Leica Biosystems) according to the manufacturer’s instructions. For Tn antigen detection, biotinylated VVA was used, followed by streptavidin– horseradish peroxidase (HRP) conjugate (Thermo Fisher Scientific, ready-to-use) and chromogenic development with ImmPACT® DAB Peroxidase Substrate (Vector Laboratories, 1:1000 dilution, 5 minutes at room temperature). Negative controls included tissue sections incubated with isotype-matched control antibodies and processed with omission of the primary antibody or lectin to assess background and non-specific staining. Sialidase digestion (α-neuraminidase treatment) was performed on selected sections before sTn staining to confirm the specificity of sialylated glycan detection; loss of signal upon treatment validated sialic acid–dependent binding. Positive controls included colorectal tumor tissues previously confirmed to express CD276, Tn, or sTn, as well as *C1GALT1*-knockout CRC cell–derived cells, known to exhibit high levels of exposed GalNAc residues for VVA specificity confirmation. Staining was semi-quantitatively evaluated based on both staining intensity (graded 0–3) and extent of positive staining across the tissue section (0–100%). A composite immunoreactivity score was derived by combining intensity and extension to reflect overall expression levels. All scoring was independently assessed by two trained observers and validated by an experienced pathologist to ensure consistency and biological relevance. All slides were imaged using a Motic BA310E microscope and analyzed with Motic Images Plus 3.0 software (Motic).

### Double Staining Immunofluorescence

A subset of FFPE tissue sections previously confirmed to be positive for Tn and CD276 was selected for double immunofluorescence staining to assess the colocalization of both epitopes. Briefly, tissue sections were deparaffinized, rehydrated, and subjected to heat-induced antigen retrieval using 1□mM EDTA buffer (pH□8.0; VWR). Tn antigens were detected using FITC-labeled VVA at a concentration of 40□μg/mL, incubated for 2□h at room temperature. CD276 was detected using an unlabeled rabbit polyclonal anti-CD276 antibody (Thermo Fisher Scientific; 1:250, 1□h at room temperature), followed by an Alexa Fluor 594–conjugated anti-rabbit secondary antibody (30□min at room temperature, in the dark). Controls were consistent with those used for immunohistochemistry, including isotype-matched antibodies, omission of primary reagents. Additionally, single-staining controls were included to exclude bleed-through or spectral overlap between fluorescence channels and to ensure specific signal attribution for each target. Fluorescence images were acquired using a Leica DMI6000 FFW microscope and analyzed with LAS X software (Leica Microsystems).

### Proximity Ligation Assay

The PLA was used to detect instances of close spatial proximity (<30□nm) between CD276 and sTn antigens in colorectal tumor and healthy tissue sections. FFPE tissues previously demonstrating co-localization of CD276 and sTn by immunohistochemistry, including normal colon samples, were selected for analysis. Tissue sections were deparaffinized, rehydrated, and subjected to antigen retrieval in boiling 1□mM EDTA buffer (pH 8.0; VWR) for 20 minutes. Non-specific binding was blocked using Blocking Solution (Sigma-Aldrich) for 1 hour at 37□°C. Primary antibodies against CD276 and sTn (anti-TAG-72) were applied and incubated overnight at 4□°C under the same conditions used for immunohistochemistry. PLA signal ligation and rolling circle amplification were performed using the Duolink® In Situ PLA kit (Sigma-Aldrich) according to the manufacturer’s instructions. Sections were then counterstained with DAPI and mounted using an aqueous mounting medium (Bio-Optica). To confirm signal specificity and exclude background noise, multiple controls were included. Tissue sections lacking expression of either CD276 or sTn, as determined by prior immunohistochemistry, were processed in parallel as negative controls. Additional negative controls included sections incubated with only one of the primary antibodies or with both primary antibodies omitted, to assess potential non-specific amplification or probe cross-reactivity. Furthermore, selected sections were pre-treated with α-neuraminidase (sialidase) prior to PLA staining to enzymatically remove sialic acids and confirm the sialylation dependence of the sTn signal. Positive controls consisted of colorectal tumor tissues previously validated for co-expression and spatial proximity of CD276 and sTn antigens by immunohistochemistry and immunofluorescence. Fluorescence images were acquired on a Leica DMI6000 FFW microscope and analyzed using LAS X software (Leica Microsystems).

### Statistical analysis

All statistical analyses were performed using GraphPad Prism (v9; Dotmatics), SPSS for MacOS (version 27; IBM), and R software (version 4.2.1; R Foundation for Statistical Computing). For *in vitro* and *ex vivo* experiments, including glycogenes and *CD276* gene expression (RT-qPCR), cell invasion and proliferation assays, cytokine production, T-cell proliferation, and immune phenotype characterization, statistical comparisons were conducted using one-way ANOVA followed by Tukey’s post hoc multiple comparisons test. Data normality was assessed using the Shapiro–Wilk test, and when variables deviated from normality, non-parametric alternatives such as the Wilcoxon rank-sum test were applied. Chi-square tests were used to evaluate associations between categorical variables such as tumor stage, anatomical location, CD276 status, and CD276–Tn/sTn expression. Differences in CD276 expression and staining extent between metastatic and non-metastatic samples were assessed using a non-parametric t-test (Mann–Whitney U test). For TCGA-based transcriptomic analyses, including comparisons between epithelial-like and mesenchymal-like CRC subtypes, Mann–Whitney U test tests were used to evaluate differences in CD276 and CD276–Tn/sTn expression levels. Spearman’s rank correlation was used to assess associations between gene expression levels, and correlation matrices were visualized using the “corrplot” R package. To explore the prognostic value of CD276 and glycosyltransferase gene expression, univariate and multivariate Cox proportional hazards regression models were performed. Kaplan–Meier survival curves were generated for overall survival (OS) and progression-free survival (PFS), and differences between survival groups were tested using the log-rank test, as implemented in the “survival” and “survminer” R packages. Optimal expression cut-offs for survival stratification were identified using the surv_cutpoint() function. Unless otherwise stated, statistical significance was considered at *p*□<□0.05.

## Acknowledgments

The authors wish to acknowledge the Portuguese Foundation for Science and Technology (FCT) for PhD grants SFRH/BD/142479/2018 (JS), 2020.09384.BD (https://doi.org/10.54499/2020.09384.BD) (DF); 2022.12980.BD (https://doi.org/10.54499/2022.12980.BD) (AM), SFRH/BD/146500/2019 (https://doi.org/10.54499/SFRH/BD/146500/2019) and COVID/BD/153652/2024 (https://doi.org/10.54499/COVID/BD/153652/2024) (MRS), 2020.09394.BD (https://doi.org/10.54499/2020.09394.BD) (SC), 2020.08708.BD (https://doi.org/10.54499/2020.08708.BD) (RF), 2024.04043.BDANA (EF), 2024.00895.BD (BMS), Principal researcher contract 2022.08311.CEECIND (https://doi.org/10.54499/2022.08311.CEECIND/CP1744/CT0001) (JAF), assistant researcher contract 2018.00091.CEECINST (PP) and junior researcher contacts 2021.03835.CEECIND (https://doi.org/10.54499/2021.03835.CEECIND/CP1688/CT0001) (AB), 2023.06366.CEECIND/CP2864/CT0001 (https://doi.org/10.54499/2023.06366.CEECIND/CP2864/CT0001) (AP). FCT is co-financed by European Social Fund under Human Potential Operation Programme from National Strategic Reference Framework. The authors also acknowledge funding from the IPO-Porto research center (PEst-OE/SAU/UI0776/201, CI-IPOP-29-2016-2022, CI-IPOP-58-2016-2022), for CQUM (UID/QUI/00686/2020), and the LAQV research unit (Project UID/QUI/50006/2019 - Laboratório Associado para a Química Verde-Tecnologias e Processos Limpos) through the Portuguese Foundation for Science and Technology/Ministry for Education, Science, and Innovation funding. This article is also a result of the project NORTE-01-0145-FEDER-000012 and “The Porto Comprehensive Cancer Center Raquel Seruca” with the reference NORTE-01-0145-FEDER-072678 - Consórcio PORTO.CCC— Porto.Comprehensive Cancer Center Raquel Seruca, supported by NORTE 2020, under the PORTUGAL 2020 Partnership Agreement, through the European Regional Development Fund. This work was also financed by FEDER through the COMPETE 2020 - Operational Programme for Competitiveness and Internationalisation (POCI-01-0145-FEDER-007274 for i3S unit). Finally, the authors also acknowledge the ICBAS PhD Program in Biomedical Sciences.

## Data Availability Statement

All data generated or analyzed during this study are included in this published article and its supplementary information files. Glycomics data have been deposited at GlycoPost (accession number GPST000606, project: GPST000606.0) [54].

## Abbreviations

ATCC: American Type Culture Collection
CHSJ: Centro Hospitalar São João
CMS: Consensus Molecular Subtypes
CORA: Cellular O-Glycome Reporter/Amplification
CRC: Colorectal Cancer
dsT: Disialyl-T
DTT: Dithiothreitol
EMT: Epithelial-Mesenchymal Transition
FBS: Fetal Bovine Serum
FC: Flow Cytometry
FDR: False Discovery Rate
FFPE: Formalin-Fixed Paraffin-Embedded
FMO: Fluorescence Minus One
HCD: Higher-energy Collisional Dissociation
HR: Hazard Ratio
ICIs: Immune Checkpoint Inhibitors
IDAA: Indel Detection by Amplicon Analysis
IF: Immunofluorescence
IgC: Immunoglobulin Constant Domain
IgV: Immunoglobulin Variable Domain
IHC: Immunohistochemistry
IP: Immunoprecipitation
IPO Porto: Portuguese Institute of Oncology of Porto
KO: Knockout
KSEA: Kinase-Substrate Enrichment Analysis
MFI: Mean Fluorescence Intensity
MS: Mass Spectrometry
MSI: Microsatellite Instability
MSS: Microsatellite Stability
Neu: Neuraminidase
NSAF: Normalized Spectral Abundance Factor
OS: Overall Survival
PBMCs: Peripheral Blood Mononuclear Cells
PCA: Principal Component Analysis
PFS: Progression-free Survival
PLA: Proximity Ligation Assay
PNA: Peanut agglutinin
ppGalNAc-Ts: Polypeptide N-acetylgalactosaminyltransferases
PRIDE: PRoteomics IDEntifications Database
RNA-seq: RNA Sequencing
RT-qPCR: Quantitative Reverse Transcription Polymerase Chain Reaction
siRNAs: Short Interfering RNA or Silencing RNA
sLe: Sialyl Lewis
SSC-A: Side Scatter Area
sT: Sialyl-T
α(2,3)sialylTs: α2,3-sialyltransferases
sTn: Sialyl Tn
TCGA: The Cancer Genome Atlas
TMB: Tumour Mutational Burden
VVA: Vicia Villosa Agglutinin
WB: Western Blot
WT: Wild Type
α(3)FucTs: α3-fucosyltransferases.

## Author Contributions

Conceptualization: JAF; Methodology: JS, DF, MG, AM, MRS, AB, PP, CP, FA, AP, LLS, JAF; Software: JS, MRS, AB, AMNS, JAF; Validation: JS, DF, MG, AM, MRS, AB, PP, LPA, AMNS, CP, FA, AP, LLS, JAF; Investigation: JS, DF, MG, AM, MRS, AB, PP, SC, RF, MM, EF, BMS, JAF; Resources: JS, DF, AM, MRS, AB, LPA, AMNS, CP, FA, AP, LLS, JAF; Data Curation: JS, DF, MG, AM, MRS, AB, SC, LPA, AMNS, CP, FA, AP, LLS, JAF; Writing-Original Draft: JAF; Writing-Review & Editing: all authors; Visualization: JS, DF, MG, AM, MRS, AB, JAF; Supervision: JAF; Project Administration: JAF; Funding Acquisition: AP, LLS and JAF.

