## Supporting Figures for "CD276 Immature Glycosylation Drives Colorectal Cancer Aggressiveness and T-cell Mediated Immune Escape"

^1^Research Center of IPO-Porto (CI-IPOP) / RISE@CI-IPOP (Health Research Network), Portuguese Oncology Institute of Porto (IPO-Porto) / Porto Comprehensive Cancer Center (P.ccc) Raquel Seruca, Porto, Portugal; ^2^School of Medicine and Biomedical Sciences (ICBAS), University of Porto, Porto, Portugal; ^3^LAQV-REQUIMTE & Department of Chemistry, University of Aveiro, Campus Universitário de Santiago, Aveiro, Portugal; ^4^i3S – Instituto de Investigação e Inovação em Saúde, Universidade do Porto, Porto, Portugal; ^5^LAQV-REQUIMTE, Department of Chemistry and Biochemistry, Faculty of Sciences, University of Porto, Porto, Portugal; ^6^Pathology Department, Portuguese Oncology Institute of Porto, Porto, Portugal; ^7^Immunology Department, Portuguese Oncology Institute of Porto, Porto, Portugal; ^8^School of Medicine and Biomedical Sciences of University Fernando Pessoa, Porto, Portugal; ^9^Department of Surgical Oncology, Portuguese Oncology Institute of Porto, Porto, Portugal.

**Corresponding author:**

José Alexandre Ferreira

Experimental Pathology and Therapeutics Group,

Research Centre, Portuguese Oncology Institute of Porto,

R. Dr. António Bernardino de Almeida 4200-072 Porto,

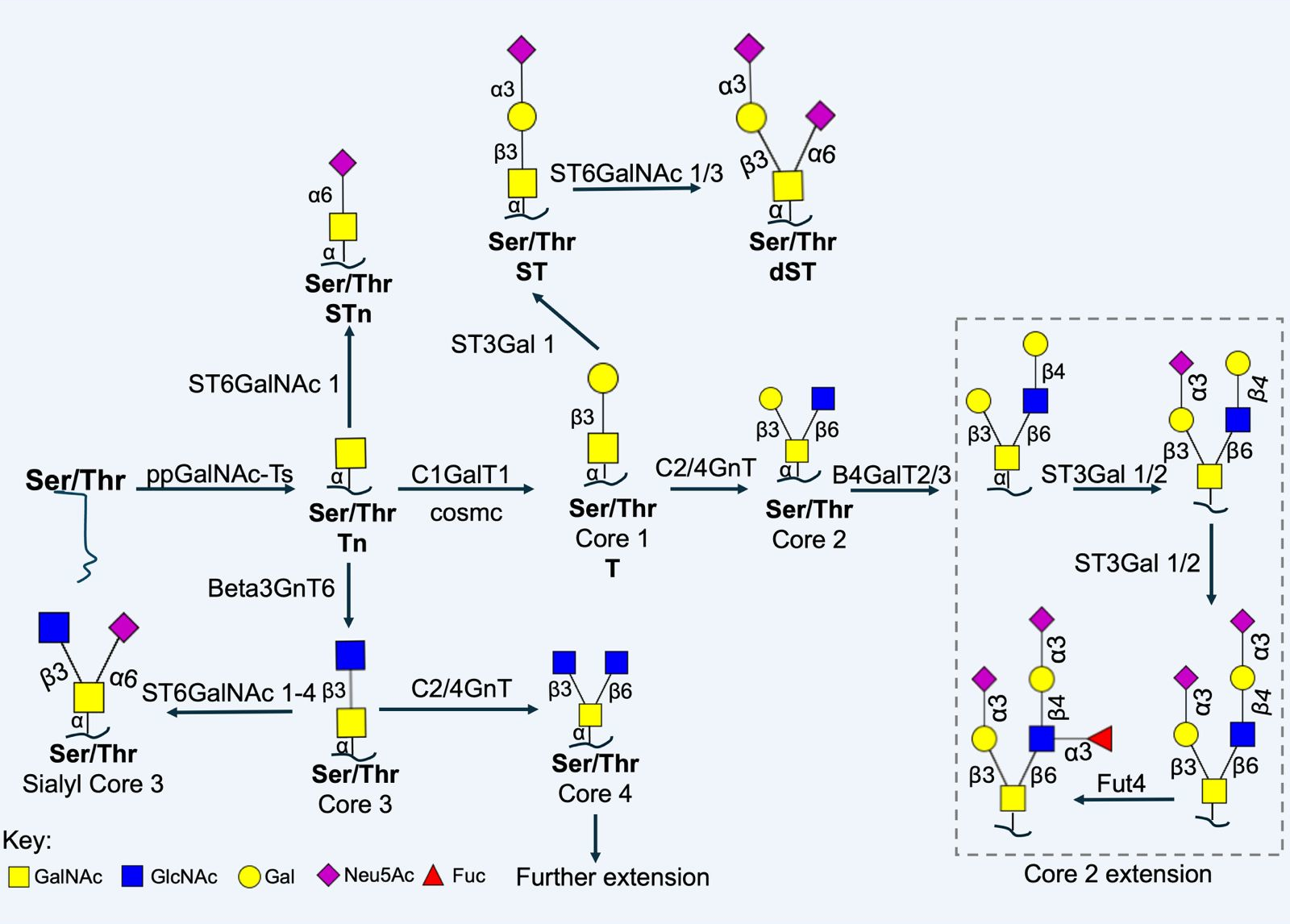

**Figure S1. Schematic representation of *O*-glycans biosynthesis.** The addition of a GalNAc to a Ser or Thr on a growing peptide chain is prompted by the action of polypeptide N-acetylgalactosamine transferases (ppGalNAc-Ts; a family of 20 enzymes), generating the Tn antigen, which can be extended by the action of C1GalT1 and its chaperone cosmc, forming the Thomsen-Friedenreich or T antigen (core 1). Alternatively, Tn can be sialylated by α2,6-sialyltransferases (ST6GalNAc; a family of 6 enzymes – ST6GalNAc 1, ST6GalNAc 2, ST6GalNAc 3, ST6GalNAc 4, ST6GalNAc 5, ST6GalNAc 6), forming the sialyl-Tn (sTn) and stopping O-glycan extension. Alternatively, Tn can be extended by B3GNT6, forming core 3, which can be further extended into core 4 by GCNT3 or sialyled by ST6GalNAc 1-4. Additionally, T antigen can be sialylated by α2,3-sialyltransferases (ST3Gal; a family of 6 enzymes - ST3Gal 1, ST3Gal 2, ST3Gal 3, ST3Gal 4, ST3Gal 5, ST3Gal 6), forming the sialyl-T, and further sialylated by ST6GalNAc 1/3, generating di-sialyl-T. T antigen can also be elongated by the action of GCNT1/4, forming core 2. Finally, core 2 can be further extended into complex structures, such as sialyl lewis epitopes, by the action of ST3Gal-Ts and α3-fucosyltransferases (α3-Fut-Ts, a family of 6 enzymes – FUT 3, FUT 4, FUT 5, FUT 6, FUT 7, FUT 9)**.**

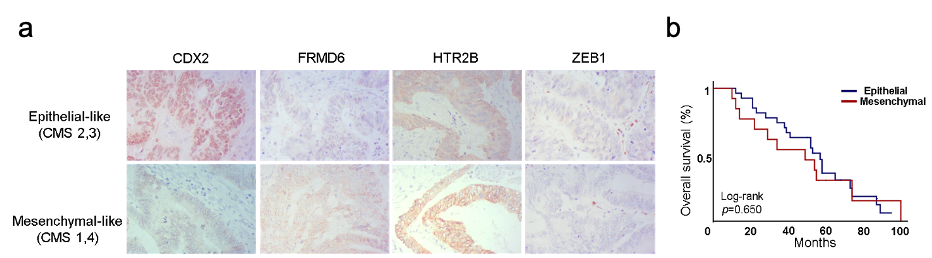

**Figure S2. Colorectal cancer differentiation state does not influence prognosis. A**. **Molecular characterisation of CRC tumours into two differentiation states.** Immunohistochemical analysis categorised CRC tumours into Epithelial-like tumours characterised by high expression of CDX2 and low levels of FRMD6 and HTR2B, while mesenchymal-like tumours show low expression of CDX2 and high expression of FRMD6 and HTR2B. ZEB1 is poorly expressed in both subtypes. **B.** **CRC overall survival based on differentiation states.** The Kaplan-Meier analysis does not show a significant association of differentiation states with overall survival.

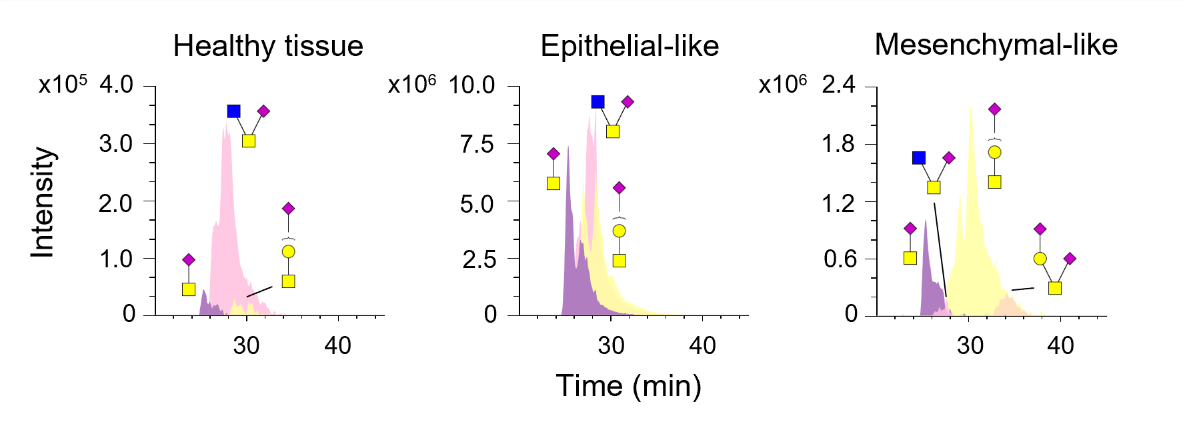

**Figure S3. *O-*glycan abundances change between healthy mucosa and colorectal tumours.** Healthy tissues are predominantly characterised by a high abundance of sialyl-core 3 (*pink*), while also presenting sTn (*purple*) and (sialyl-)T (*yellow* ??) antigens. Meanwhile, epithelial-like tumours present a higher abundance sTn and (sialyl-)T antigen compared to healthy tissues, although still presenting sialyl-core 3 structures. Finally, mesenchymal-like tumours are mainly characterised by mono- or di-sialylated core 1 structures (*orange*), while presenting lower sTn and sialylated core-3, compared to epithelial-like tumours and healthy tissues.

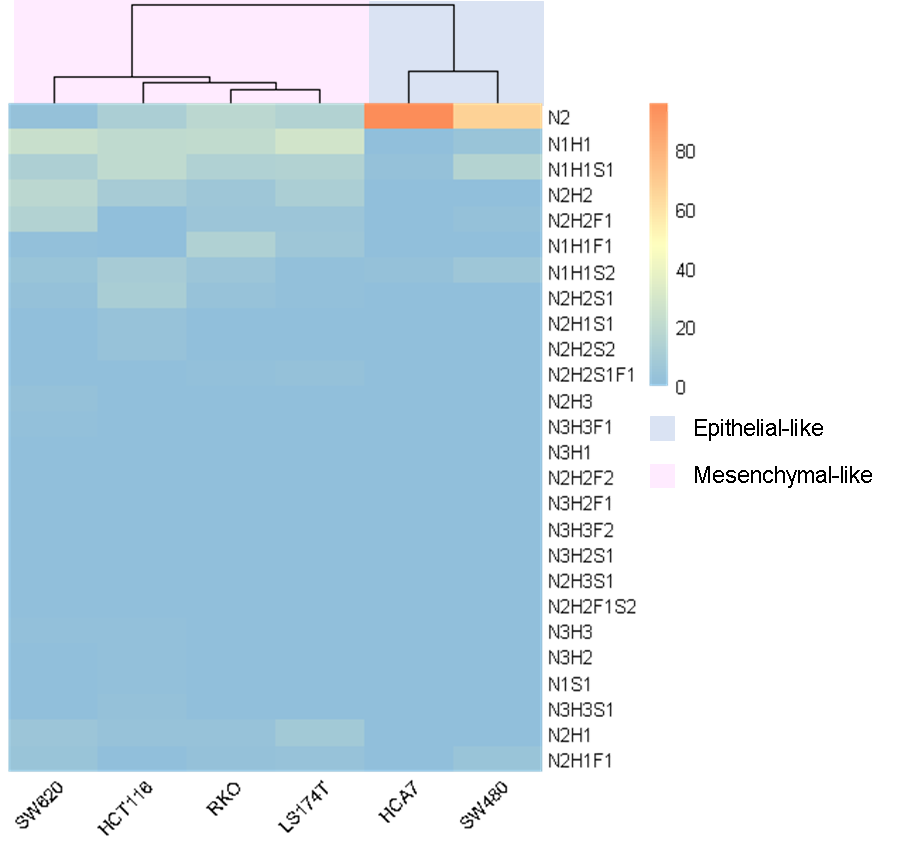

**Figure S4. Glycome profiling of colorectal cancer cell lines.** Colorectal cancer cell lines mainly show simple short *O*-glycosylation, exhibiting similarities with CRC tumours. According to colorectal tumours similarities, cell lines were clustered into two differentiation states commonly defining CRC, namely epithelial-like and mesenchymal-like phenotypes. HCA7 and SW480 cell lines presented high abundances of core 3 structures, while SW620, HCT116, RKO, and LS174T were mainly characterised by core 1, mono-sialylated core 1, and elongated core 2. Accordingly, HCA7 and SW480 cell lines may be classified as epithelial-like, while SW620, HCT116, RKO, and LS174T are more mesenchymal-like cells.

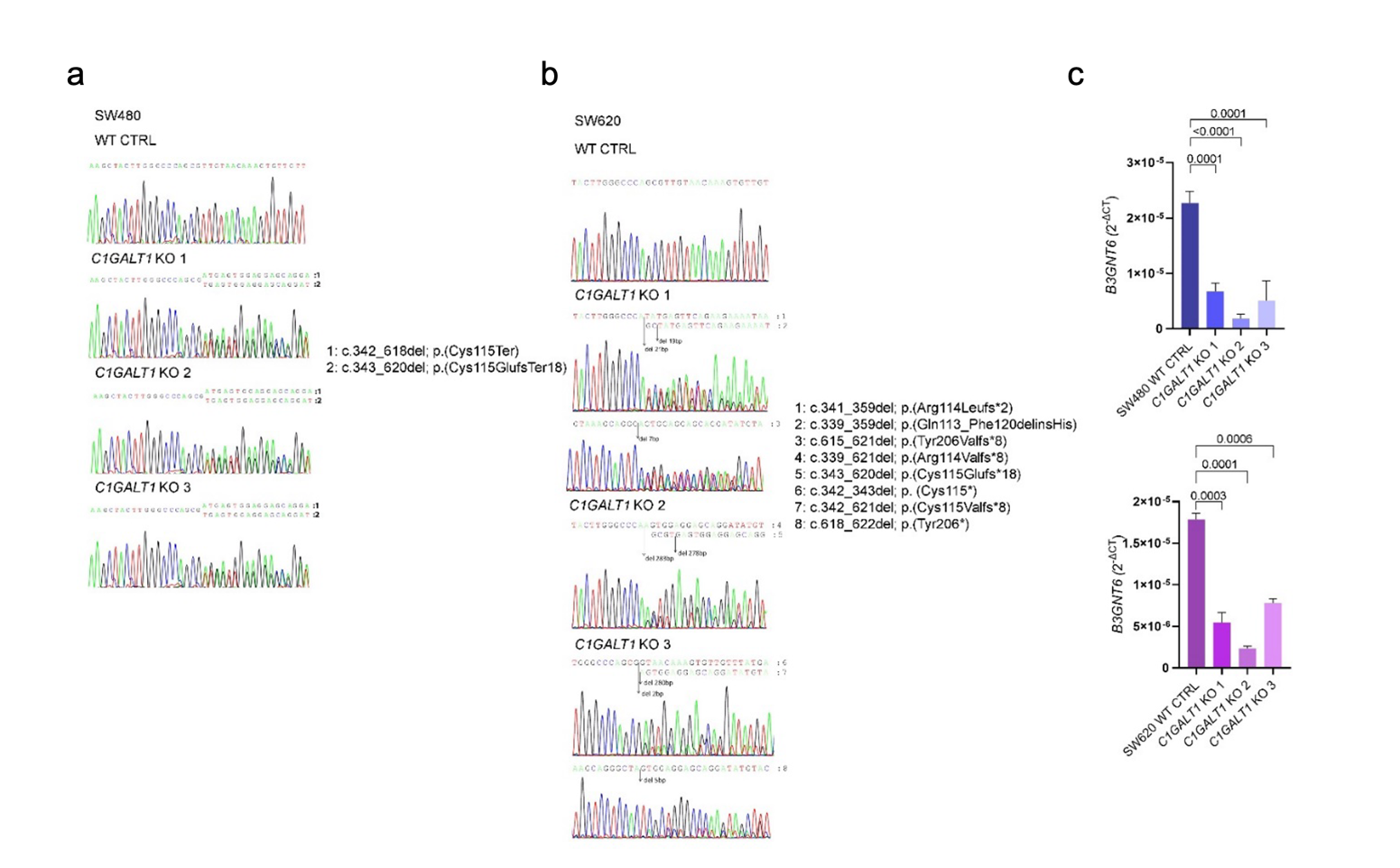

**Figure S5. SW480 and SW620 colorectal cancer cell lines were successfully knocked out for the *C1GALT1* gene, subsequently downregulating *B3GNT6*. A. Sanger sequencing of SW480 *C1GALT1* knockout clones.** Cancer cell line models resembling glycosyltransferases phenotype associated with worse prognosis were generated by CRISPR/Cas9 technology. Sanger sequencing showed that SW480 *C1GALT1* KO clones are characterized by two different DNA sequences, with the following observed variants and predicted consequences: 1: c.342_618del; p.(Cys115Ter), leading to a 277bp deletion and introduction of a stop codon at Cys 115; 2: c.343_620del; p.(Cys115Glufs*18), leading to a 278bp deletion and replacement of Cys 115 by Glu, resulting in a frameshift introducing a stop codon 18 a.a ahead. **B. Sanger sequencing of SW620 *C1GALT1* knockout clones.** Sanger sequencing showed that SW620 *C1GALT1* KO clone 1 is characterized by three different DNA sequences, with the following observed variants and predicted consequences: 1: c.341_359del; p.(Arg114Leufs*2), leading to a 19bp deletion and the replacement of Arg 114 by Leu, resulting in a frameshift introducing a stop codon 2 a.a ahead; 2: c.339_359del; p.(Gln113_Phe120delinsHis), leading to a 21bp deletion and replacement of Gln 113 by Phe 120 and insertion of His; and 3: c.615_621del; p.(Tyr206Valfs*8), leading to a 7bp deletion and the replacement of Tyr 206 by Val, resulting in a frameshift introducing a stop codon 8 a.a ahead. SW620 *C1GALT1* KO clone 2 is characterized by two different DNA sequences, with the following observed variants and predicted consequences: 4: c.339_621del; p.(Arg114Valfs*8), leading to a 283bp deletion and the replacement of Arg 114 by Val, resulting in a frameshift introducing a stop codon 8 a.a ahead; and 5: c.343_620del; p.(Cys115Glufs*18), leading to a 278bp deletion and replacement of Cys 115 by Glu, resulting in a frameshift introducing a stop codon 18 a.a ahead. SW620 *C1GALT1* KO clone 3 is characterized by three different DNA sequences, with the following observed variants and predicted consequences: 6: c.342_343del; p. (Cys115*), leading to a 2bp deletion and introducing a stop codon at Cys 115; 7: c.342_621del; p.(Cys115Valfs*8), leading to a 280bp deletion and the replacement of of Cys 115 by Val, resulting in a frameshift introducing a stop codon 8 a.a ahead; and 8: c.618_622del; p.(Tyr206*), leading to a 5bp deletion and introducing a stop codon at Tyr 206. **C. *B3GNT6* gene expression evaluation in SW480 and SW620 *C1GALT1* KO cell models.** *B3GNT6* gene expression was significantly decreased in *C1GALT1* KO clones for both cell lines when compared to WT controls.

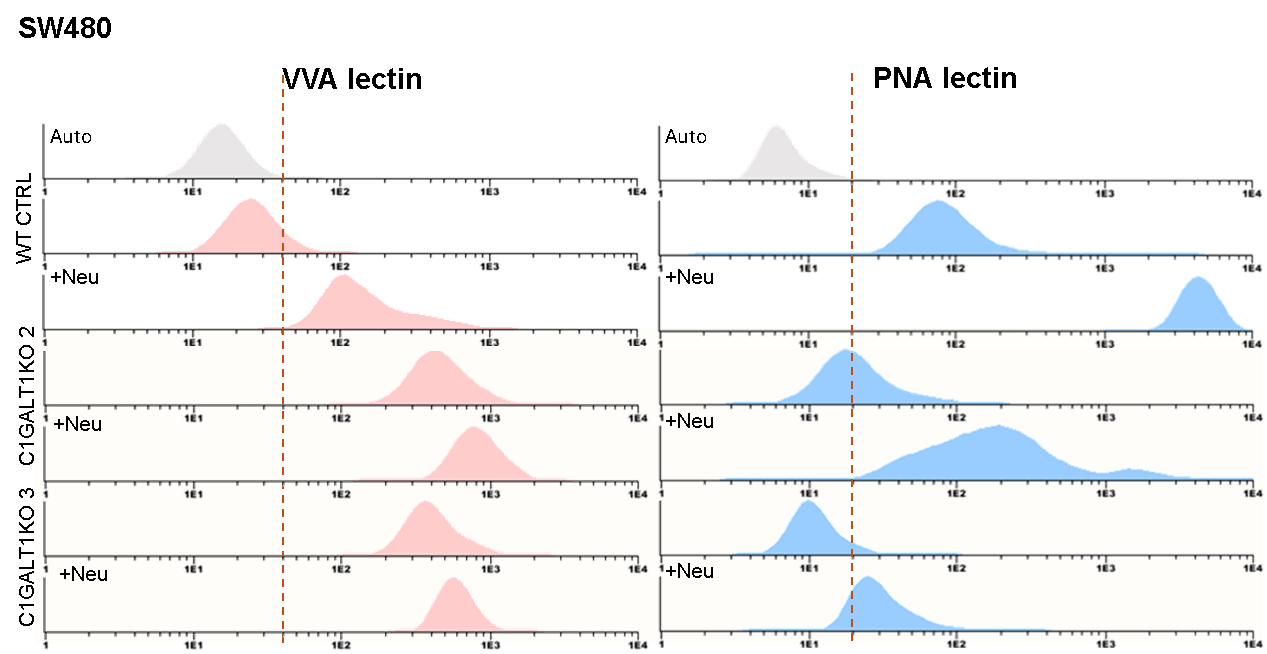

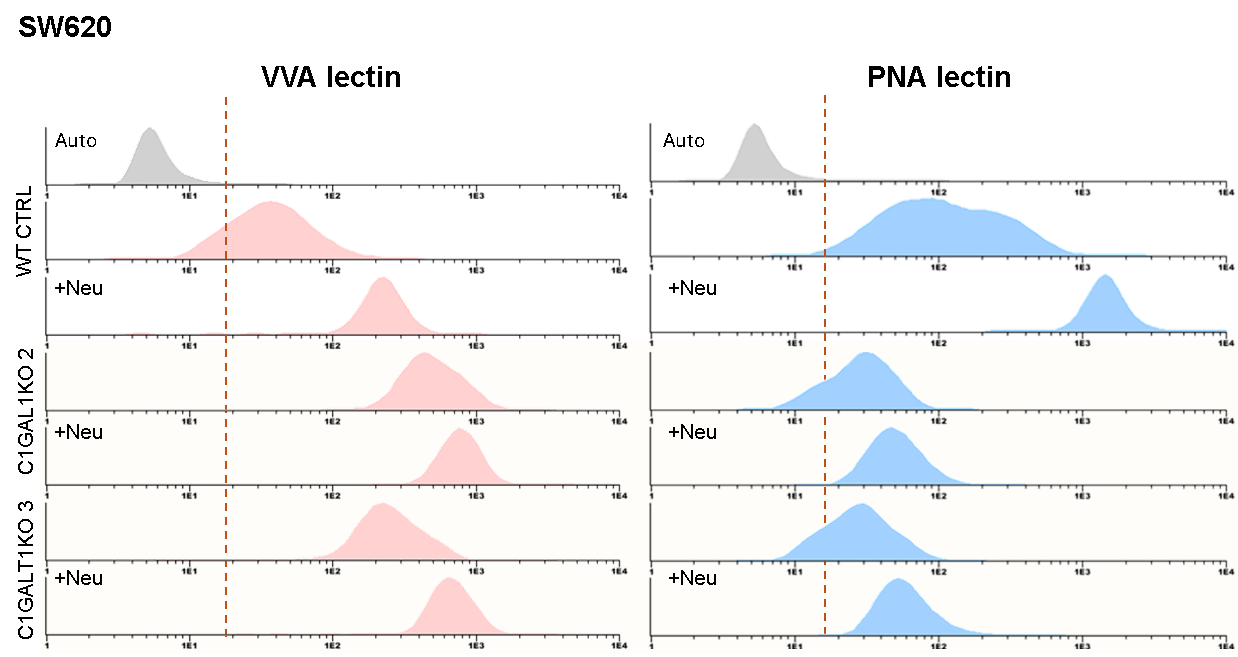

**Figure S6. Flow cytometry analysis of glycoengineered C1GALT1 KO SW480 and SW620 cell lines.** Flow cytometry using VVA lectin confirmed the accumulation of the Tn antigen on the cell surface of *C1GALT1* knockout cells (left panels, red), while PNA lectin confirmed the abrogation of *O*-glycan extension (right panels, blue). To distinguish between sialylated and non-sialylated *O*-glycans, cells were also treated with neuraminidase (Neu) prior to lectin staining. Neuraminidase treatment enhanced PNA binding, consistent with the presence of sialyl-T antigen (sT) in control cells and its loss in *C1GALT1*-deficient backgrounds. Similarly, VVA binding increased after desialylation, supporting the presence of sTn in the glycoengineered cells. Together, these results confirm the truncation of *O*-glycans to Tn and sTn structures following *C1GALT1* deletion.

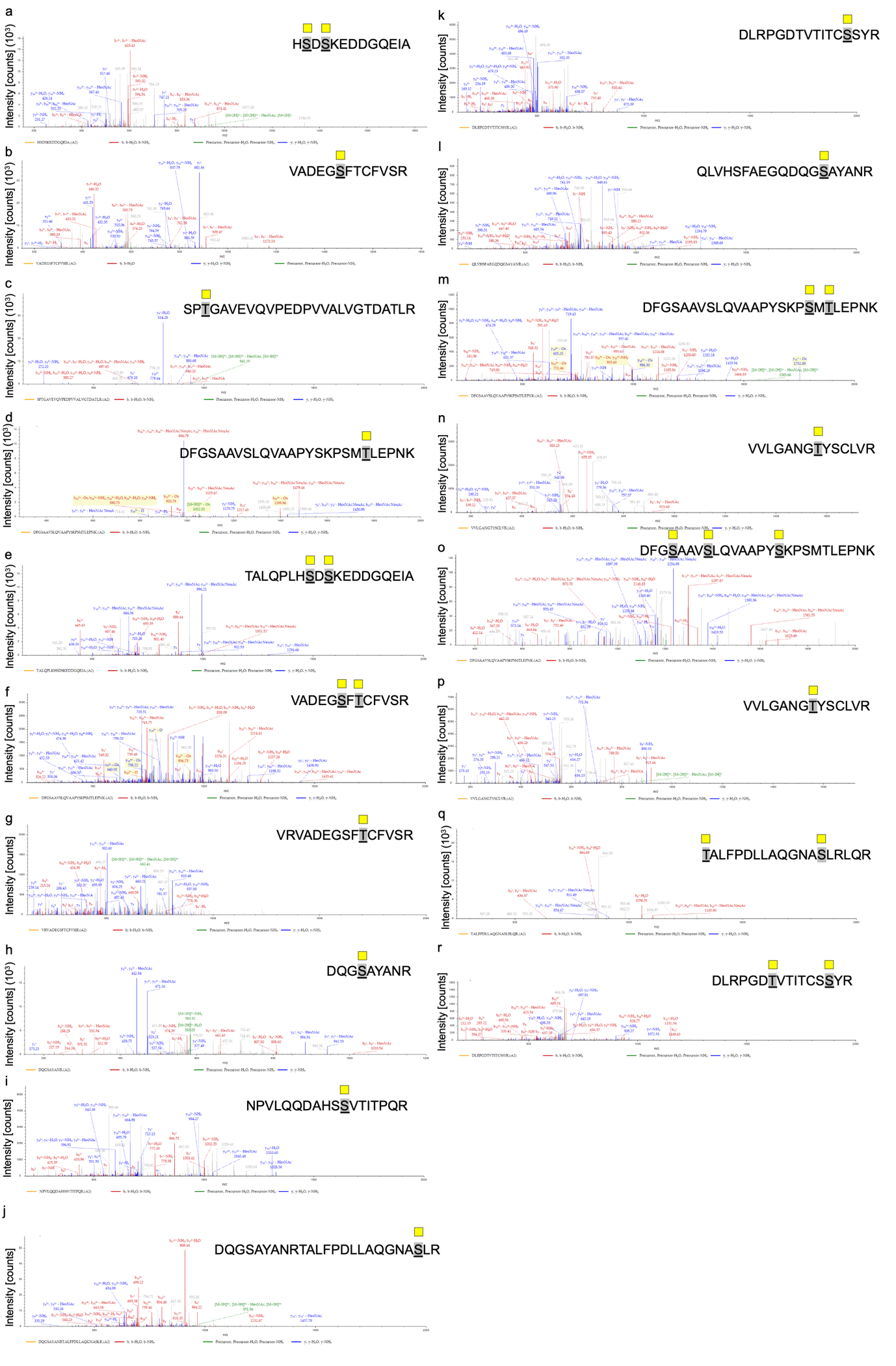

**Figure S7. CID-MS/MS spectra of CD276 glycopeptides bearing immature O‑glycosylation in CRC patient samples.** Representative collision‑induced dissociation (CID) MS/MS spectra of CD276‑derived glycopeptides were identified in CRC samples from a proteomics dataset (dataset identifier PXD002080). The spectra display characteristic oxonium ions, together with extensive b‑, y‑, and internal fragment ions typical of CID fragmentation, providing high sequence coverage while retaining the HexNAc (Tn antigen) on the peptide backbone (yellow square). Glycopeptides were confidently assigned and manually validated. These data offer direct evidence in tissue that CD276 carries immature O‑glycans in CRC. While the spectra support putative glycosite assignments, the inherent limitations of CID for precise site localization underscore the need for orthogonal validation using dedicated fragmentation methods. Further details on peptide fragments are available in **Supporting Data File 2**.

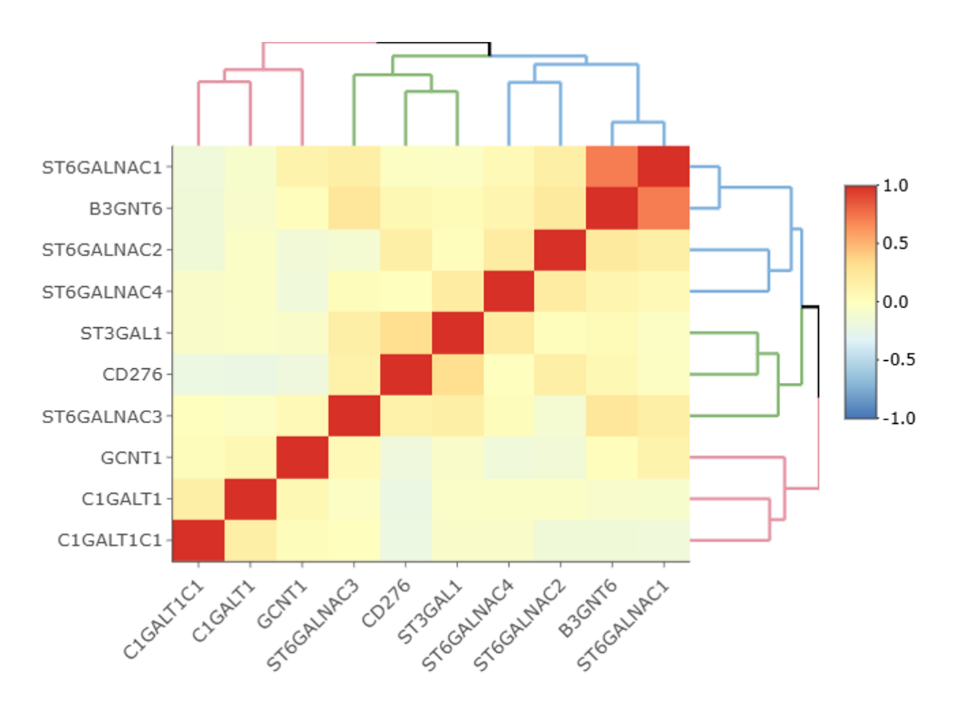

**Figure S8. Correlogram of selected glycogenes and *CD276* expression in colorectal cancer (TCGA dataset).** Pearson correlation coefficients are shown from negative (blue) to positive (red), with hierarchical clustering revealing distinct gene clusters. *B3GNT6* and *ST6GALNAC1*, typically abundant in healthy colon mucosa, cluster together, reflecting their coordinated role in core 3 biosynthesis and sialylation. *CD276* clusters with *ST3GAL1* and *ST6GALNAC* family members, consistent with enhanced sialylation of immature O‑glycans in tumours. Notably, *C1GALT1* and *C1GALT1C1*, key enzymes for core 1 elongation, show negative correlations with *CD276*, supporting a relation between immature glycosylation and high *CD276* expression.

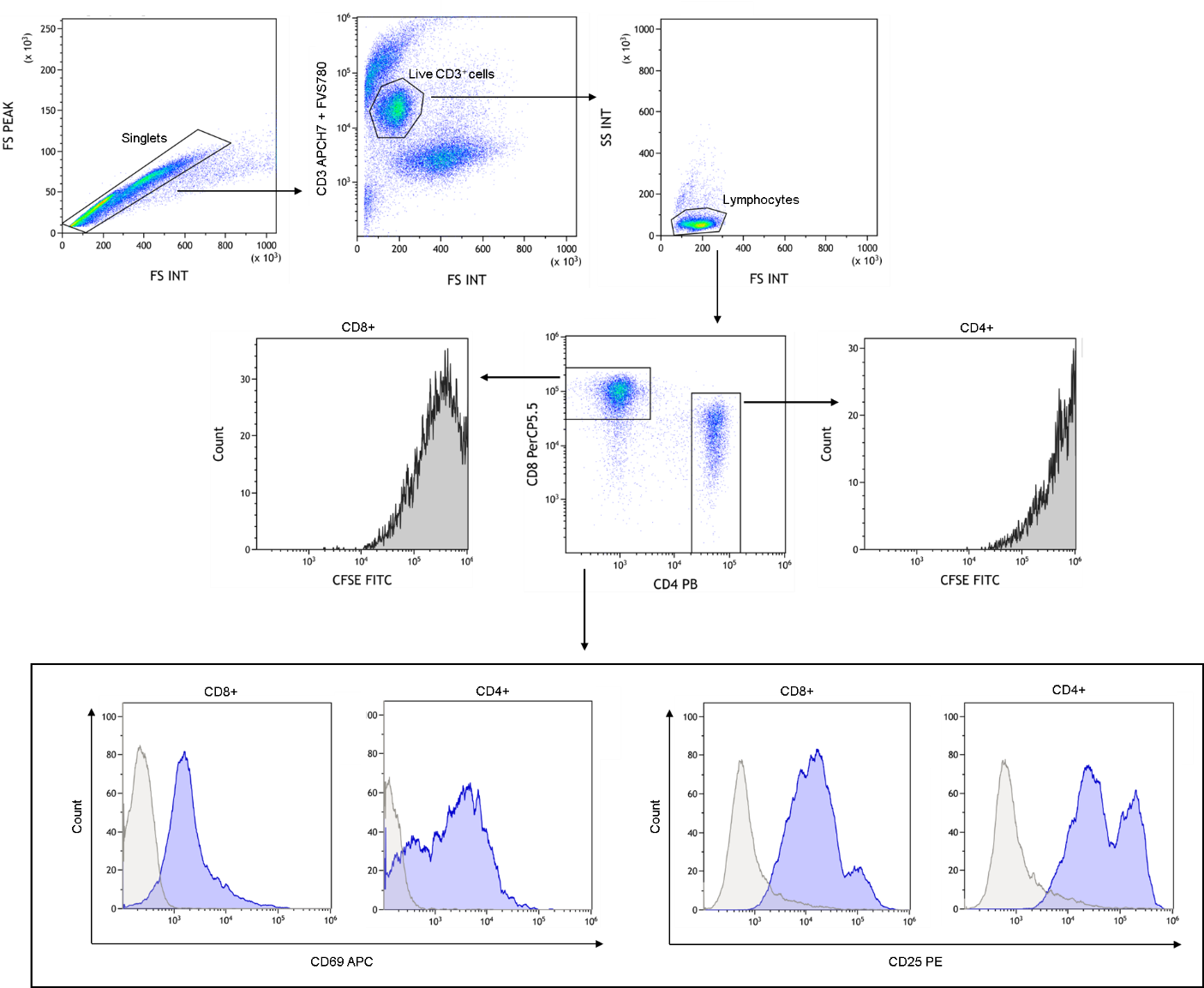

**Figure S9. Flow cytometric gating strategy to evaluate T cell activation and proliferation in co-culture settings with tumour cells.** Singlets were selected according to forward scatter-height (FSC-H) versus forward scatter-area (FSC-A). Then, live CD3 positive cells were isolated according to Fixable Viability Stain 780 (FVS780), CD3 expression, size (FSC-A) and granularity characteristics (side scatter-area (SSC-A)). Gated on T cells (CD3+ lymphocytes), CD4+ and CD8+ T cell subsets were identified. For each subset (CD4+ and CD8+), histograms of CFSE expression allowed the evaluation of T cell proliferation by CFSE dilution over each cell division during the 5 days of co-culturing. To investigate the impact of CD276-Tn in T cell activation, early and late activation markers (CD69 and CD25, respectively) were evaluated on CD4+ and CD8+ T cells. Fluorescence minus one (FMO) controls were used to set the gates for CD25 and CD69 positive populations.

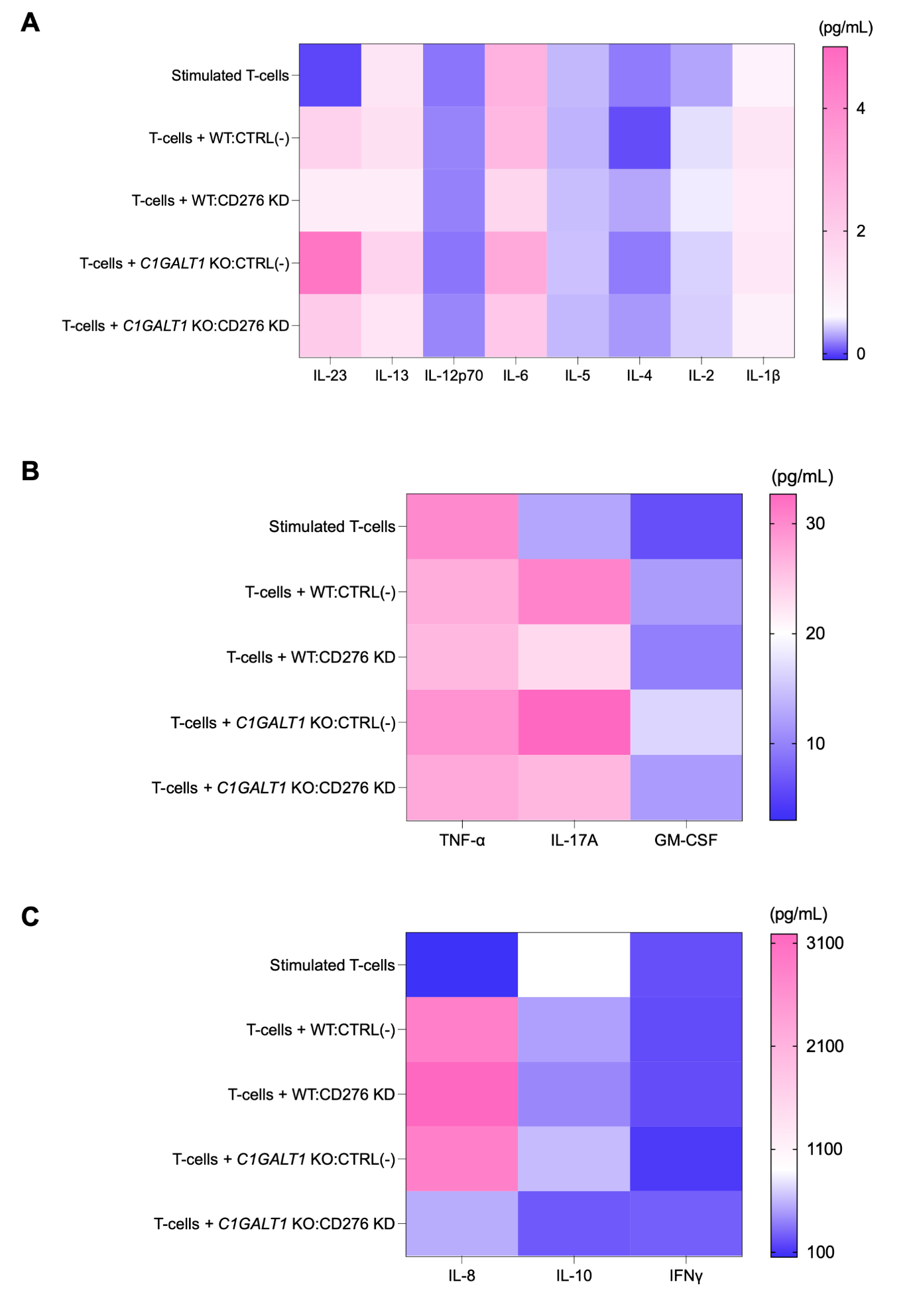

**Figure S10. Cytokine secretion profiles of T cells co-cultured with *C1GALT1*-deficient CRC cells, with or without *CD276* knockdown.** Heatmaps depicting the levels of cytokines secreted by stimulated human T cells co-cultured with either wild-type (WT) or *C1GALT1* knockout (KO) SW620 cells, in the presence or absence of *CD276* knockdown (*CD276* KD). **A) Immature *O*-glycosylation in cancer cells disrupts the cytokine environment that supports T cell activation and polarization.** Analysis of key pro- and anti-inflammatory cytokines (IL-23, IL-13, IL12p70, IL-6, IL-5, IL-1β) reveals that co-cultures with *C1GALT1* KO cells increased production of most cytokines, relative to WT controls, except for IL12p70, whose levels were reduced. Upon *CD276* knockdown, cytokine secretion was partially restored, notably for IL-23, IL17A and IL13 and IL-12p70. **B) The loss of *O*-glycan extension increases tumor-elicited inflammation, a defect that is partly reversed by targeting CD276.** Pro-inflammatory cytokines (TNF-α and IL-17A) are increased in *C1GALT1* KO co-cultures compared to WT, while *CD276* knockdown in this background restores TNF-α and IL-17A levels to those comparable with or lower than WT controls. **C) Aberrant glycosylation of CD276 reshapes the cytokine landscape, dampening both innate and adaptive immune outputs.** Cytokines associated with innate and immune regulatory responses, such as IL-8 and IL-10, show a similar trend, with marked suppression in the co-cultures with *C1GALT1* KO following CD276 knockdown. By contrast, IFN-γ was markedly reduced in the co-cultures with *C1GALT1* KO cells and restored after CD276 silencing. Together, these results suggest that loss of core 1 *O*-glycosylation in cancer cells broadly suppresses effective T cell cytokine responses, an effect partially mediated by CD276. Its silencing restores specific immune functions in vitro, implicating immaturely glycosylated CD276 as a potential key modulator of T cell activity.
