## Supporting Data Files 2 for "CD276 Immature Glycosylation Drives Colorectal Cancer Aggressiveness and T-cell Mediated Immune Escape"

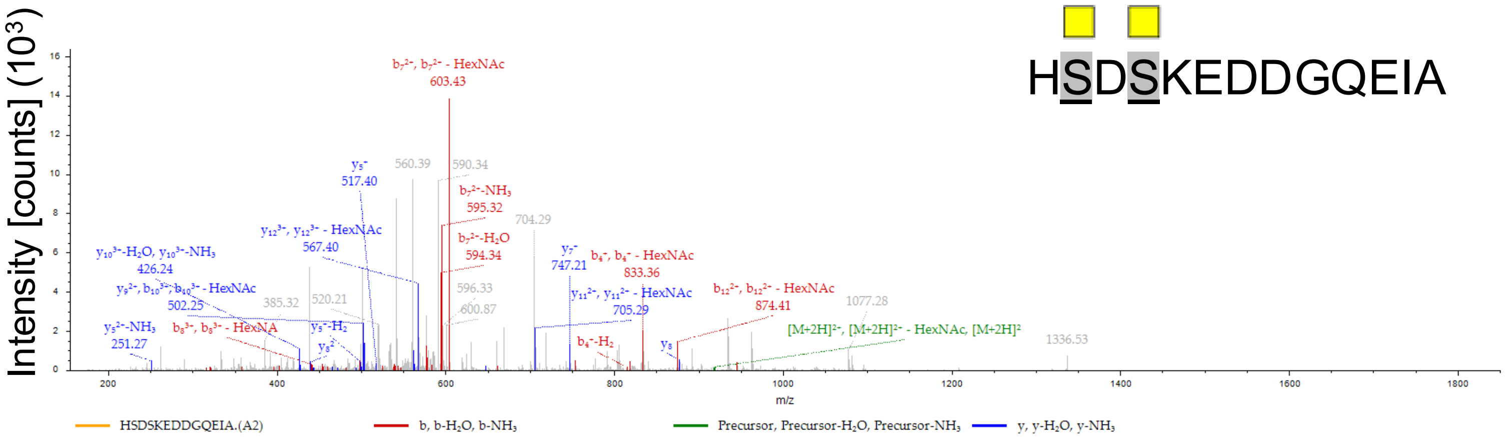


HSDSKEDDGQEIA

| #1 | $b^+$ | $b^{2+}$ | $b^{3+}$ | Seq. | $y^+$ | $y^{2+}$ | $y^{3+}$ | #2 |
|---|---|---|---|---|---|---|---|---|
| 1 | 138.06619 | 69.53673 | 46.69358 | H | | | | 13 |
| 2 | 428.17759 | 214.59243 | 143.39738 | S-HexNAc | 1699.70284 | 850.35506 | 567.23913 | 12 |
| 3 | 543.20453 | 272.10590 | 181.73970 | D | 1409.59144 | 705.29936 | 470.53533 | 11 |
| 4 | 833.31593 | 417.16161 | 278.44350 | S-HexNAc | 1294.56450 | 647.78589 | 432.19302 | 10 |
| 5 | 961.41090 | 481.20909 | 321.14182 | K | 1004.45309 | 502.73019 | 335.48922 | 9 |
| 6 | 1090.45349 | 545.73038 | 364.15601 | E | 876.35813 | 438.68270 | 292.79090 | 8 |
| 7 | 1205.48043 | 603.24385 | 402.49833 | D | 747.31554 | 374.16141 | 249.77670 | 7 |
| 8 | 1320.50738 | 660.75733 | 440.84064 | D | 632.28860 | 316.64794 | 211.43438 | 6 |
| 9 | 1377.52884 | 689.26806 | 459.84780 | G | 517.26165 | 259.13446 | 173.09207 | 5 |
| 10 | 1505.58742 | 753.29735 | 502.53399 | Q | 460.24019 | 230.62373 | 154.08491 | 4 |
| 11 | 1634.63001 | 817.81864 | 545.54819 | E | 332.18161 | 166.59444 | 111.39872 | 3 |
| 12 | 1747.71407 | 874.36068 | 583.24288 | I | 203.13902 | 102.07315 | 68.38452 | 2 |
| 13 | | | | A | 90.05496 | 45.53112 | 30.68984 | 1 |

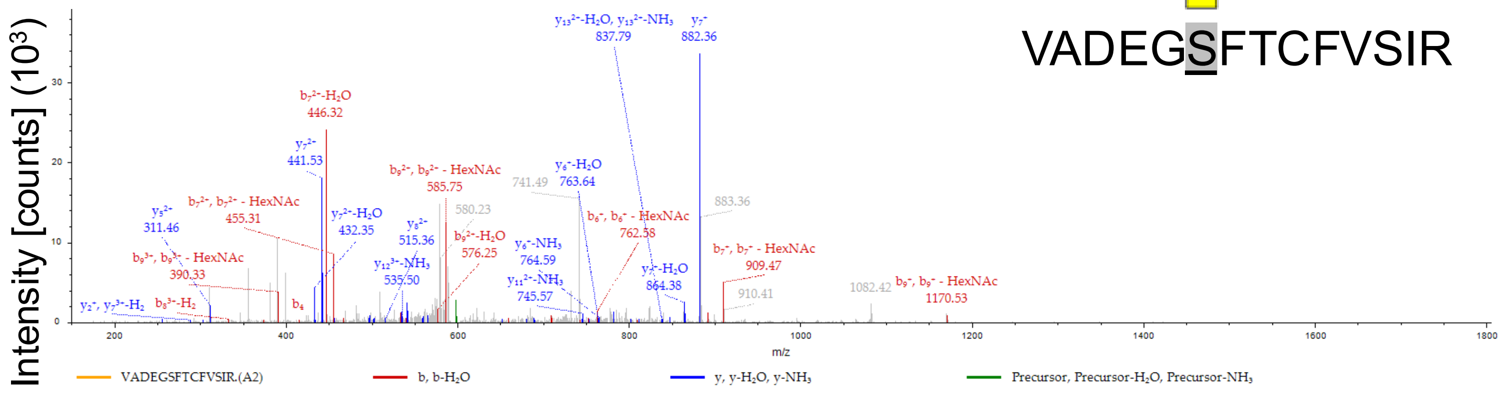


VADEGSFTCFVSIR

| #1 | $b^{+}$ | $b^{2+}$ | $b^{3+}$ | Seq. | $y^{+}$ | $y^{2+}$ | $y^{3+}$ | #2 |
|---|---|---|---|---|---|---|---|---|
| 1 | 100.07569 | 50.54148 | 34.03008 | V | | | | 14 |
| 2 | 171.11280 | 86.06004 | 57.70912 | A | 1691.75813 | 846.38270 | 564.59089 | 13 |
| 3 | 286.13975 | 143.57351 | 96.05143 | D | 1620.72101 | 810.86415 | 540.91186 | 12 |
| 4 | 415.18234 | 208.09481 | 139.06563 | E | 1505.69407 | 753.35067 | 502.56954 | 11 |
| 5 | 472.20380 | 236.60554 | 158.07279 | G | 1376.65148 | 688.82938 | 459.55534 | 10 |
| 6 | 762.31521 | 381.66124 | 254.77659 | S-HexNAc | 1319.63001 | 660.31865 | 440.54819 | 9 |
| 7 | 909.38362 | 455.19545 | 303.79939 | F | 1029.51861 | 515.26294 | 343.84439 | 8 |
| 8 | 1010.43130 | 505.71929 | 337.48195 | T | 882.45020 | 441.72874 | 294.82158 | 7 |
| 9 | 1170.46195 | 585.73461 | 390.82550 | -Carbamidometh | 781.40252 | 391.20490 | 261.13902 | 6 |
| 10 | 1317.53036 | 659.26882 | 439.84830 | F | 621.37187 | 311.18957 | 207.79548 | 5 |
| 11 | 1416.59877 | 708.80303 | 472.87111 | V | 474.30346 | 237.65537 | 158.77267 | 4 |
| 12 | 1503.63080 | 752.31904 | 501.88179 | S | 375.23504 | 188.12116 | 125.74987 | 3 |
| 13 | 1616.71487 | 808.86107 | 539.57647 | I | 288.20302 | 144.60515 | 96.73919 | 2 |
| 14 | | | | R | 175.11895 | 88.06311 | 59.04450 | 1 |

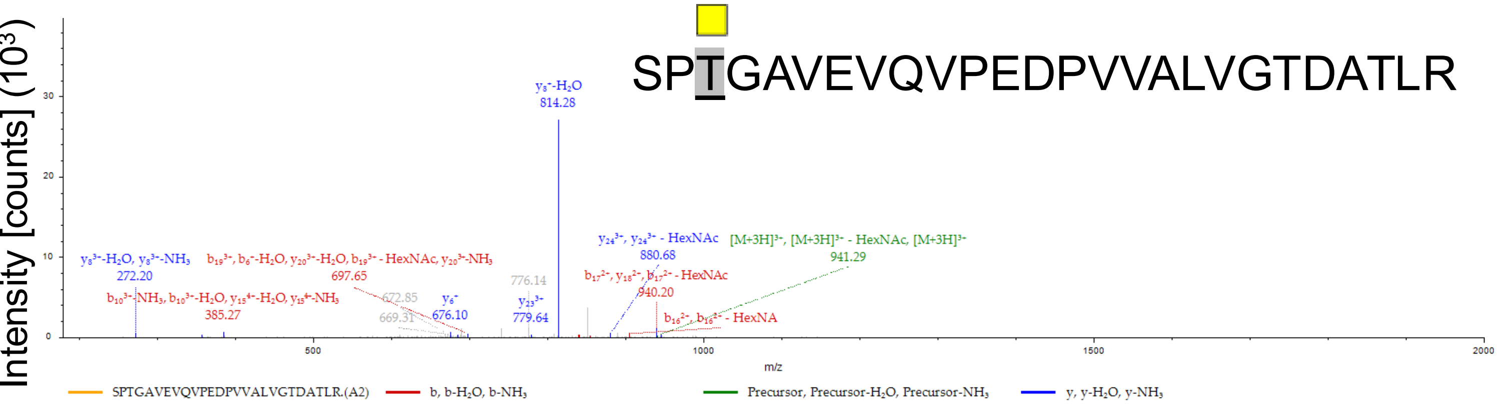


| #1 | b⁺ | b²⁺ | b³⁺ | b⁴⁺ | Seq. | y⁺ | y²⁺ | y³⁺ | y⁴⁺ | #2 |
|---|---|---|---|---|---|---|---|---|---|---|
| 1 | 88.03930 | 44.52329 | 30.01795 | 22.76528 | S | | | | | 26 |
| 2 | 185.09207 | 93.04967 | 62.36887 | 47.02847 | P | 2736.43018 | 1368.71873 | 912.81491 | 684.86300 | 25 |
| 3 | 489.21912 | 245.11320 | 163.74456 | 123.06024 | T-HexNAc | 2639.37742 | 1320.19235 | 880.46399 | 660.59981 | 24 |
| 4 | 546.24058 | 273.62393 | 182.75171 | 137.31560 | G | 2335.25037 | 1168.12882 | 779.08831 | 584.56805 | 23 |
| 5 | 617.27770 | 309.14249 | 206.43075 | 155.07488 | A | 2278.22890 | 1139.61809 | 760.08115 | 570.31268 | 22 |
| 6 | 716.34611 | 358.67669 | 239.45355 | 179.84199 | V | 2207.19179 | 1104.09953 | 736.40211 | 552.55340 | 21 |
| 7 | 845.38870 | 423.19799 | 282.46775 | 212.10263 | E | 2108.12337 | 1054.56533 | 703.37931 | 527.78630 | 20 |
| 8 | 944.45712 | 472.73220 | 315.49056 | 236.86974 | V | 1979.08078 | 990.04403 | 660.36511 | 495.52565 | 19 |
| 9 | 1072.51570 | 536.76149 | 358.17675 | 268.88438 | Q | 1880.01237 | 940.50982 | 627.34231 | 470.75855 | 18 |
| 10 | 1171.58411 | 586.29569 | 391.19955 | 293.65148 | V | 1751.95379 | 876.48053 | 584.65611 | 438.74391 | 17 |
| 11 | 1268.63687 | 634.82208 | 423.55048 | 317.91468 | P | 1652.88538 | 826.94633 | 551.63331 | 413.97680 | 16 |
| 12 | 1397.67947 | 699.34337 | 466.56467 | 350.17532 | E | 1555.83261 | 778.41994 | 519.28239 | 389.71361 | 15 |
| 13 | 1512.70641 | 756.85684 | 504.90699 | 378.93206 | D | 1426.79002 | 713.89865 | 476.26819 | 357.45296 | 14 |
| 14 | 1609.75917 | 805.38322 | 537.25791 | 403.19525 | P | 1311.76308 | 656.38518 | 437.92588 | 328.69623 | 13 |
| 15 | 1708.82759 | 854.91743 | 570.28071 | 427.96235 | V | 1214.71031 | 607.85879 | 405.57496 | 304.43304 | 12 |
| 16 | 1807.89600 | 904.45164 | 603.30352 | 452.72946 | V | 1115.64190 | 558.32459 | 372.55215 | 279.66593 | 11 |
| 17 | 1878.93311 | 939.97020 | 626.98256 | 470.48874 | A | 1016.57348 | 508.79038 | 339.52935 | 254.89883 | 10 |
| 18 | 1992.01718 | 996.51223 | 664.67724 | 498.75975 | L | 945.53637 | 473.27182 | 315.85031 | 237.13955 | 9 |
| 19 | 2091.08559 | 1046.04643 | 697.70005 | 523.52686 | V | 832.45231 | 416.72979 | 278.15562 | 208.86853 | 8 |
| 20 | 2148.10706 | 1074.55717 | 716.70720 | 537.78222 | G | 733.38389 | 367.19559 | 245.13282 | 184.10143 | 7 |
| 21 | 2249.15473 | 1125.08101 | 750.38976 | 563.04414 | T | 676.36243 | 338.68485 | 226.12566 | 169.84606 | 6 |
| 22 | 2364.18168 | 1182.59448 | 788.73208 | 591.80088 | D | 575.31475 | 288.16101 | 192.44310 | 144.58415 | 5 |
| 23 | 2435.21879 | 1218.11303 | 812.41111 | 609.56016 | A | 460.28781 | 230.64754 | 154.10079 | 115.82741 | 4 |
| 24 | 2536.26647 | 1268.63687 | 846.09367 | 634.82207 | T | 389.25069 | 195.12899 | 130.42175 | 98.06813 | 3 |
| 25 | 2649.35053 | 1325.17891 | 883.78836 | 663.09309 | L | 288.20302 | 144.60515 | 96.73919 | 72.80621 | 2 |
| 26 | | | | | R | 175.11895 | 88.06311 | 59.04450 | 44.53520 | 1 |

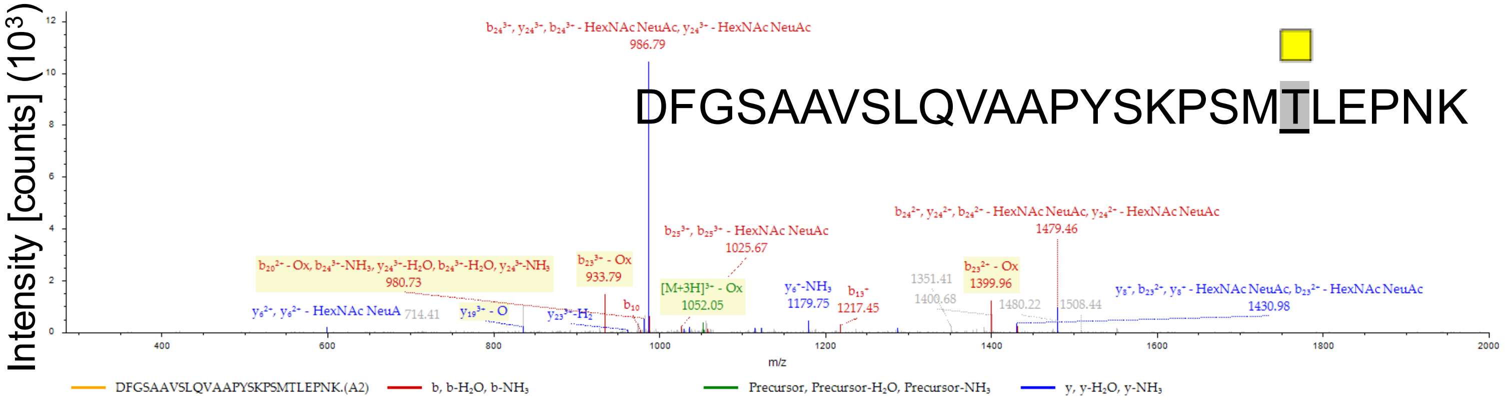


| #1 | b⁺ | b²⁺ | b³⁺ | b⁴⁺ | Seq. | y⁺ | y²⁺ | y³⁺ | y⁴⁺ | #2 |
|---|---|---|---|---|---|---|---|---|---|---|
| 1 | 116.03422 | 58.52075 | 39.34959 | 29.76401 | D | | | | | 26 |
| 2 | 263.10263 | 132.05496 | 88.37240 | 66.53112 | F | 3015.48671 | 1508.24699 | 1005.83375 | 754.62714 | 25 |
| 3 | 320.12410 | 160.56569 | 107.37955 | 80.78648 | G | 2868.41830 | 1434.71279 | 956.81095 | 717.86003 | 24 |
| 4 | 407.15613 | 204.08170 | 136.39023 | 102.54449 | S | 2811.39683 | 1406.20205 | 937.80380 | 703.60467 | 23 |
| 5 | 478.19324 | 239.60026 | 160.06926 | 120.30377 | A | 2724.36480 | 1362.68604 | 908.79312 | 681.84666 | 22 |
| 6 | 549.23035 | 275.11881 | 183.74830 | 138.06305 | A | 2653.32769 | 1327.16748 | 885.11408 | 664.08738 | 21 |
| 7 | 648.29877 | 324.65302 | 216.77111 | 162.83015 | V | 2582.29058 | 1291.64893 | 861.43504 | 646.32810 | 20 |
| 8 | 735.33080 | 368.16904 | 245.78178 | 184.58816 | S | 2483.22216 | 1242.11472 | 828.41224 | 621.56100 | 19 |
| 9 | 848.41486 | 424.71107 | 283.47647 | 212.85917 | L | 2396.19013 | 1198.59871 | 799.40156 | 599.80299 | 18 |
| 10 | 976.47344 | 488.74036 | 326.16266 | 244.87382 | Q | 2283.10607 | 1142.05667 | 761.70687 | 571.53198 | 17 |
| 11 | 1075.54185 | 538.27456 | 359.18547 | 269.64092 | V | 2155.04749 | 1078.02739 | 719.02068 | 539.51733 | 16 |
| 12 | 1146.57896 | 573.79312 | 382.86451 | 287.40020 | A | 2055.97908 | 1028.49318 | 685.99788 | 514.75023 | 15 |
| 13 | 1217.61608 | 609.31168 | 406.54354 | 305.15948 | A | 1984.94197 | 992.97462 | 662.31884 | 496.99095 | 14 |
| 14 | 1314.66884 | 657.83806 | 438.89447 | 329.42267 | P | 1913.90485 | 957.45606 | 638.63980 | 479.23167 | 13 |
| 15 | 1477.73217 | 739.36972 | 493.24891 | 370.18850 | Y | 1816.85209 | 908.92968 | 606.28888 | 454.96848 | 12 |
| 16 | 1564.76420 | 782.88574 | 522.25958 | 391.94651 | S | 1653.78876 | 827.39802 | 551.93444 | 414.20265 | 11 |
| 17 | 1692.85916 | 846.93322 | 564.95790 | 423.97025 | K | 1566.75673 | 783.88200 | 522.92376 | 392.44464 | 10 |
| 18 | 1789.91193 | 895.45960 | 597.30883 | 448.23344 | P | 1438.66177 | 719.83452 | 480.22544 | 360.42090 | 9 |
| 19 | 2080.02333 | 1040.51530 | 694.01263 | 520.76129 | S-HexNAc | 1341.60900 | 671.30814 | 447.87452 | 336.15771 | 8 |
| 20 | 2227.05873 | 1114.03300 | 743.02443 | 557.52014 | M-Oxidation | 1051.49760 | 526.25244 | 351.17072 | 263.62986 | 7 |
| 21 | 2531.18578 | 1266.09653 | 844.40011 | 633.55190 | T-HexNAc | 904.46220 | 452.73474 | 302.15892 | 226.87101 | 6 |
| 22 | 2644.26984 | 1322.63856 | 882.09480 | 661.82292 | L | 600.33515 | 300.67121 | 200.78324 | 150.83925 | 5 |
| 23 | 2773.31243 | 1387.15986 | 925.10900 | 694.08357 | E | 487.25109 | 244.12918 | 163.08855 | 122.56823 | 4 |
| 24 | 2870.36520 | 1435.68624 | 957.45992 | 718.34676 | P | 358.20850 | 179.60789 | 120.07435 | 90.30758 | 3 |
| 25 | 2984.40813 | 1492.70770 | 995.47423 | 746.85749 | N | 261.15573 | 131.08150 | 87.72343 | 66.04439 | 2 |
| 26 | | | | | K | 147.11280 | 74.06004 | 49.70912 | 37.53366 | 1 |

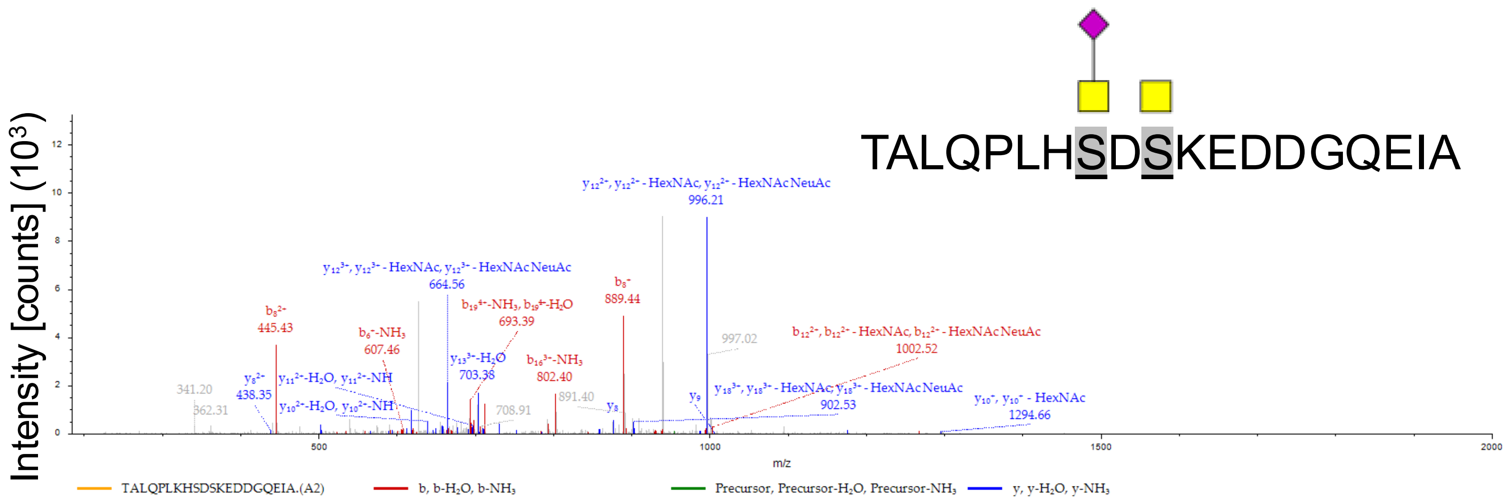


| #1 | $b^{+}$ | $b^{2+}$ | $b^{3+}$ | $b^{4+}$ | Seq. | $y^{+}$ | $y^{2+}$ | $y^{3+}$ | $y^{4+}$ | #2 |
|---|---|---|---|---|---|---|---|---|---|---|
| 1 | 102.05496 | 51.53112 | 34.68984 | 26.26920 | T | | | | | 20 |
| 2 | 173.09207 | 87.04967 | 58.36887 | 44.02847 | A | 2778.26871 | 1389.63800 | 926.76109 | 695.32264 | 19 |
| 3 | 286.17613 | 143.59170 | 96.06356 | 72.29949 | L | 2707.23160 | 1354.11944 | 903.08205 | 677.56336 | 18 |
| 4 | 414.23471 | 207.62099 | 138.74975 | 104.31414 | Q | 2594.14754 | 1297.57741 | 865.38736 | 649.29234 | 17 |
| 5 | 511.28747 | 256.14738 | 171.10068 | 128.57733 | P | 2466.08896 | 1233.54812 | 822.70117 | 617.27770 | 16 |
| 6 | 624.37154 | 312.68941 | 208.79536 | 156.84834 | L | 2369.03620 | 1185.02174 | 790.35025 | 593.01451 | 15 |
| 7 | 752.46650 | 376.73689 | 251.49368 | 188.87208 | K | 2255.95213 | 1128.47970 | 752.65556 | 564.74349 | 14 |
| 8 | 889.52541 | 445.26634 | 297.17999 | 223.13681 | H | 2127.85717 | 1064.43222 | 709.95724 | 532.71975 | 13 |
| 9 | 1470.73223 | 735.86975 | 490.91559 | 368.43851 | HexNAc(1)NeuAc | 1990.79826 | 995.90277 | 664.27094 | 498.45502 | 12 |
| 10 | 1585.75917 | 793.38322 | 529.25791 | 397.19525 | D | 1409.59144 | 705.29936 | 470.53533 | 353.15332 | 11 |
| 11 | 1875.87057 | 938.43893 | 625.96171 | 469.72310 | S-HexNAc | 1294.56450 | 647.78589 | 432.19302 | 324.39658 | 10 |
| 12 | 2003.96554 | 1002.48641 | 668.66003 | 501.74684 | K | 1004.45309 | 502.73019 | 335.48922 | 251.86873 | 9 |
| 13 | 2133.00813 | 1067.00770 | 711.67423 | 534.00749 | E | 876.35813 | 438.68270 | 292.79090 | 219.84499 | 8 |
| 14 | 2248.03507 | 1124.52117 | 750.01654 | 562.76423 | D | 747.31554 | 374.16141 | 249.77670 | 187.58434 | 7 |
| 15 | 2363.06202 | 1182.03465 | 788.35886 | 591.52096 | D | 632.28860 | 316.64794 | 211.43438 | 158.82761 | 6 |
| 16 | 2420.08348 | 1210.54538 | 807.36601 | 605.77633 | G | 517.26165 | 259.13446 | 173.09207 | 130.07087 | 5 |
| 17 | 2548.14206 | 1274.57467 | 850.05220 | 637.79097 | Q | 460.24019 | 230.62373 | 154.08491 | 115.81550 | 4 |
| 18 | 2677.18465 | 1339.09596 | 893.06640 | 670.05162 | E | 332.18161 | 166.59444 | 111.39872 | 83.80086 | 3 |
| 19 | 2790.26871 | 1395.63800 | 930.76109 | 698.32264 | I | 203.13902 | 102.07315 | 68.38452 | 51.54021 | 2 |
| 20 | | | | | A | 90.05496 | 45.53112 | 30.68984 | 23.26920 | 1 |

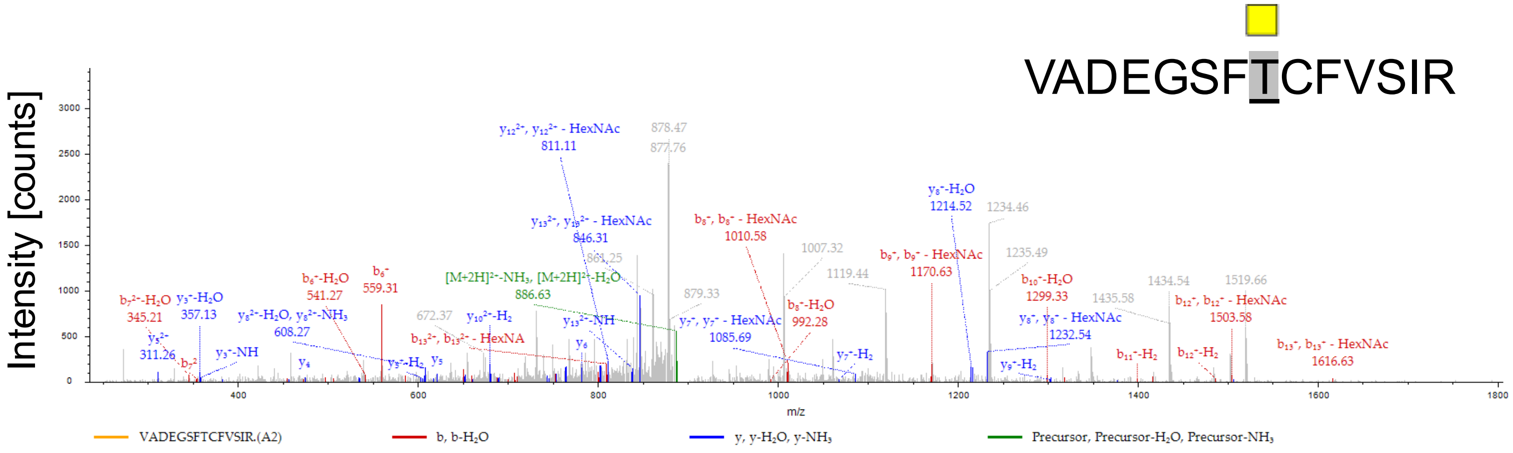


| #1 | b⁺ | b²⁺ | Seq. | y⁺ | y²⁺ | #2 |
|---|---|---|---|---|---|---|
| 1 | 100.07569 | 50.54148 | V | | | 14 |
| 2 | 171.11280 | 86.06004 | A | 1691.75813 | 846.38270 | 13 |
| 3 | 286.13975 | 143.57351 | D | 1620.72101 | 810.86415 | 12 |
| 4 | 415.18234 | 208.09481 | E | 1505.69407 | 753.35067 | 11 |
| 5 | 472.20380 | 236.60554 | G | 1376.65148 | 688.82938 | 10 |
| 6 | 559.23583 | 280.12155 | S | 1319.63001 | 660.31865 | 9 |
| 7 | 706.30425 | 353.65576 | F | 1232.59799 | 616.80263 | 8 |
| 8 | 1010.43130 | 505.71929 | T-HexNAc | 1085.52957 | 543.26842 | 7 |
| 9 | 1170.46195 | 585.73461 | -Carbamidometh | 781.40252 | 391.20490 | 6 |
| 10 | 1317.53036 | 659.26882 | F | 621.37187 | 311.18957 | 5 |
| 11 | 1416.59877 | 708.80303 | V | 474.30346 | 237.65537 | 4 |
| 12 | 1503.63080 | 752.31904 | S | 375.23504 | 188.12116 | 3 |
| 13 | 1616.71487 | 808.86107 | I | 288.20302 | 144.60515 | 2 |
| 14 | | | R | 175.11895 | 88.06311 | 1 |

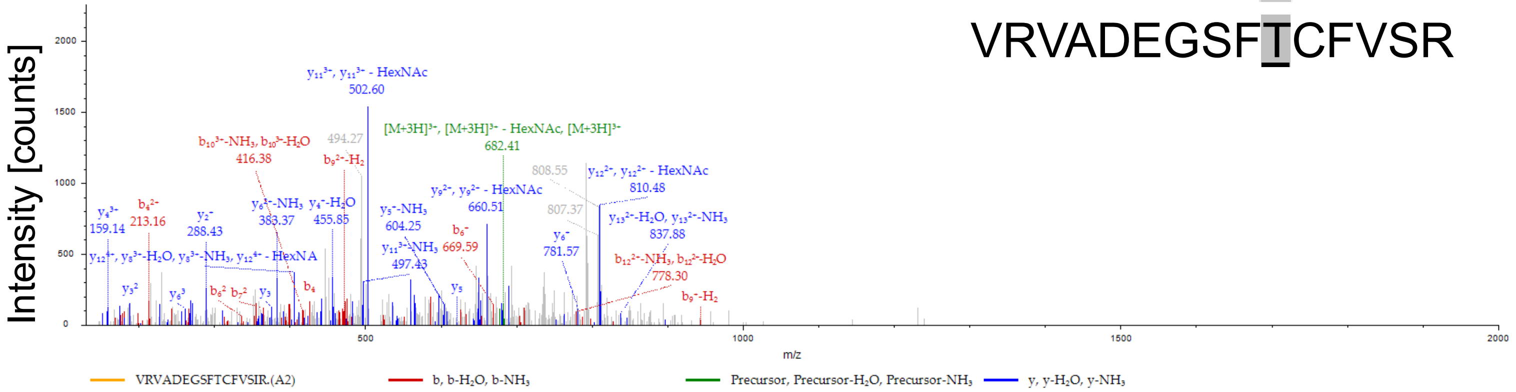


VRVADEGSFTCFVSR

| #1 | b⁺ | b²⁺ | b³⁺ | b⁴⁺ | Seq. | y⁺ | y²⁺ | y³⁺ | y⁴⁺ | #2 |
|---|---|---|---|---|---|---|---|---|---|---|
| 1 | 100.07569 | 50.54148 | 34.03008 | 25.77438 | V | | | | | 16 |
| 2 | 256.17680 | 128.59204 | 86.06378 | 64.79966 | R | 1946.92765 | 973.96746 | 649.64740 | 487.48737 | 15 |
| 3 | 355.24522 | 178.12625 | 119.08659 | 89.56676 | V | 1790.82654 | 895.91691 | 597.61370 | 448.46209 | 14 |
| 4 | 426.28233 | 213.64480 | 142.76563 | 107.32604 | A | 1691.75813 | 846.38270 | 564.59089 | 423.69499 | 13 |
| 5 | 541.30927 | 271.15827 | 181.10794 | 136.08278 | D | 1620.72101 | 810.86415 | 540.91186 | 405.93571 | 12 |
| 6 | 670.35187 | 335.67957 | 224.12214 | 168.34342 | E | 1505.69407 | 753.35067 | 502.56954 | 377.17898 | 11 |
| 7 | 727.37333 | 364.19030 | 243.12929 | 182.59879 | G | 1376.65148 | 688.82938 | 459.55534 | 344.91833 | 10 |
| 8 | 814.40536 | 407.70632 | 272.13997 | 204.35680 | S | 1319.63001 | 660.31865 | 440.54819 | 330.66296 | 9 |
| 9 | 961.47377 | 481.24052 | 321.16277 | 241.12390 | F | 1232.59799 | 616.80263 | 411.53751 | 308.90495 | 8 |
| 10 | 1265.60082 | 633.30405 | 422.53846 | 317.15566 | T-HexNAc | 1085.52957 | 543.26842 | 362.51471 | 272.13785 | 7 |
| 11 | 1425.63147 | 713.31937 | 475.88201 | 357.16333 | -Carbamidometh | 781.40252 | 391.20490 | 261.13902 | 196.10609 | 6 |
| 12 | 1572.69988 | 786.85358 | 524.90481 | 393.93043 | F | 621.37187 | 311.18957 | 207.79548 | 156.09843 | 5 |
| 13 | 1671.76830 | 836.38779 | 557.92762 | 418.69753 | V | 474.30346 | 237.65537 | 158.77267 | 119.33132 | 4 |
| 14 | 1758.80033 | 879.90380 | 586.93829 | 440.45554 | S | 375.23504 | 188.12116 | 125.74987 | 94.56422 | 3 |
| 15 | 1871.88439 | 936.44583 | 624.63298 | 468.72656 | I | 288.20302 | 144.60515 | 96.73919 | 72.80621 | 2 |
| 16 | | | | | R | 175.11895 | 88.06311 | 59.04450 | 44.53520 | 1 |

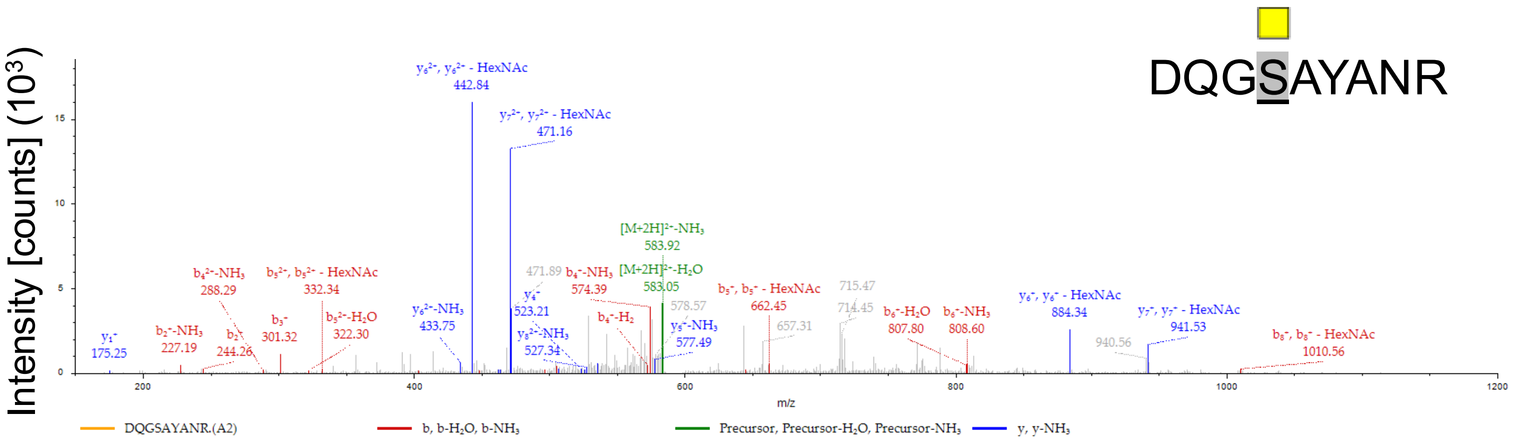


| #1 | b⁺ | b²⁺ | Seq. | y⁺ | y²⁺ | #2 |
|---|---|---|---|---|---|---|
| 1 | 116.03422 | 58.52075 | D | | | 9 |
| 2 | 244.09280 | 122.55004 | Q | 1069.49088 | 535.24908 | 8 |
| 3 | 301.11426 | 151.06077 | G | 941.43230 | 471.21979 | 7 |
| 4 | 591.22566 | 296.11647 | S-HexNAc | 884.41084 | 442.70906 | 6 |
| 5 | 662.26278 | 331.63503 | A | 594.29944 | 297.65336 | 5 |
| 6 | 825.32610 | 413.16669 | Y | 523.26232 | 262.13480 | 4 |
| 7 | 896.36322 | 448.68525 | A | 360.19899 | 180.60314 | 3 |
| 8 | 1010.40615 | 505.70671 | N | 289.16188 | 145.08458 | 2 |
| 9 | | | R | 175.11895 | 88.06311 | 1 |

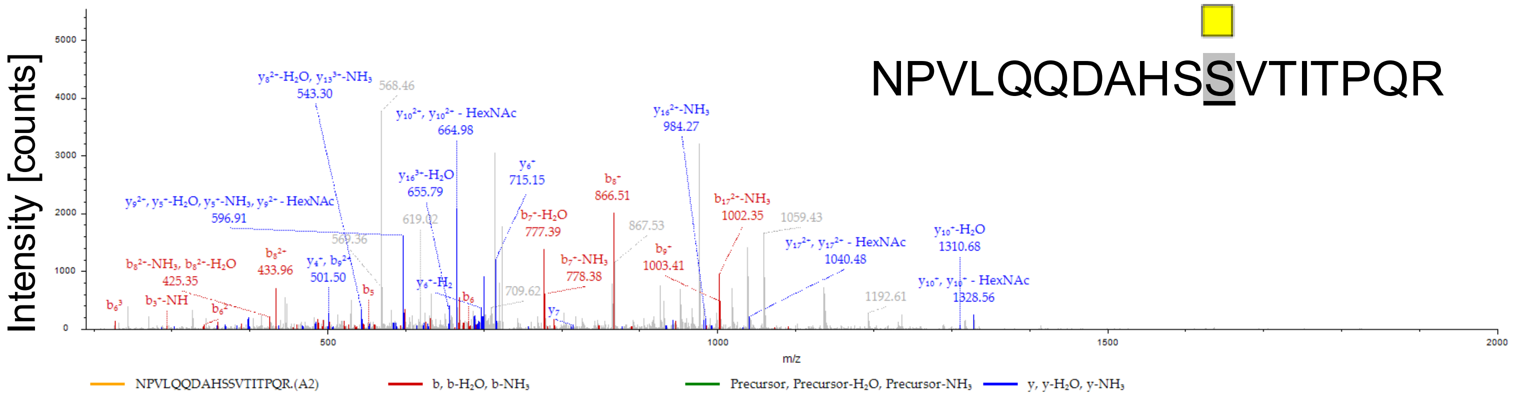


NPVLQQDAHSSVTITPQR

| #1 | $b^{+}$ | $b^{2+}$ | $b^{3+}$ | Seq. | $y^{+}$ | $y^{2+}$ | $y^{3+}$ | #2 |
|---|---|---|---|---|---|---|---|---|
| 1 | 115.05020 | 58.02874 | 39.02159 | N | | | | 18 |
| 2 | 212.10297 | 106.55512 | 71.37251 | P | 2080.06692 | 1040.53710 | 694.02716 | 17 |
| 3 | 311.17138 | 156.08933 | 104.39531 | V | 1983.01416 | 992.01072 | 661.67624 | 16 |
| 4 | 424.25545 | 212.63136 | 142.09000 | L | 1883.94575 | 942.47651 | 628.65343 | 15 |
| 5 | 552.31402 | 276.66065 | 184.77619 | Q | 1770.86168 | 885.93448 | 590.95874 | 14 |
| 6 | 680.37260 | 340.68994 | 227.46238 | Q | 1642.80310 | 821.90519 | 548.27255 | 13 |
| 7 | 795.39954 | 398.20341 | 265.80470 | D | 1514.74453 | 757.87590 | 505.58636 | 12 |
| 8 | 866.43666 | 433.72197 | 289.48374 | A | 1399.71758 | 700.36243 | 467.24405 | 11 |
| 9 | 1003.49557 | 502.25142 | 335.17004 | H | 1328.68047 | 664.84387 | 443.56501 | 10 |
| 10 | 1090.52760 | 545.76744 | 364.18072 | S | 1191.62156 | 596.31442 | 397.87870 | 9 |
| 11 | 1380.63900 | 690.82314 | 460.88452 | S-HexNAc | 1104.58953 | 552.79840 | 368.86803 | 8 |
| 12 | 1479.70741 | 740.35734 | 493.90732 | V | 814.47813 | 407.74270 | 272.16423 | 7 |
| 13 | 1580.75509 | 790.88118 | 527.58988 | T | 715.40971 | 358.20850 | 239.14142 | 6 |
| 14 | 1693.83915 | 847.42322 | 565.28457 | I | 614.36204 | 307.68466 | 205.45886 | 5 |
| 15 | 1794.88683 | 897.94705 | 598.96713 | T | 501.27797 | 251.14262 | 167.76418 | 4 |
| 16 | 1891.93960 | 946.47344 | 631.31805 | P | 400.23029 | 200.61879 | 134.08162 | 3 |
| 17 | 2019.99817 | 1010.50273 | 674.00424 | Q | 303.17753 | 152.09240 | 101.73069 | 2 |
| 18 | | | | R | 175.11895 | 88.06311 | 59.04450 | 1 |

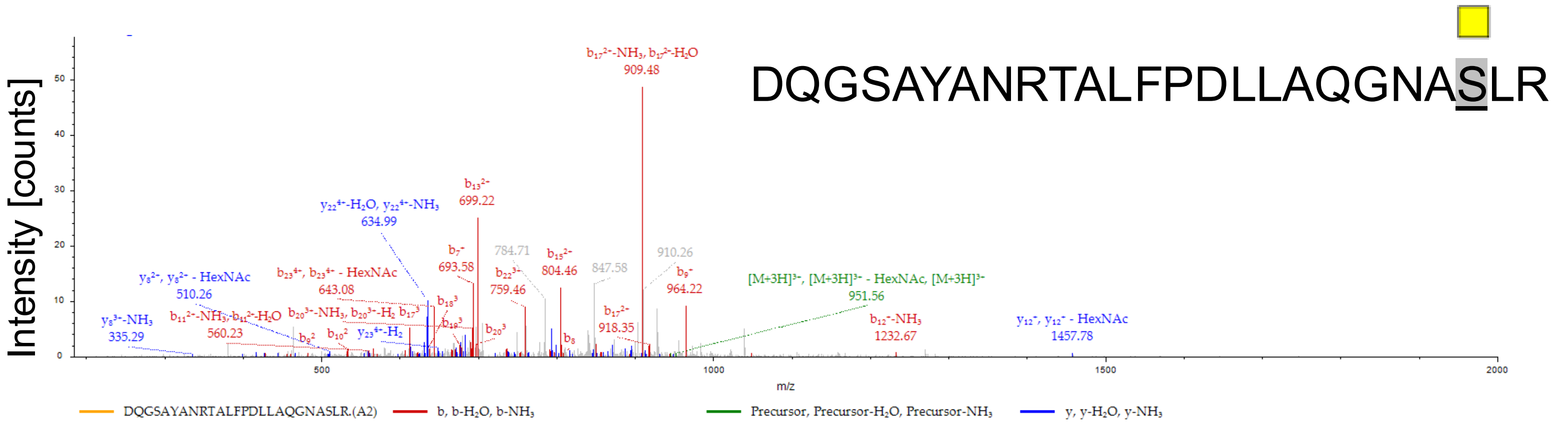


| #1 | $b^{+}$ | $b^{2+}$ | $b^{3+}$ | $b^{4+}$ | Seq. | $y^{+}$ | $y^{2+}$ | $y^{3+}$ | $y^{4+}$ | #2 |
|---|---|---|---|---|---|---|---|---|---|---|
| 1 | 116.03422 | 58.52075 | 39.34959 | 29.76401 | D | | | | | 25 |
| 2 | 244.09280 | 122.55004 | 82.03578 | 61.77866 | Q | 2738.37440 | 1369.69084 | 913.46298 | 685.34906 | 24 |
| 3 | 301.11426 | 151.06077 | 101.04294 | 76.03402 | G | 2610.31582 | 1305.66155 | 870.77679 | 653.33441 | 23 |
| 4 | 388.14629 | 194.57678 | 130.05361 | 97.79203 | S | 2553.29436 | 1277.15082 | 851.76964 | 639.07905 | 22 |
| 5 | 459.18340 | 230.09534 | 153.73265 | 115.55131 | A | 2466.26233 | 1233.63480 | 822.75896 | 617.32104 | 21 |
| 6 | 622.24673 | 311.62700 | 208.08709 | 156.31714 | Y | 2395.22521 | 1198.11625 | 799.07992 | 599.56176 | 20 |
| 7 | 693.28385 | 347.14556 | 231.76613 | 174.07642 | A | 2232.16189 | 1116.58458 | 744.72548 | 558.79593 | 19 |
| 8 | 808.31079 | 404.65903 | 270.10845 | 202.83315 | N-Deamidated | 2161.12477 | 1081.06602 | 721.04644 | 541.03665 | 18 |
| 9 | 964.41190 | 482.70959 | 322.14215 | 241.85843 | R | 2046.09783 | 1023.55255 | 682.70413 | 512.27991 | 17 |
| 10 | 1065.45958 | 533.23343 | 355.82471 | 267.12035 | T | 1889.99672 | 945.50200 | 630.67042 | 473.25464 | 16 |
| 11 | 1136.49669 | 568.75198 | 379.50375 | 284.87963 | A | 1788.94904 | 894.97816 | 596.98786 | 447.99272 | 15 |
| 12 | 1249.58076 | 625.29402 | 417.19844 | 313.15065 | L | 1717.91193 | 859.45960 | 573.30883 | 430.23344 | 14 |
| 13 | 1396.64917 | 698.82822 | 466.22124 | 349.91775 | F | 1604.82786 | 802.91757 | 535.61414 | 401.96242 | 13 |
| 14 | 1493.70193 | 747.35460 | 498.57216 | 374.18094 | P | 1457.75945 | 729.38336 | 486.59133 | 365.19532 | 12 |
| 15 | 1608.72888 | 804.86808 | 536.91448 | 402.93768 | D | 1360.70668 | 680.85698 | 454.24041 | 340.93213 | 11 |
| 16 | 1721.81294 | 861.41011 | 574.60916 | 431.20869 | L | 1245.67974 | 623.34351 | 415.89810 | 312.17539 | 10 |
| 17 | 1834.89700 | 917.95214 | 612.30385 | 459.47971 | L | 1132.59568 | 566.80148 | 378.20341 | 283.90438 | 9 |
| 18 | 1905.93412 | 953.47070 | 635.98289 | 477.23899 | A | 1019.51161 | 510.25945 | 340.50872 | 255.63336 | 8 |
| 19 | 2033.99270 | 1017.49999 | 678.66908 | 509.25363 | Q | 948.47450 | 474.74089 | 316.82968 | 237.87408 | 7 |
| 20 | 2091.01416 | 1046.01072 | 697.67624 | 523.50900 | G | 820.41592 | 410.71160 | 274.14349 | 205.85944 | 6 |
| 21 | 2205.05709 | 1103.03218 | 735.69055 | 552.01973 | N | 763.39446 | 382.20087 | 255.13634 | 191.60407 | 5 |
| 22 | 2276.09420 | 1138.55074 | 759.36958 | 569.77901 | A | 649.35153 | 325.17940 | 217.12203 | 163.09334 | 4 |
| 23 | 2566.20560 | 1283.60644 | 856.07338 | 642.30686 | S-HexNAc | 578.31442 | 289.66085 | 193.44299 | 145.33406 | 3 |
| 24 | 2679.28967 | 1340.14847 | 893.76807 | 670.57787 | L | 288.20302 | 144.60515 | 96.73919 | 72.80621 | 2 |
| 25 | | | | | R | 175.11895 | 88.06311 | 59.04450 | 44.53520 | 1 |

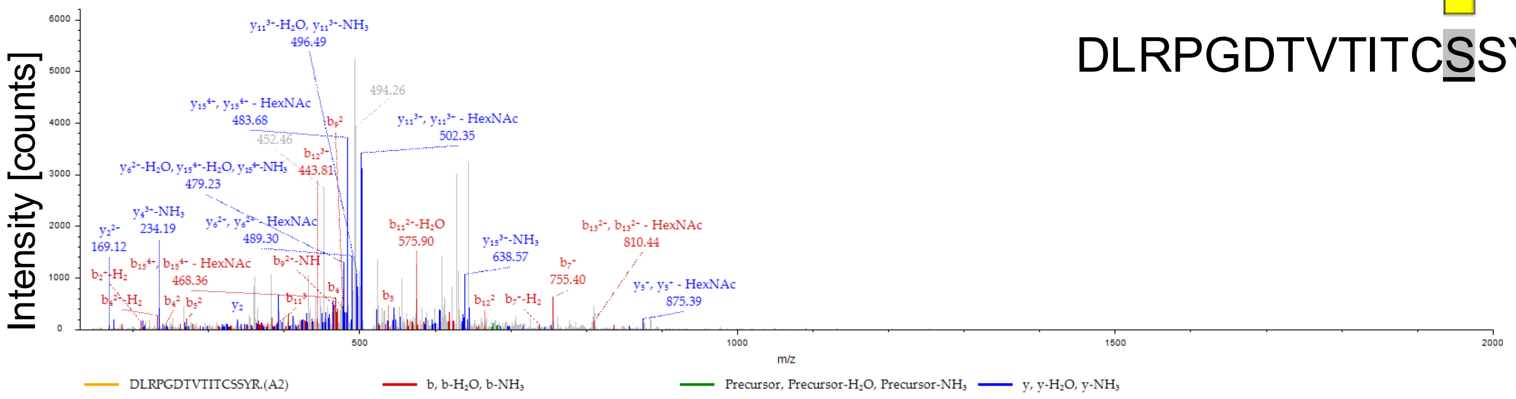


DLRPGDTVTITCSSYR

| #1 | $b^+$ | $b^{2+}$ | $b^{3+}$ | Seq. | $y^+$ | $y^{2+}$ | $y^{3+}$ | #2 |
|---|---|---|---|---|---|---|---|---|
| 1 | 116.03422 | 58.52075 | 39.34959 | D | | | | 16 |
| 2 | 229.11828 | 115.06278 | 77.04428 | L | 1928.93822 | 964.97275 | 643.65092 | 15 |
| 3 | 385.21939 | 193.11334 | 129.07798 | R | 1815.85415 | 908.43071 | 605.95624 | 14 |
| 4 | 482.27216 | 241.63972 | 161.42890 | P | 1659.75304 | 830.38016 | 553.92253 | 13 |
| 5 | 539.29362 | 270.15045 | 180.43606 | G | 1562.70028 | 781.85378 | 521.57161 | 12 |
| 6 | 654.32056 | 327.66392 | 218.77837 | D | 1505.67881 | 753.34305 | 502.56446 | 11 |
| 7 | 755.36824 | 378.18776 | 252.46093 | T | 1390.65187 | 695.82957 | 464.22214 | 10 |
| 8 | 854.43666 | 427.72197 | 285.48374 | V | 1289.60419 | 645.30574 | 430.53958 | 9 |
| 9 | 955.48434 | 478.24581 | 319.16630 | T | 1190.53578 | 595.77153 | 397.51678 | 8 |
| 10 | 1068.56840 | 534.78784 | 356.86098 | I | 1089.48810 | 545.24769 | 363.83422 | 7 |
| 11 | 1169.61608 | 585.31168 | 390.54354 | T | 976.40404 | 488.70566 | 326.13953 | 6 |
| 12 | 1329.64673 | 665.32700 | 443.88709 | -Carbamidometh | 875.35636 | 438.18182 | 292.45697 | 5 |
| 13 | 1619.75813 | 810.38270 | 540.59089 | S-HexNAc | 715.32571 | 358.16649 | 239.11342 | 4 |
| 14 | 1706.79016 | 853.89872 | 569.60157 | S | 425.21431 | 213.11079 | 142.40962 | 3 |
| 15 | 1869.85348 | 935.43038 | 623.95601 | Y | 338.18228 | 169.59478 | 113.39894 | 2 |
| 16 | | | | R | 175.11895 | 88.06311 | 59.04450 | 1 |

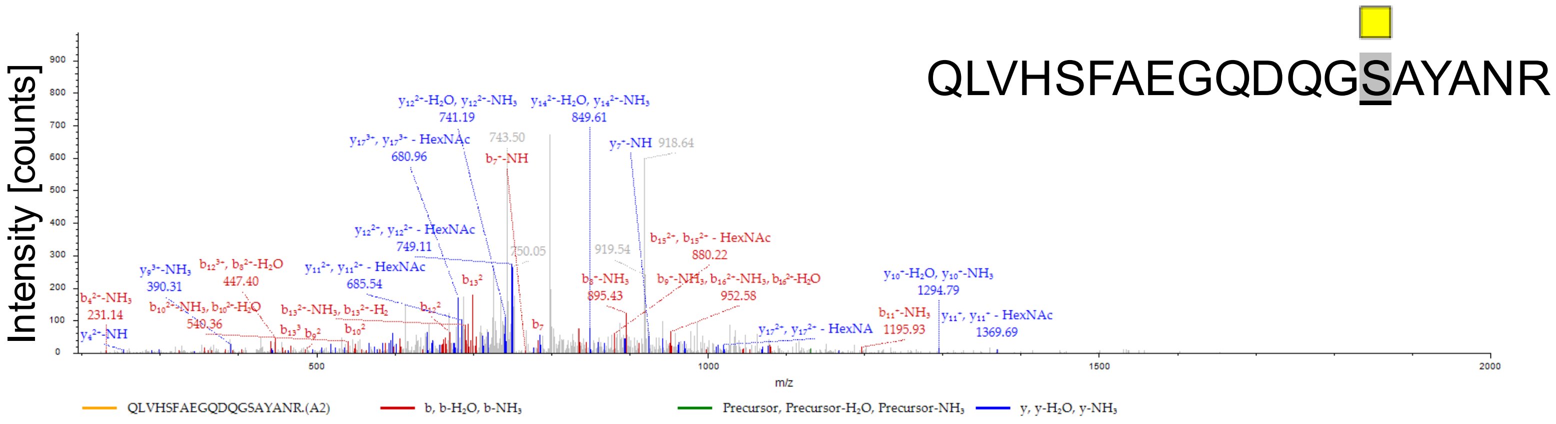


| #1 | b⁺ | b²⁺ | b³⁺ | Seq. | y⁺ | y²⁺ | y³⁺ | #2 |
|---|---|---|---|---|---|---|---|---|
| 1 | 129.06585 | 65.03657 | 43.69347 | Q | | | | 19 |
| 2 | 242.14992 | 121.57860 | 81.38816 | L | 2152.98940 | 1076.99834 | 718.33465 | 18 |
| 3 | 341.21833 | 171.11280 | 114.41096 | V | 2039.90534 | 1020.45631 | 680.63996 | 17 |
| 4 | 478.27724 | 239.64226 | 160.09727 | H | 1940.83692 | 970.92210 | 647.61716 | 16 |
| 5 | 565.30927 | 283.15827 | 189.10794 | S | 1803.77801 | 902.39264 | 601.93085 | 15 |
| 6 | 712.37769 | 356.69248 | 238.13075 | F | 1716.74598 | 858.87663 | 572.92018 | 14 |
| 7 | 783.41480 | 392.21104 | 261.80978 | A | 1569.67757 | 785.34242 | 523.89737 | 13 |
| 8 | 912.45739 | 456.73233 | 304.82398 | E | 1498.64046 | 749.82387 | 500.21834 | 12 |
| 9 | 969.47886 | 485.24307 | 323.83114 | G | 1369.59786 | 685.30257 | 457.20414 | 11 |
| 10 | 1097.53743 | 549.27236 | 366.51733 | Q | 1312.57640 | 656.79184 | 438.19698 | 10 |
| 11 | 1212.56438 | 606.78583 | 404.85964 | D | 1184.51782 | 592.76255 | 395.51079 | 9 |
| 12 | 1340.62295 | 670.81512 | 447.54584 | Q | 1069.49088 | 535.24908 | 357.16848 | 8 |
| 13 | 1397.64442 | 699.32585 | 466.55299 | G | 941.43230 | 471.21979 | 314.48228 | 7 |
| 14 | 1687.75582 | 844.38155 | 563.25679 | S-HexNAc | 884.41084 | 442.70906 | 295.47513 | 6 |
| 15 | 1758.79293 | 879.90010 | 586.93583 | A | 594.29944 | 297.65336 | 198.77133 | 5 |
| 16 | 1921.85626 | 961.43177 | 641.29027 | Y | 523.26232 | 262.13480 | 175.09229 | 4 |
| 17 | 1992.89338 | 996.95033 | 664.96931 | A | 360.19899 | 180.60314 | 120.73785 | 3 |
| 18 | 2106.93630 | 1053.97179 | 702.98362 | N | 289.16188 | 145.08458 | 97.05881 | 2 |
| 19 | | | | R | 175.11895 | 88.06311 | 59.04450 | 1 |

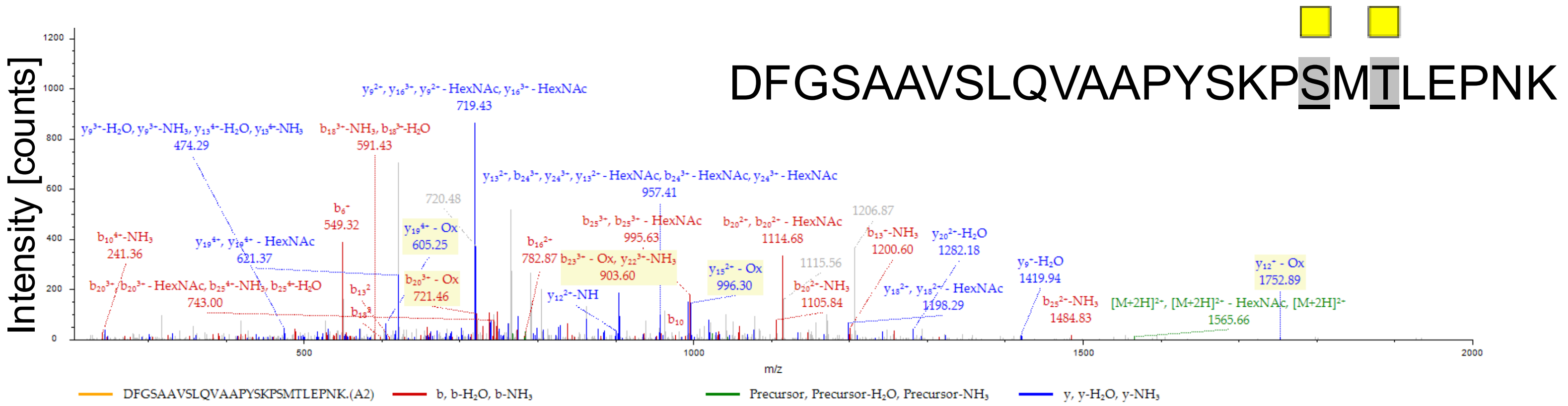


| #1 | b⁺ | b²⁺ | b³⁺ | b⁴⁺ | Seq. | y⁺ | y²⁺ | y³⁺ | y⁴⁺ | #2 |
|---|---|---|---|---|---|---|---|---|---|---|
| 1 | 116.03422 | 58.52075 | 39.34959 | 29.76401 | D | | | | | 26 |
| 2 | 263.10263 | 132.05496 | 88.37240 | 66.53112 | F | 3015.48671 | 1508.24699 | 1005.83375 | 754.62714 | 25 |
| 3 | 320.12410 | 160.56569 | 107.37955 | 80.78648 | G | 2868.41830 | 1434.71279 | 956.81095 | 717.86003 | 24 |
| 4 | 407.15613 | 204.08170 | 136.39023 | 102.54449 | S | 2811.39683 | 1406.20205 | 937.80380 | 703.60467 | 23 |
| 5 | 478.19324 | 239.60026 | 160.06926 | 120.30377 | A | 2724.36480 | 1362.68604 | 908.79312 | 681.84666 | 22 |
| 6 | 549.23035 | 275.11881 | 183.74830 | 138.06305 | A | 2653.32769 | 1327.16748 | 885.11408 | 664.08738 | 21 |
| 7 | 648.29877 | 324.65302 | 216.77111 | 162.83015 | V | 2582.29058 | 1291.64893 | 861.43504 | 646.32810 | 20 |
| 8 | 735.33080 | 368.16904 | 245.78178 | 184.58816 | S | 2483.22216 | 1242.11472 | 828.41224 | 621.56100 | 19 |
| 9 | 848.41486 | 424.71107 | 283.47647 | 212.85917 | L | 2396.19013 | 1198.59871 | 799.40156 | 599.80299 | 18 |
| 10 | 976.47344 | 488.74036 | 326.16266 | 244.87382 | Q | 2283.10607 | 1142.05667 | 761.70687 | 571.53198 | 17 |
| 11 | 1075.54185 | 538.27456 | 359.18547 | 269.64092 | V | 2155.04749 | 1078.02739 | 719.02068 | 539.51733 | 16 |
| 12 | 1146.57896 | 573.79312 | 382.86451 | 287.40020 | A | 2055.97908 | 1028.49318 | 685.99788 | 514.75023 | 15 |
| 13 | 1217.61608 | 609.31168 | 406.54354 | 305.15948 | A | 1984.94197 | 992.97462 | 662.31884 | 496.99095 | 14 |
| 14 | 1314.66884 | 657.83806 | 438.89447 | 329.42267 | P | 1913.90485 | 957.45606 | 638.63980 | 479.23167 | 13 |
| 15 | 1477.73217 | 739.36972 | 493.24891 | 370.18850 | Y | 1816.85209 | 908.92968 | 606.28888 | 454.96848 | 12 |
| 16 | 1564.76420 | 782.88574 | 522.25958 | 391.94651 | S | 1653.78876 | 827.39802 | 551.93444 | 414.20265 | 11 |
| 17 | 1692.85916 | 846.93322 | 564.95790 | 423.97025 | K | 1566.75673 | 783.88200 | 522.92376 | 392.44464 | 10 |
| 18 | 1789.91193 | 895.45960 | 597.30883 | 448.23344 | P | 1438.66177 | 719.83452 | 480.22544 | 360.42090 | 9 |
| 19 | 2080.02333 | 1040.51530 | 694.01263 | 520.76129 | S-HexNAc | 1341.60900 | 671.30814 | 447.87452 | 336.15771 | 8 |
| 20 | 2227.05873 | 1114.03300 | 743.02443 | 557.52014 | M-Oxidation | 1051.49760 | 526.25244 | 351.17072 | 263.62986 | 7 |
| 21 | 2531.18578 | 1266.09653 | 844.40011 | 633.55190 | T-HexNAc | 904.46220 | 452.73474 | 302.15892 | 226.87101 | 6 |
| 22 | 2644.26984 | 1322.63856 | 882.09480 | 661.82292 | L | 600.33515 | 300.67121 | 200.78324 | 150.83925 | 5 |
| 23 | 2773.31243 | 1387.15986 | 925.10900 | 694.08357 | E | 487.25109 | 244.12918 | 163.08855 | 122.56823 | 4 |
| 24 | 2870.36520 | 1435.68624 | 957.45992 | 718.34676 | P | 358.20850 | 179.60789 | 120.07435 | 90.30758 | 3 |
| 25 | 2984.40813 | 1492.70770 | 995.47423 | 746.85749 | N | 261.15573 | 131.08150 | 87.72343 | 66.04439 | 2 |
| 26 | | | | | K | 147.11280 | 74.06004 | 49.70912 | 37.53366 | 1 |

O

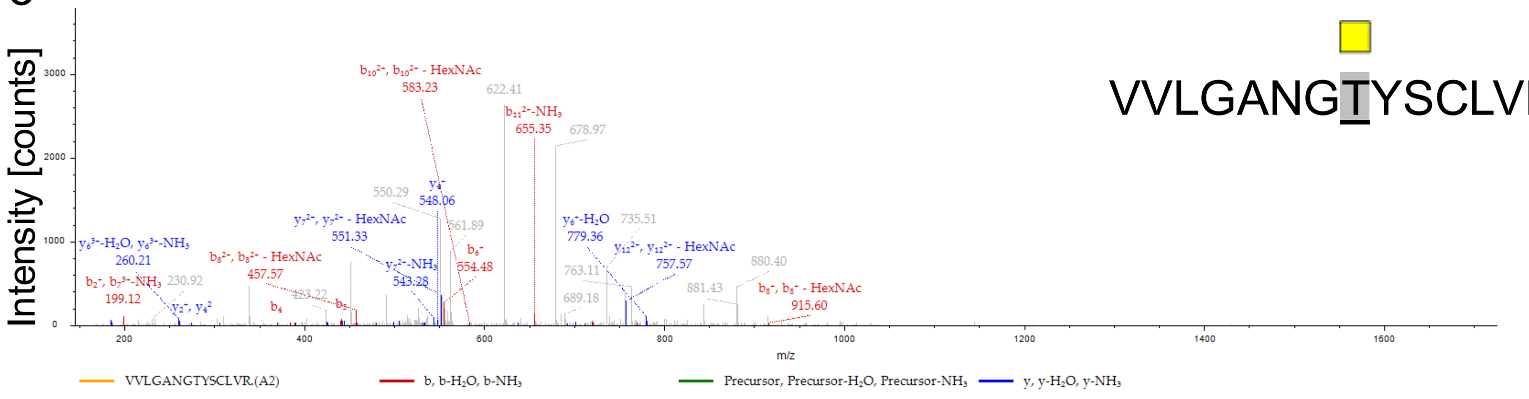


VVLGANGTYSCLVR

| #1 | $b^+$ | $b^{2+}$ | $b^{3+}$ | Seq. | $y^+$ | $y^{2+}$ | $y^{3+}$ | #2 |
|---|---|---|---|---|---|---|---|---|
| 1 | 100.07569 | 50.54148 | 34.03008 | V | | | | 14 |
| 2 | 199.14410 | 100.07569 | 67.05289 | V | 1612.79993 | 806.90360 | 538.27150 | 13 |
| 3 | 312.22817 | 156.61772 | 104.74757 | L | 1513.73152 | 757.36940 | 505.24869 | 12 |
| 4 | 369.24963 | 185.12845 | 123.75473 | G | 1400.64746 | 700.82737 | 467.55400 | 11 |
| 5 | 440.28675 | 220.64701 | 147.43377 | A | 1343.62599 | 672.31663 | 448.54685 | 10 |
| 6 | 554.32967 | 277.66847 | 185.44808 | N | 1272.58888 | 636.79808 | 424.86781 | 9 |
| 7 | 611.35114 | 306.17921 | 204.45523 | G | 1158.54595 | 579.77661 | 386.85350 | 8 |
| 8 | 915.47819 | 458.24273 | 305.83091 | T-HexNAc | 1101.52449 | 551.26588 | 367.84635 | 7 |
| 9 | 1078.54152 | 539.77440 | 360.18536 | Y | 797.39744 | 399.20236 | 266.47066 | 6 |
| 10 | 1165.57355 | 583.29041 | 389.19603 | S | 634.33411 | 317.67069 | 212.11622 | 5 |
| 11 | 1325.60419 | 663.30574 | 442.53958 | -Carbamidometh | 547.30208 | 274.15468 | 183.10554 | 4 |
| 12 | 1438.68826 | 719.84777 | 480.23427 | L | 387.27143 | 194.13935 | 129.76199 | 3 |
| 13 | 1537.75667 | 769.38197 | 513.25707 | V | 274.18737 | 137.59732 | 92.06731 | 2 |
| 14 | | | | R | 175.11895 | 88.06311 | 59.04450 | 1 |

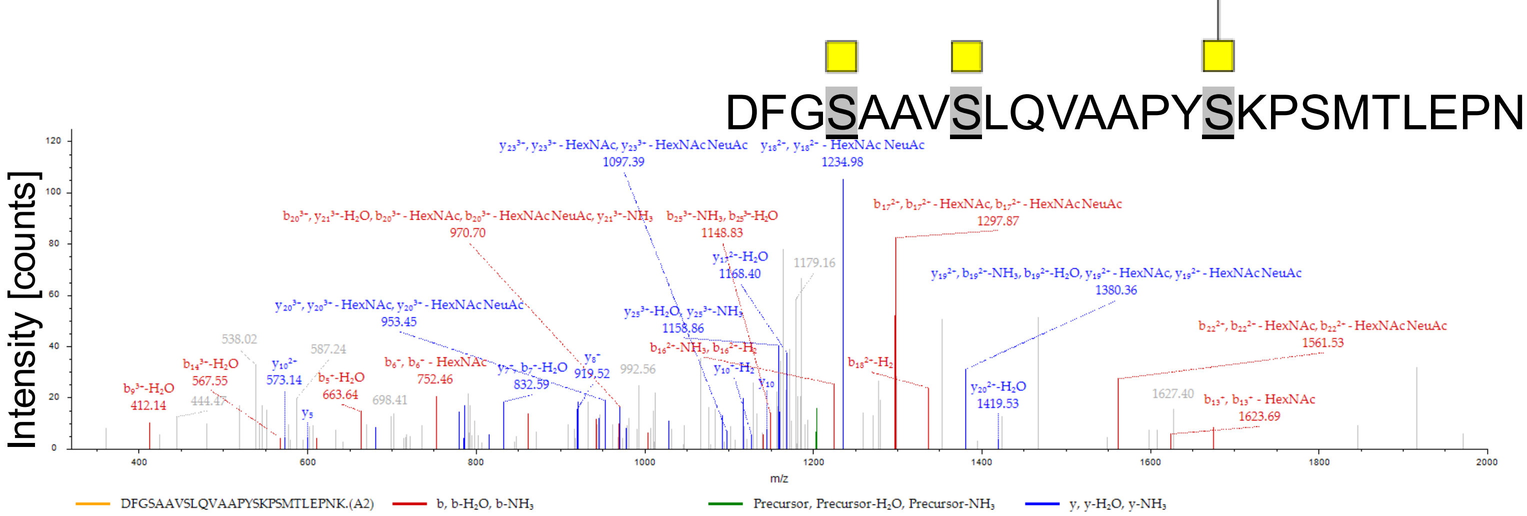


| #1 | $b^{+}$ | $b^{2+}$ | $b^{3+}$ | Seq. | $y^{+}$ | $y^{2+}$ | $y^{3+}$ | #2 |
|---|---|---|---|---|---|---|---|---|
| 1 | 116.03422 | 58.52075 | 39.34959 | D | | | | 26 |
| 2 | 263.10263 | 132.05496 | 88.37240 | F | 3493.66658 | 1747.33693 | 1165.22705 | 25 |
| 3 | 320.12410 | 160.56569 | 107.37955 | G | 3346.59817 | 1673.80272 | 1116.20424 | 24 |
| 4 | 610.23550 | 305.62139 | 204.08335 | S-HexNAc | 3289.57671 | 1645.29199 | 1097.19709 | 23 |
| 5 | 681.27261 | 341.13994 | 227.76239 | A | 2999.46531 | 1500.23629 | 1000.49329 | 22 |
| 6 | 752.30973 | 376.65850 | 251.44143 | A | 2928.42819 | 1464.71773 | 976.81425 | 21 |
| 7 | 851.37814 | 426.19271 | 284.46423 | V | 2857.39108 | 1429.19918 | 953.13521 | 20 |
| 8 | 1141.48954 | 571.24841 | 381.16803 | S-HexNAc | 2758.32266 | 1379.66497 | 920.11241 | 19 |
| 9 | 1254.57361 | 627.79044 | 418.86272 | L | 2468.21126 | 1234.60927 | 823.40861 | 18 |
| 10 | 1382.63218 | 691.81973 | 461.54891 | Q | 2355.12720 | 1178.06724 | 785.71392 | 17 |
| 11 | 1481.70060 | 741.35394 | 494.57172 | V | 2227.06862 | 1114.03795 | 743.02772 | 16 |
| 12 | 1552.73771 | 776.87249 | 518.25075 | A | 2128.00021 | 1064.50374 | 710.00492 | 15 |
| 13 | 1623.77482 | 812.39105 | 541.92979 | A | 2056.96309 | 1028.98519 | 686.32588 | 14 |
| 14 | 1720.82759 | 860.91743 | 574.28071 | P | 1985.92598 | 993.46663 | 662.64684 | 13 |
| 15 | 1883.89092 | 942.44910 | 628.63516 | Y | 1888.87322 | 944.94025 | 630.29592 | 12 |
| 16 | 2465.09773 | 1233.05251 | 822.37076 | lexNAc(1)NeuAc | 1725.80989 | 863.40858 | 575.94148 | 11 |
| 17 | 2593.19270 | 1297.09999 | 865.06908 | K | 1144.60307 | 572.80517 | 382.20587 | 10 |
| 18 | 2690.24546 | 1345.62637 | 897.42000 | P | 1016.50811 | 508.75769 | 339.50755 | 9 |
| 19 | 2777.27749 | 1389.14238 | 926.43068 | S | 919.45534 | 460.23131 | 307.15663 | 8 |
| 20 | 2908.31797 | 1454.66262 | 970.11084 | M | 832.42332 | 416.71530 | 278.14596 | 7 |
| 21 | 3009.36565 | 1505.18646 | 1003.79340 | T | 701.38283 | 351.19505 | 234.46579 | 6 |
| 22 | 3122.44972 | 1561.72850 | 1041.48809 | L | 600.33515 | 300.67121 | 200.78324 | 5 |
| 23 | 3251.49231 | 1626.24979 | 1084.50229 | E | 487.25109 | 244.12918 | 163.08855 | 4 |
| 24 | 3348.54507 | 1674.77617 | 1116.85321 | P | 358.20850 | 179.60789 | 120.07435 | 3 |
| 25 | 3462.58800 | 1731.79764 | 1154.86752 | N | 261.15573 | 131.08150 | 87.72343 | 2 |
| 26 | | | | K | 147.11280 | 74.06004 | 49.70912 | 1 |

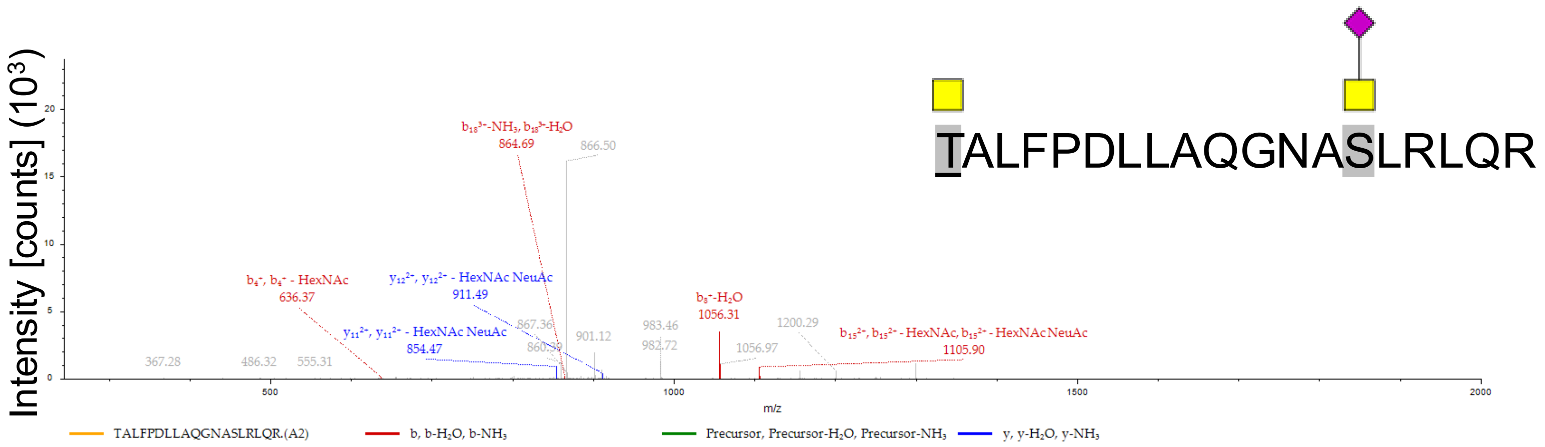


| #1 | $b^+$ | $b^{2+}$ | $b^{3+}$ | Seq. | $y^+$ | $y^{2+}$ | $y^{3+}$ | #2 |
|---|---|---|---|---|---|---|---|---|
| 1 | 305.13433 | 153.07080 | 102.38296 | T-HexNAc | | | | 19 |
| 2 | 376.17144 | 188.58936 | 126.06200 | A | 2478.27222 | 1239.63975 | 826.76226 | 18 |
| 3 | 489.25551 | 245.13139 | 163.75669 | L | 2407.23511 | 1204.12119 | 803.08322 | 17 |
| 4 | 636.32392 | 318.66560 | 212.77949 | F | 2294.15105 | 1147.57916 | 765.38853 | 16 |
| 5 | 733.37668 | 367.19198 | 245.13041 | P | 2147.08263 | 1074.04495 | 716.36573 | 15 |
| 6 | 848.40363 | 424.70545 | 283.47273 | D | 2050.02987 | 1025.51857 | 684.01481 | 14 |
| 7 | 961.48769 | 481.24748 | 321.16741 | L | 1935.00293 | 968.00510 | 645.67249 | 13 |
| 8 | 1074.57175 | 537.78952 | 358.86210 | L | 1821.91886 | 911.46307 | 607.97781 | 12 |
| 9 | 1145.60887 | 573.30807 | 382.54114 | A | 1708.83480 | 854.92104 | 570.28312 | 11 |
| 10 | 1273.66745 | 637.33736 | 425.22733 | Q | 1637.79768 | 819.40248 | 546.60408 | 10 |
| 11 | 1330.68891 | 665.84809 | 444.23449 | G | 1509.73911 | 755.37319 | 503.91789 | 9 |
| 12 | 1445.71585 | 723.36156 | 482.57680 | N-Deamidated | 1452.71764 | 726.86246 | 484.91073 | 8 |
| 13 | 1516.75297 | 758.88012 | 506.25584 | A | 1337.69070 | 669.34899 | 446.56842 | 7 |
| 14 | 2097.95978 | 1049.48353 | 699.99145 | HexNAc(1)NeuAc | 1266.65359 | 633.83043 | 422.88938 | 6 |
| 15 | 2211.04385 | 1106.02556 | 737.68613 | L | 685.44677 | 343.22702 | 229.15377 | 5 |
| 16 | 2367.14496 | 1184.07612 | 789.71984 | R | 572.36270 | 286.68499 | 191.45909 | 4 |
| 17 | 2480.22902 | 1240.61815 | 827.41453 | L | 416.26159 | 208.63444 | 139.42538 | 3 |
| 18 | 2608.28760 | 1304.64744 | 870.10072 | Q | 303.17753 | 152.09240 | 101.73069 | 2 |
| 19 | | | | R | 175.11895 | 88.06311 | 59.04450 | 1 |

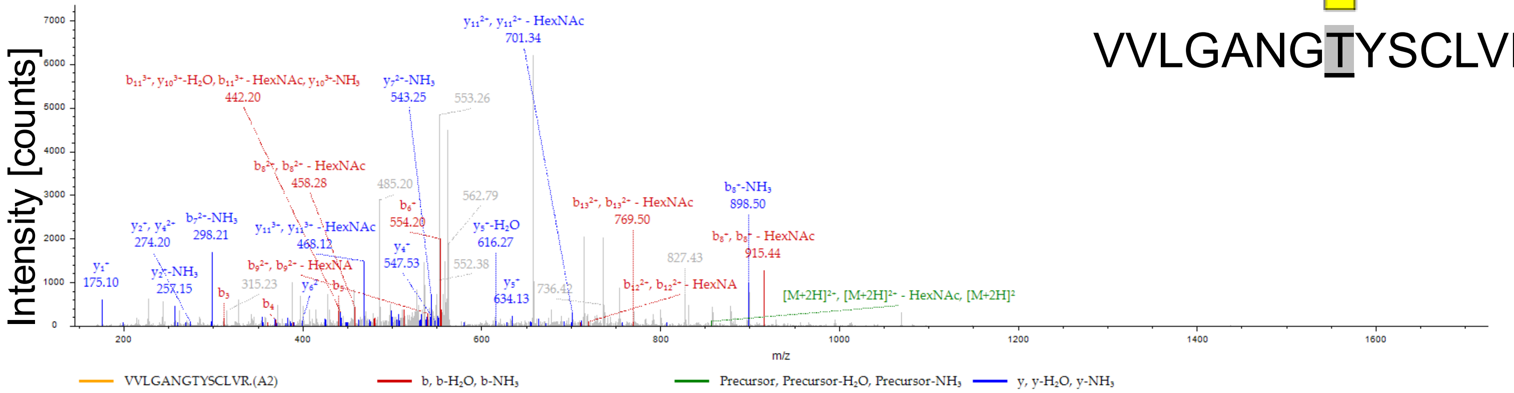


VVLGANGTYSCLVR

| #1 | b+ | b2+ | b3+ | Seq. | y+ | y2+ | y3+ | #2 |
|---|---|---|---|---|---|---|---|---|
| 1 | 100.07569 | 50.54148 | 34.03008 | V | | | | 14 |
| 2 | 199.14410 | 100.07569 | 67.05289 | V | 1612.79993 | 806.90360 | 538.27150 | 13 |
| 3 | 312.22817 | 156.61772 | 104.74757 | L | 1513.73152 | 757.36940 | 505.24869 | 12 |
| 4 | 369.24963 | 185.12845 | 123.75473 | G | 1400.64746 | 700.82737 | 467.55400 | 11 |
| 5 | 440.28675 | 220.64701 | 147.43377 | A | 1343.62599 | 672.31663 | 448.54685 | 10 |
| 6 | 554.32967 | 277.66847 | 185.44808 | N | 1272.58888 | 636.79808 | 424.86781 | 9 |
| 7 | 611.35114 | 306.17921 | 204.45523 | G | 1158.54595 | 579.77661 | 386.85350 | 8 |
| 8 | 915.47819 | 458.24273 | 305.83091 | T-HexNAc | 1101.52449 | 551.26588 | 367.84635 | 7 |
| 9 | 1078.54152 | 539.77440 | 360.18536 | Y | 797.39744 | 399.20236 | 266.47066 | 6 |
| 10 | 1165.57355 | 583.29041 | 389.19603 | S | 634.33411 | 317.67069 | 212.11622 | 5 |
| 11 | 1325.60419 | 663.30574 | 442.53958 | -Carbamidometh | 547.30208 | 274.15468 | 183.10554 | 4 |
| 12 | 1438.68826 | 719.84777 | 480.23427 | L | 387.27143 | 194.13935 | 129.76199 | 3 |
| 13 | 1537.75667 | 769.38197 | 513.25707 | V | 274.18737 | 137.59732 | 92.06731 | 2 |
| 14 | | | | R | 175.11895 | 88.06311 | 59.04450 | 1 |

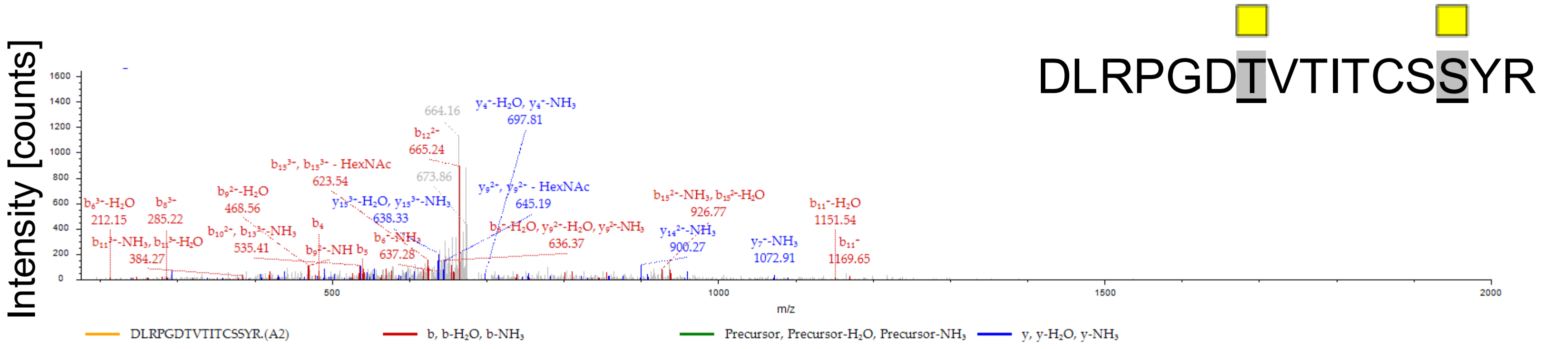


| #1 | b⁺ | b²⁺ | b³⁺ | Seq. | y⁺ | y²⁺ | y³⁺ | #2 |
|---|---|---|---|---|---|---|---|---|
| 1 | 116.03422 | 58.52075 | 39.34959 | D | | | | 16 |
| 2 | 229.11828 | 115.06278 | 77.04428 | L | 2132.01759 | 1066.51243 | 711.34405 | 15 |
| 3 | 385.21939 | 193.11334 | 129.07798 | R | 2018.93353 | 1009.97040 | 673.64936 | 14 |
| 4 | 482.27216 | 241.63972 | 161.42890 | P | 1862.83242 | 931.91985 | 621.61566 | 13 |
| 5 | 539.29362 | 270.15045 | 180.43606 | G | 1765.77965 | 883.39346 | 589.26473 | 12 |
| 6 | 654.32056 | 327.66392 | 218.77837 | D | 1708.75819 | 854.88273 | 570.25758 | 11 |
| 7 | 958.44762 | 479.72745 | 320.15406 | T-HexNAc | 1593.73124 | 797.36926 | 531.91527 | 10 |
| 8 | 1057.51603 | 529.26165 | 353.17686 | V | 1289.60419 | 645.30574 | 430.53958 | 9 |
| 9 | 1158.56371 | 579.78549 | 386.85942 | T | 1190.53578 | 595.77153 | 397.51678 | 8 |
| 10 | 1271.64777 | 636.32752 | 424.55411 | I | 1089.48810 | 545.24769 | 363.83422 | 7 |
| 11 | 1372.69545 | 686.85136 | 458.23667 | T | 976.40404 | 488.70566 | 326.13953 | 6 |
| 12 | 1532.72610 | 766.86669 | 511.58022 | -Carbamidometh | 875.35636 | 438.18182 | 292.45697 | 5 |
| 13 | 1619.75813 | 810.38270 | 540.59089 | S | 715.32571 | 358.16649 | 239.11342 | 4 |
| 14 | 1909.86953 | 955.43840 | 637.29469 | S-HexNAc | 628.29368 | 314.65048 | 210.10275 | 3 |
| 15 | 2072.93286 | 1036.97007 | 691.64914 | Y | 338.18228 | 169.59478 | 113.39894 | 2 |
| 16 | | | | R | 175.11895 | 88.06311 | 59.04450 | 1 |
